# Transitions from monotonic to tuned responses in recurrent neural network models during timing prediction

**DOI:** 10.1101/2024.08.29.610320

**Authors:** Evi Hendrikx, Daniel Manns, Nathan van der Stoep, Alberto Testolin, Marco Zorzi, Ben M. Harvey

## Abstract

The brain exhibits a gradual transition in responses to visual event duration and frequency through the visual processing hierarchy: from monotonically increasing to timing-tuned responses. Over their hierarchies, properties of both response types are progressively transformed. Here, we implement simulations based on artificial neural networks to investigate the requirements of neural systems for the emergence of such responses and their properties’ transformations. We see that recurrent networks develop monotonic responses whose properties’ progressions over network layers resemble those over brain areas. Responses to another sensory quantity, Furthermore, recurrent networks can further develop tuned responses, but only with training, a gradual transition between monotonic and tuned responses emerges. Particularly, if this training is done on predictable sequences, the tuned properties’ progressions resemble those observed in the brain. These results suggest that the emergence of visual timing-tuned responses and the subsequent hierarchical transformations of these responses result from recurrent neural computation and predictive processing of sensory event timing.

## Introduction

Perceiving the durations and frequency of sensory events is vital to understanding our dynamic environment. In the brain, early stages of visual processing show neural responses that monotonically increase in response amplitude with event duration and frequency, reflecting the inherent dynamics of sensory processing (Hendrikx et al., 2022; Stigliani et al., 2017; Zhou et al., 2018). Tuned responses peaking for specific durations and frequencies gradually emerge through the visual processing hierarchy, into temporal-occipital, parietal and frontal areas. This gradual emergence suggests that sensory timing representations may straightforwardly be derived from early sensory responses, rather than from dedicated processes responding to passing time. Once timing-tuned responses have emerged, the properties of these response functions progressively change in a hierarchy of timing maps. However, the computational mechanisms underlying these hierarchical changes of responses to event timing remain unclear.

Responses to another sensory quantity, visual numerosity, show similar progressions. As with timing, early visual responses monotonically increase with numerosity (DeWind et al., 2019; Park et al., 2016; Paul et al., 2022), while tuned responses to specific numerosities are found in temporal-occipital, parietal and frontal areas (Harvey et al., 2013; Harvey & Dumoulin, 2017; Nieder et al., 2002; Nieder & Miller, 2004). Computational studies reveal both monotonic and tuned responses to numerosity in artificial neural networks trained on a numerosity generation task (Stoianov & Zorzi, 2012; Zorzi & Testolin, 2018). The distribution of tuned responses changes with training to eventually match that seen in monkey parietal neurons (Viswanathan & Nieder, 2013; Zorzi & Testolin, 2018). Even deep convolutional networks with no training (i.e., random weights) show monotonically increasing responses to numerosity in earlier layers and numerosity-tuned responses in later layers (Kim et al., 2021), suggesting that this neural network architecture alone is sufficient to produce responses like those seen in the brain. Could timing-tuned responses and their subsequent transformations similarly reflect the basic architecture of the brain’s neural networks?

Numerosity and timing processing require quite different ways of integrating neural responses. For spatial quantities like numerosity, spatial integration of image features can be straightforwardly achieved by neurons with a spatial spread of their inputs, as in convolutional neural network models (Lecun et al., 1990). Conversely, integration of temporal responses (to event onsets, offsets and sustained responses to ongoing events) requires neurons to have information about past events. Recurrent connections in neural networks make the nodes’ current activity dependent on both ongoing input and their past activity, and therefore allow such temporal integration (Elman, 1990; Jordan, 1997).

In the current work, we therefore asked whether monotonic and tuned responses to event timing, and the progressions of their response properties seen in the brain, could spontaneously emerge in multi-layer recurrent neural networks. We hypothesized that the combination of recurrent processing, between-layer integration of the resulting responses and training to predict upcoming inputs in predictable sequences could support the development of timing-tuned responses. This predicts that recurrent networks with two or more hidden layers would show more tuned responses in their final layers that networks with one hidden layer and have superior performance in this input prediction task. Furthermore, we hypothesized that recurrent and between layer interactions over multiple processing stages may result in the gradual transition from monotonic to tuned responses and the gradual transformations of properties of both types of responses that we have observed in the brain.

## Results

### Recurrent networks can predict upcoming events in repetitive sequences

We created two sets of artificial neural networks with one to five hidden layers with independent recurrent nodes (each node connected only to its own state on the previous time step; Li et al., 2018) and full connections between layers (IndRNNs). We chose this neural architecture over a recurrent neural network with recurrent connections to all nodes in each layer, because it makes each nodes’ responses easier to assess and it does not allow more complexity than a biological system (see Methods). In our ‘parameter-matched’ networks, the number of nodes in each layer was chosen to keep the total number of connection weights and biases as similar as possible between network depths, thereby separating effects of network depth and network complexity. However, this resulted in very few nodes in each layer of deeper networks. In our ‘layer-size-matched’ networks, the number of nodes in each layer was constant (at 16) regardless of network depth, so deeper networks were more complex. We also created five-layer networks without recurrent connections, which matched the number of connection weights and biases in the parameter-matched networks. For each of these network architectures we created 50 network repetitions each initialized with different random weights and trained these separately, allowing us to make statistical comparisons between architectures. We also investigated the responses of the same networks before training, separating the effects of network architecture and learning mechanisms.

We trained these networks to predict the states of upcoming time steps in a time series consisting of repetitive events (henceforth: input sequence). This is a form of self-supervised learning, as the network was not trained to explicitly label the timing of the events in the input sequence (Hinton, 2007; Testolin & Zorzi, 2016), but rather to generate supervisory signals based on the difference between its predicted and actual states of future time steps in the sequence. Events consisted of consecutive “on” and consecutive “off” states. Different input sequences differed in event duration (consecutive time steps of an input state), event period (time steps between the onsets of consecutive events, i.e. the sum of the time steps in the “on” duration and “off” duration), and phase (the time step of the first event onset) of the events (Fig. 1, top rows). We also trained the five-layer networks on shuffled versions of these input sequences, which lacked predictable timing but allow achieving some accuracy by predicting the most common input state.

**Fig. 1:**
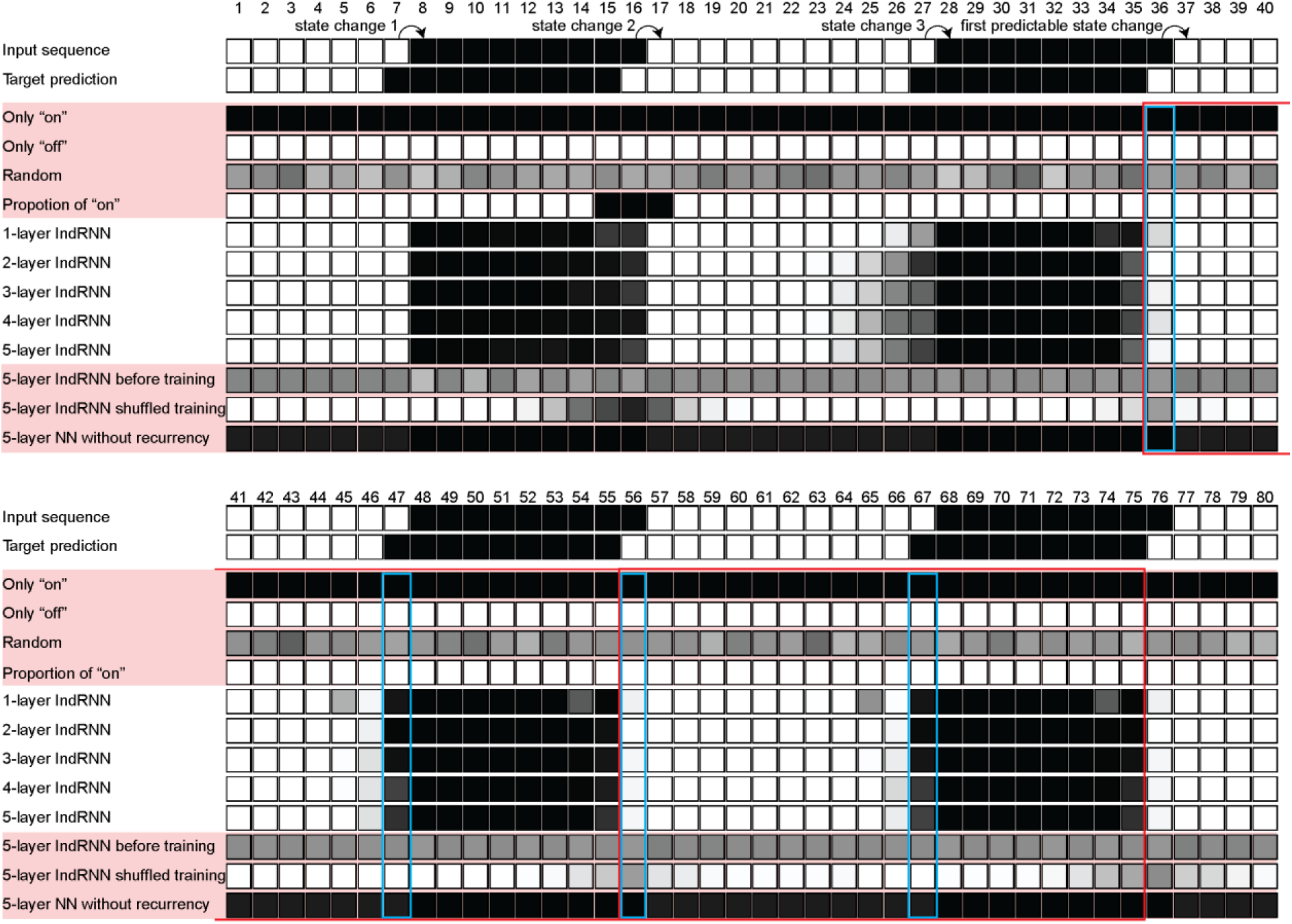
An example input sequence, target predictions, outcomes of possible strategies, and responses of the networks. For the input sequence and the target predictions a black square represents the “on” state and white represents the “off” state at each time step (numbers). This input sequence has an event “on” duration of 9 time steps, an event period of 20 time steps, and a phase shift of +8 time steps. Arrows indicate the location of state changes needed to establish the sequence’s predictability. Time steps used for per-event accuracy assessments are marked by red rectangles. Time steps used for per-state-change accuracy assessments are marked by blue rectangles. For the outcomes of possible strategies, and responses of the networks, the grayscale of the square indicates the proportion of network repetitions predicting an “on” output state.

All networks improved their prediction performance with additional training epochs, eventually reaching convergence (Fig. S1). For the parameter-matched networks, the final training loss of the one-layer network converged to a significantly higher loss than the other networks, while the two-layer network’s training loss was significantly lower than that of the four- and five-layer networks (*H*_(4)_=45.71, *p*=3×10^-9^, *ε*^2^=0.17; Table S1A). Networks without recurrent connections and networks trained on shuffled input sequences both exhibited far higher training loss than all recurrent networks (Fig. S1; Table S1B). Layer-size-matched networks revealed that increased complexity led to decreased training loss (*H*_(4)_=148.38, *p*=5×10^-31^, *ε*^2^=0.58; Fig. S1; Table S1C).

Fig. 1 outlines the task’s requirements, some hypothetical strategies and some networks’ outputs for an example input sequence. A network that always predicts its current input, “on” time steps, “off” time steps, or random values should not improve its performance during processing of the sequence. A network that simply represents the proportion of “on” time steps can at best predict the most common input state; its accuracy should improve throughout the sequence, but remain low when the proportion of on and off states are similar (as in Fig. 1’s example). However, a network with an internal representation of the input sequence’s timing can predict the next time step specifically after three state changes (from “on” to “off” or from “off” to “on”) in the sequence. Only at this time step has the network processed one whole event input. Therefore, we assessed accuracy in predicting whole events after this this third state change (Fig. 1).

After training, the predictions made by trained IndRNNs were consistent with an internal representation of the input sequences’ timing (Fig. 1). These networks initially predict similar values to their input. After three state changes, the networks’ predictions matched the target state rather than the network’s current input.

The predictions of IndRNNs before training mimicked the hypothesized pattern for random predictions. Those of a neural network without recurrency mimicked the hypothesized pattern of “on”-only predictions. Those of a network trained on shuffled data mimicked the hypothesized pattern for a network representing the proportion of “on” time steps, predicting the most common state seen before each time step. Therefore, recurrency, training, and temporal structure in the training sequences were all necessary for the network to develop an internal representation of event timing.

### Maximum accuracy improved with a second hidden layer

We kept 20% of input sequences as a test set to evaluate each network’s performance in predicting times steps in unseen input sequences after training was complete. Time steps where the upcoming input state differed from the current input state particularly require an accurate event timing representation, but most upcoming time steps simply repeat the current input state. We therefore assessed accuracy first as the average accuracy across all of an event’s time steps (per-event accuracy) and second as the average accuracy across the time steps where the input sequence changes state (per-state-change accuracy).

The mean prediction accuracy differed between network depths (parameter-matched: per-event: *H*_(4)_=38.73, *p*=8×10^-8^, *ε*^2^=0.14; per-state-change: *H*_(4)_=24.12, *p*=8×10^-5^, *ε*^2^=0.08; Fig. 2A, Table S2A). For the parameter-matched networks, the one-layer networks gave lower per-event accuracies than the other networks and lower per-state-change accuracies than the two-, three-, and five-layer networks. However, accuracy did not significantly improve with additional layers (beyond two) and indeed the two-layer networks gave a higher per-event accuracy than the four-layer networks.

**Fig. 2:**
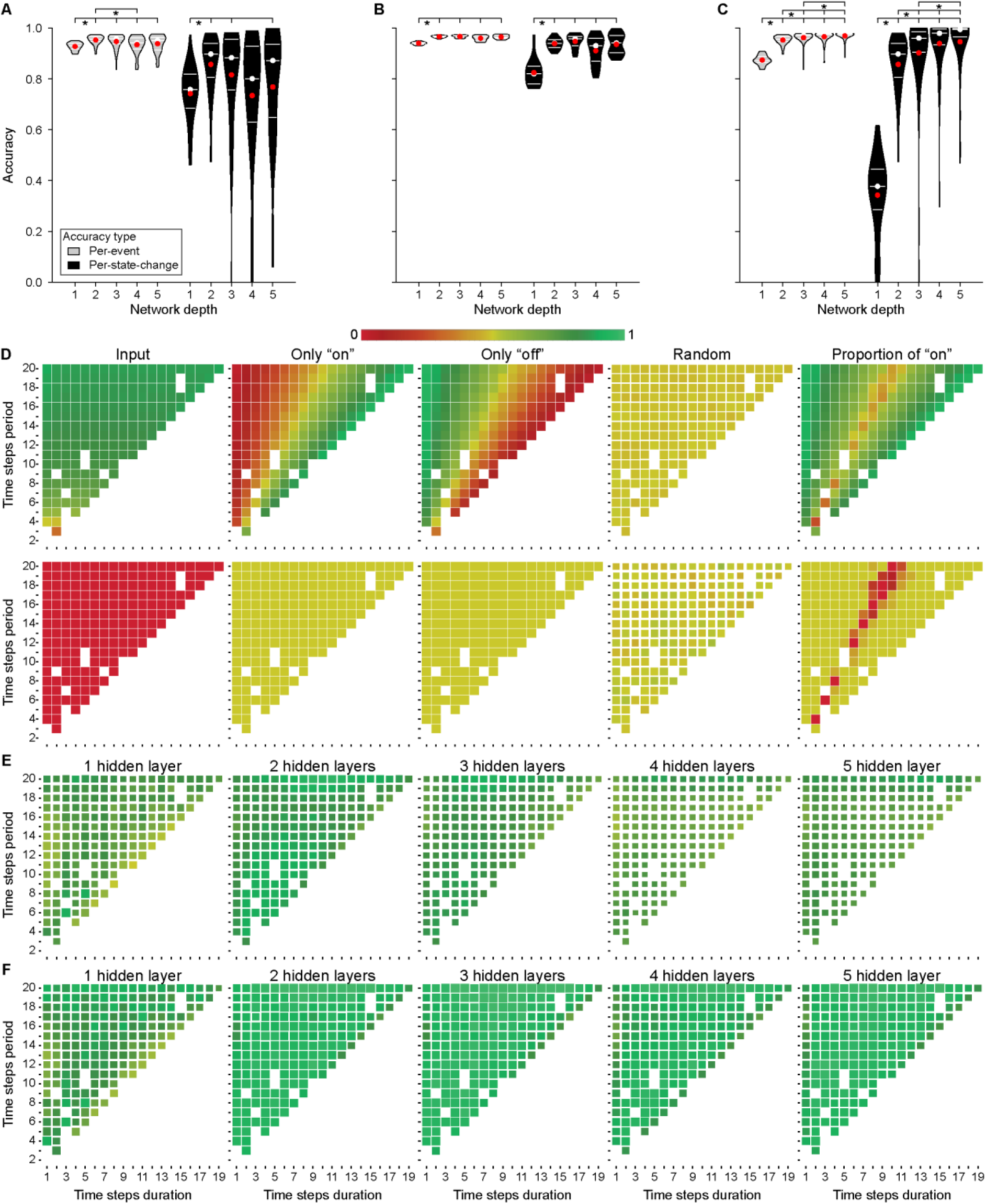
Best target prediction accuracies were greater and more consistent for recurrent networks with at least two hidden layers and did not resemble predictions of hypothetical strategies. (**A**) Per-event and per-state-change accuracies for various network depths for all 50 network repetitions of the parameter-matched networks. (**B**) Accuracies for the top 25 network repetitions of the parameter-matched networks. (**C**) Accuracies for all 50 network repetitions of the layer-size-matched networks. White lines and circles indicate interquartile ranges and medians, respectively. Red circles indicate the mean. All violins’ maximum widths are kept the same. Brackets and stars show significant differences in multiple comparisons between all network depths: all brackets to the left of the star are significantly different from all brackets to the right of the star (two-sided Dunn’s test, corrected for multiple comparisons using Šidák-Holm, *p* < 0.05) **(D)** The per-event (top) and per-state-change (bottom) accuracies for the hypothetical strategies shown in Fig. 1. **(E)** Per-state-change accuracies per timing for all 50 repetitions of the parameter-matched IndRNNs (per-event in Fig. S3A). **(F)** Per-state-change accuracies per timing for most accurate 25 repetitions of the parameter-matched IndRNNs (per-event in Fig. S3B). Colors indicate the mean accuracy over repetitions and presentations of each timing in the test set. The test set contained 20% of the input sequences, so omitted some timings. The sizes of the squares scale with standard deviation of accuracy between network repetitions: more variable outcomes between repetitions are shown as smaller squares

The parameter-matched networks with more layers had fewer nodes in each layer, which led to more variable accuracy in different repetitions with different random initializations (per-event: *W=*5.35, *p=*0.0004; per-state-change: *W=*4.57, *p*=0.001; Table S3; Fig. 2A). This variability could mask increases in maximum accuracy in deeper networks. We therefore assessed whether the networks’ maximum possible accuracy benefits from more layers by comparing the 25 (of 50) best-performing network repetitions for each network depth (Fig. 2B). Here, the one-layer networks gave a higher accuracy than all other depths (per-event: *H*_(4)_*=*61.03, *p=*2×10^-12^, *ε*^2^=0.23; per-state-change: *H*_(4)_=49.84, *p*=4×10^-10^, *ε*^2^=0.18; Table S2B), but there were no significant accuracy differences between other network depths.

In the layer-size-matched networks, where the network complexity increased with depth, prediction accuracy increased with network depth (per-event: *H*_(4)_=147.29, *p*=8×10^-31^, *ε*^2^=0.58; per-state-change: *H*_(4)_=147.47, *p*=7×10^-31^, *ε*^2^=0.58; Fig. 2C; Table S2C). These networks lacked the issue of decreasing nodes in deeper networks and as such different network depths lacked significant differences in the variance of their accuracy (per-event: *W=*0.83, *p=*0.508; per-state-change: *W=*2.21, *p*=0.068).

The median accuracy of the neural networks without recurrency, IndRNNs before training, and the IndRNNs trained on shuffled data were significantly lower than those of the comparable five-layer IndRNN (Fig. S2, Table S4).

Overall, multi-layer networks generally gave greater accuracy than one-layer networks. While more complex networks with more parameters also gave greater accuracy than simpler networks, increasing the number of layers above two without also increasing network complexity did not further increase accuracy. Furthermore, this accurate prediction required recurrency, training and predictable temporal structure in the input sequences.

### Two-layer networks allow high prediction accuracy at all event timings

Next, we asked whether deeper parameter-matched networks achieved more consistent performance across event timing. Fig. 2D shows how accurately the hypothetical strategies shown in Fig. 1 predicted targets for each event timing. Although lacking any predictive process, these strategies gave a high per-event accuracy for specific event timings. Nevertheless, their per-state-change accuracy was low for all timings, demonstrating this accuracy measure is more sensitive to predictive power. Again, networks without recurrency, IndRNNs without any training, and the IndRNNs trained on shuffled data mimicked the accuracy patterns in these hypothetical situations (Fig. S3D).

The one-layer networks had a higher variance in per-state-change accuracy across timings than the two-layer networks, but not other network depths (*H*_(4)_=14.44, *p=*0.006, *ε*^2^=0.04; Table S5A). Again, this lack of significant differences between one-layer networks and three-, four- or five-layer networks likely reflected the greater variability among repetitions of deeper parameter-matched networks. Indeed, among the most accurate 25 repetitions, one-layer networks had a higher variance in per-state-change accuracy across timings than all other networks (*H*_(4)_=39.48, *p=*6×10^-8^, *ε*^2^=0.14; Table S5B). Therefore, networks with at least two layers allow more consistent target prediction accuracy across event timings.

### Transitions from monotonic to tuned responses in successive IndRNN layers

Human brain recordings show different responses to event timing that either: (1) monotonically increase with event duration and frequency (following monotonic response functions); (2) peak for a particular combination of event duration and period (following tuned response functions); or (3) more ambiguously show characteristics of both response functions (mixed response functions). Monotonic responses predominate in early visual areas (Hendrikx et al., 2022; Stigliani et al., 2017; Zhou et al., 2018), while ambiguously and then clearly tuned responses gradually emerge through the visual hierarchy (Hendrikx et al., 2022). Do recurrent networks trained for timing prediction likewise show a transition from monotonic to tuned responses?

For each network node we analyzed the per-event activation for each timing, averaged across input sequences with the same duration and period but different phases. We fit the free parameters of both monotonic and tuned response functions on one half of the input sequences (Fig. 3, top) and used the complementary half (Fig. 3, second row) to evaluate the resulting functions’ predictions (Fig. 3, bottom rows). This determined which response function best captured each node’s response. Where tuned response functions predicted the response better but the peak of the response function was outside the range of timings used, these response functions had properties of both monotonic and tuned functions so were categorized as mixed responses.

**Fig. 3:**
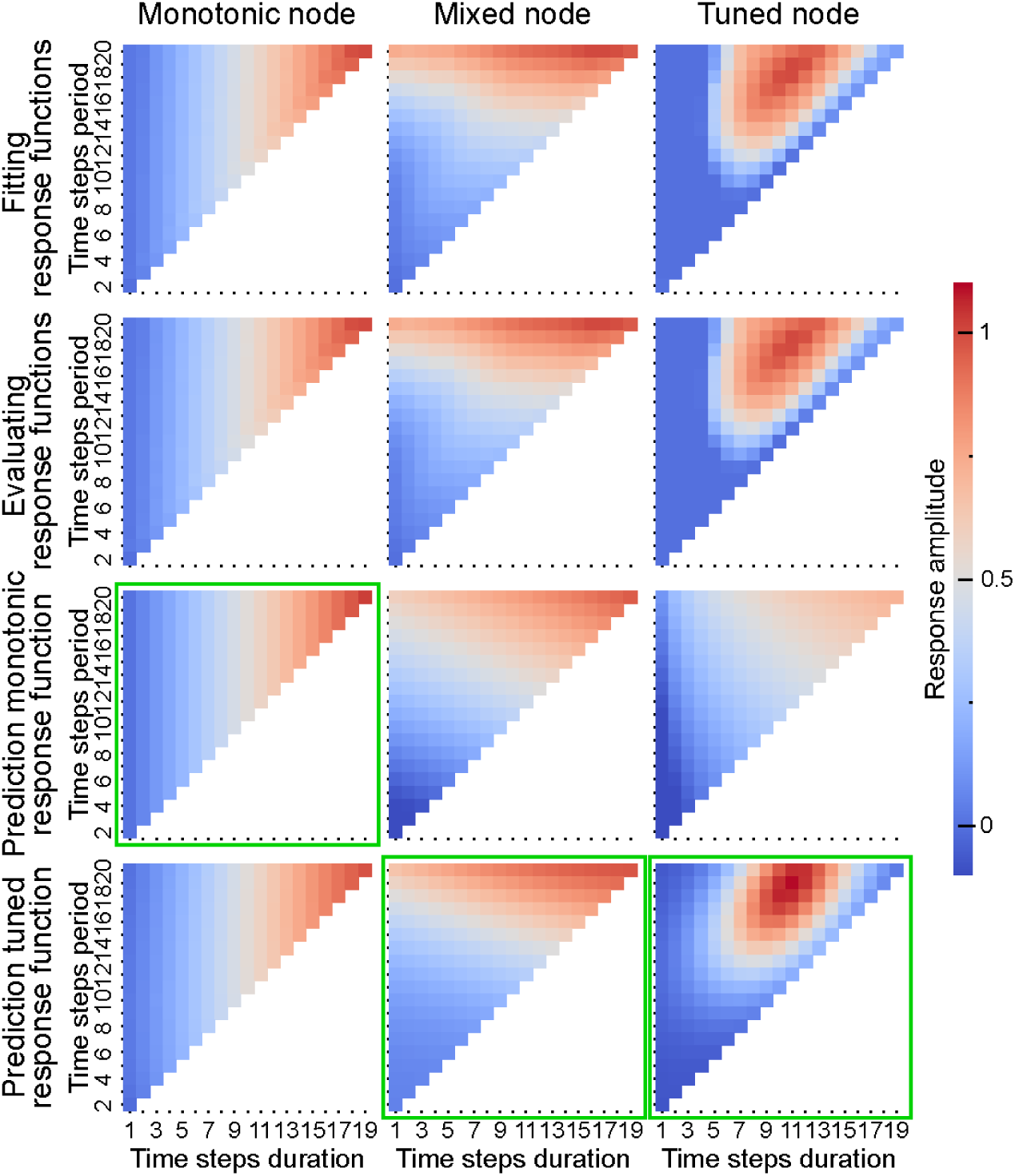
Responses and fit response functions for example nodes with diverse responses. The top row shows the node’s response for each timing (colors) in the half of input sequences used for response functions fitting. The second row shows the node’s responses in the complementary half of input sequences used for response function evaluation. The bottom rows show the fit monotonic and tuned response functions for each node, with the best evaluated response function marked in green. The left, middle and right columns follow monotonic, mixed and tuned nodes, respectively.

For all multi-layer networks, the proportion of monotonic responses differed significantly between layers (Figs. 4A & S4; Table S6A). Post hoc comparisons (Tables S6B-C) generally show significant steady decreases in the proportion of monotonic responses through the networks’ successive layers. The proportion of mixed and tuned responses likewise differed between layers (Table S6A). In contrast to the monotonic responses, these proportions generally showed significant steady increases through each network’s successive layers.

**Fig. 4:**
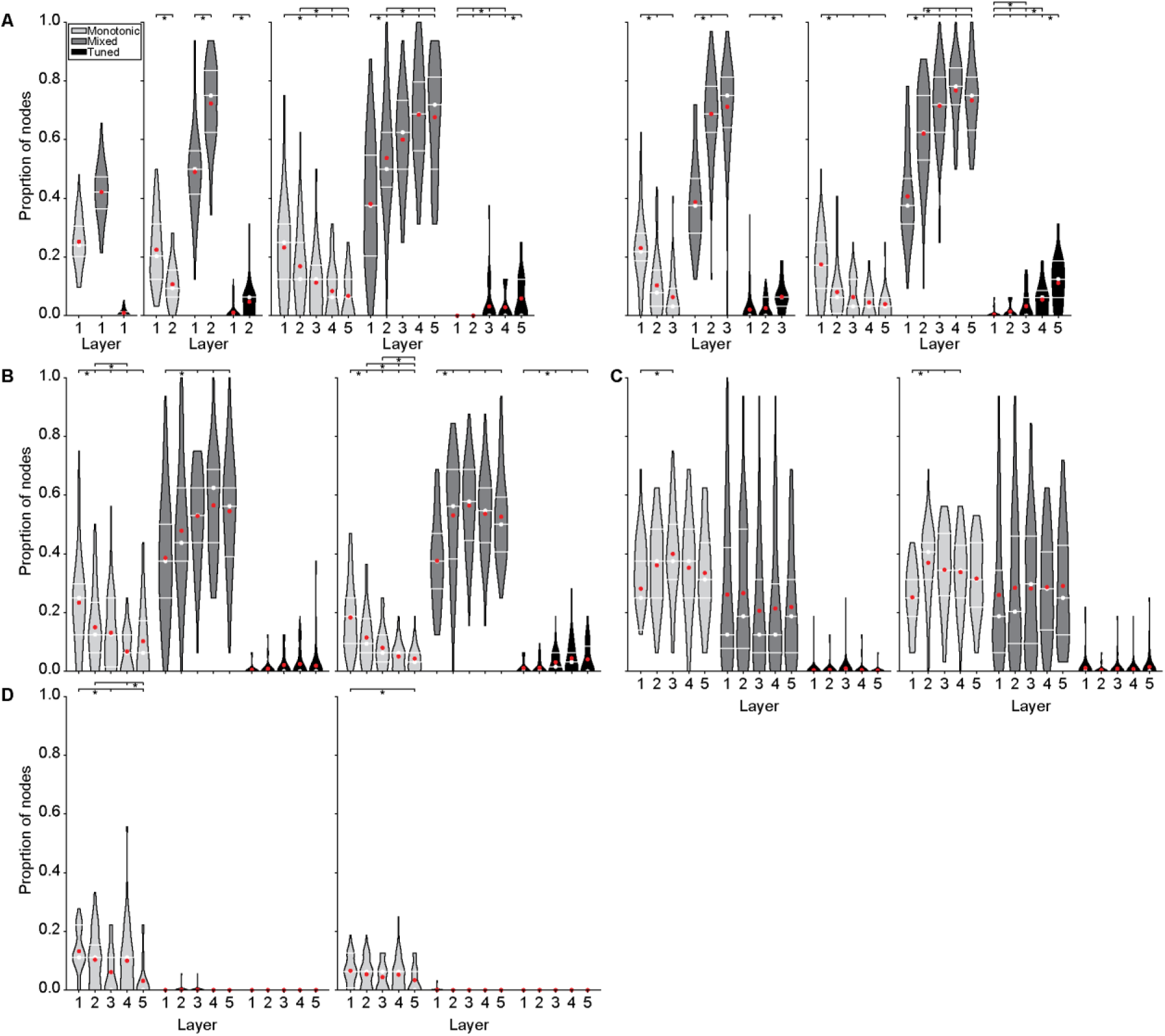
Progressive transitions from monotonic to mixed and tuned responses across network layers in trained IndRNNs. (**A**) Proportions of nodes classified as monotonic, mixed, or tuned in parameter-matched IndRNNs of 1, 2 and 5 layers (left) or layer-size-matched IndRNNs of 3 and 5 layers (right). See Fig. S4A for other depths. (**B**) As for the five-layer network in (A) for IndRNNs trained on shuffled data. (**C**) As for the five-layer network in (A) for IndRNNs before training. (**D**) As for the five-layer network in (A) for NNs without recurrency. Format follows Fig. 2A.

Five-layer IndRNNs trained on shuffled data showed similar transitions from monotonic to mixed and tuned responses, though less clearly (Fig. 4B; Tables S6A & S6D-E). However, five-layer IndRNNs before training showed all types of responses and some significant differences in the proportion of monotonic responses (Fig. 4C; Tables S6A & S6F-G), but no clear increase or decrease over layers. In contrast to all trained IndRNNs, this untrained network also showed (in all layers) the highest proportion of monotonic responses, fewer mixed responses and very few tuned responses. Five-layer neural networks without recurrency showed no response to any timing in most nodes. Their few responsive nodes again exhibited a higher proportion of monotonic responses in early layers (Fig. 4D; Table S6A & S6H-I), but extremely few mixed responses and no tuned responses in any layer.

Therefore, tuned responses progressively emerged through the layers of trained IndRNNs, while monotonic responses progressively decreased. These progressions were clearer when trained on predictable input sequences and absent before training or without recurrency. No tuned responses (and very few mixed responses) were found without recurrency.

### Proportions of tuned responses depend on network complexity and between-layer integration

Both parameter-matched and layer-size-matched IndRNNs showed a progressive emergence of tuned responses and reduction of monotonic responses across network layers. Do successive layers of deeper IndRNNs eventually allow more tuned and mixed responses to emerge, or simply allow a more gradual transition to similar responses in their final layers? Comparing the proportions of monotonic, mixed and tuned responses in the final layers of parameter-matched and layer-size-matched IndRNNs with different depths separates effects of depth and complexity: Parameter-matched networks are similarly complex regardless of depth while layer-size-matched networks increase in complexity with depth.

In parameter-matched IndRNNs, the proportion of monotonic, mixed and tuned responses in last layer of networks from two to five layers showed no significant differences (Fig. 5A, Table S7A). Only the last (and only) layer of one-layer networks had significantly more monotonic responses and fewer mixed responses than the last layer of all multi-layer networks. For tuned responses, no differences between network depths reached significance.

**Fig. 5:**
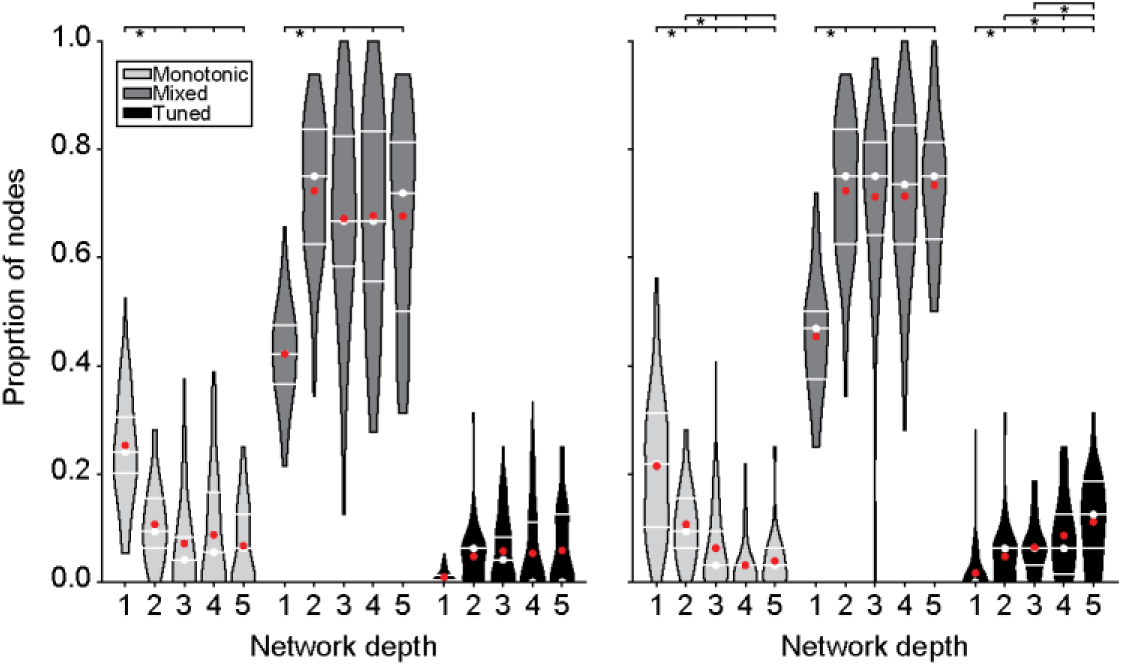
Decrease in the proportion of final-layer monotonic responses and increase in the proportion of final-layer tuned responses follows network complexity in multi-layer networks. (**A**) Parameter-matched IndRNNs from two to five layers do not significantly differ in the proportions of monotonic, mixed or tuned nodes, while one-layer IndRNNs have more monotonic and fewer mixed nodes than these multi-layer IndRNNs. (**B**) In layer-size-matched IndRNNs, where network complexity increases with depth, more complex five-layer networks show fewer monotonic nodes and more tuned nodes than simpler two-layer networks. Format follows Fig. 2A.

Conversely, in layer-size-matched IndRNNs from two to five layers, the proportions of monotonic, mixed, and tuned nodes generally differed between the last layers for many depths (Fig. 5B, Table S7B). The proportion of monotonic nodes decreased from two- to five-layer networks, while the proportions of tuned nodes increased. The proportion of mixed responses did not change from two- to five-layer networks. As in parameter-matched networks, one-layer networks again showed more monotonic responses and fewer mixed and tuned responses than multi-layer networks.

Therefore, the pattern of responses in the final layer reflects the network’s complexity rather than the number of layers. One-layer networks allow more limited computations as they lack between-layer integration across nodes. Multi-layer parameter-matched networks always allow a similar set of computations and so similar final response properties. Only more complex (not deeper) networks allow further computations. These differences in the proportions of response types in each networks’ final layers mirror the differences in maximum task accuracy seen in Figures 2B-C.

### Monotonic parameters of IndRNNs follow similar progressions to the brain, regardless of training

Next, we asked how parameters of the monotonic and tuned response functions differed between network layers, and whether these differences resembled differences in response function parameters between human brain areas. Here we used layer-sized-matched IndRNNs because these showed at least some tuned nodes in each layer, and their five-layer versions to reveal gradual progressions between layers more clearly. Nevertheless, other depths generally showed similar patterns, although sometimes less clearly (Fig. S5). To investigate how the progressions depended on training, we repeated these analyses in the same network architecture before training and that trained on shuffled input sequences. We also repeated this for networks without recurrent connections, but limited this to analysis of monotonic response function parameters because these networks showed no tuned responses.

For the response functions of monotonic nodes, Kruskal-Wallis tests revealed significant differences between layers in the exponents on duration (Fig. 6A) for IndRNNs trained on repetitive events (*H*_(4)_=74.79, *p*=2×10^-15^, *ε*^2^=0.11), and also IndRNNs before training (*H*_(4)_=196.55, *p*=2×10^-41^, *ε*^2^=0.07) and those trained on shuffled data (*H*_(4)_=98.69, *p*=2×10^-20^, *ε*^2=^0.13). Generally, this exponent decreases from layers one to two in all of these IndRNNs, then does not differ significantly from layers two to five (Tables S8A-C). The network without recurrency also showed significant differences in this exponent between network layers (*H*_(4)_=22.76, *p*=0.0001, *ε*^2^=0.05, Table S8D), but unlike IndRNNs showed an increase over network layers. In the human brain, the exponent on duration of monotonic response functions differed between visual field maps (*H*_(9)_=52.29, *p*=2×10^-5^, *ε*^2^=0.10; Fig. 6B; Table S8E), generally decreasing from earlier to later maps as between IndRNN layers.

**Fig. 6:**
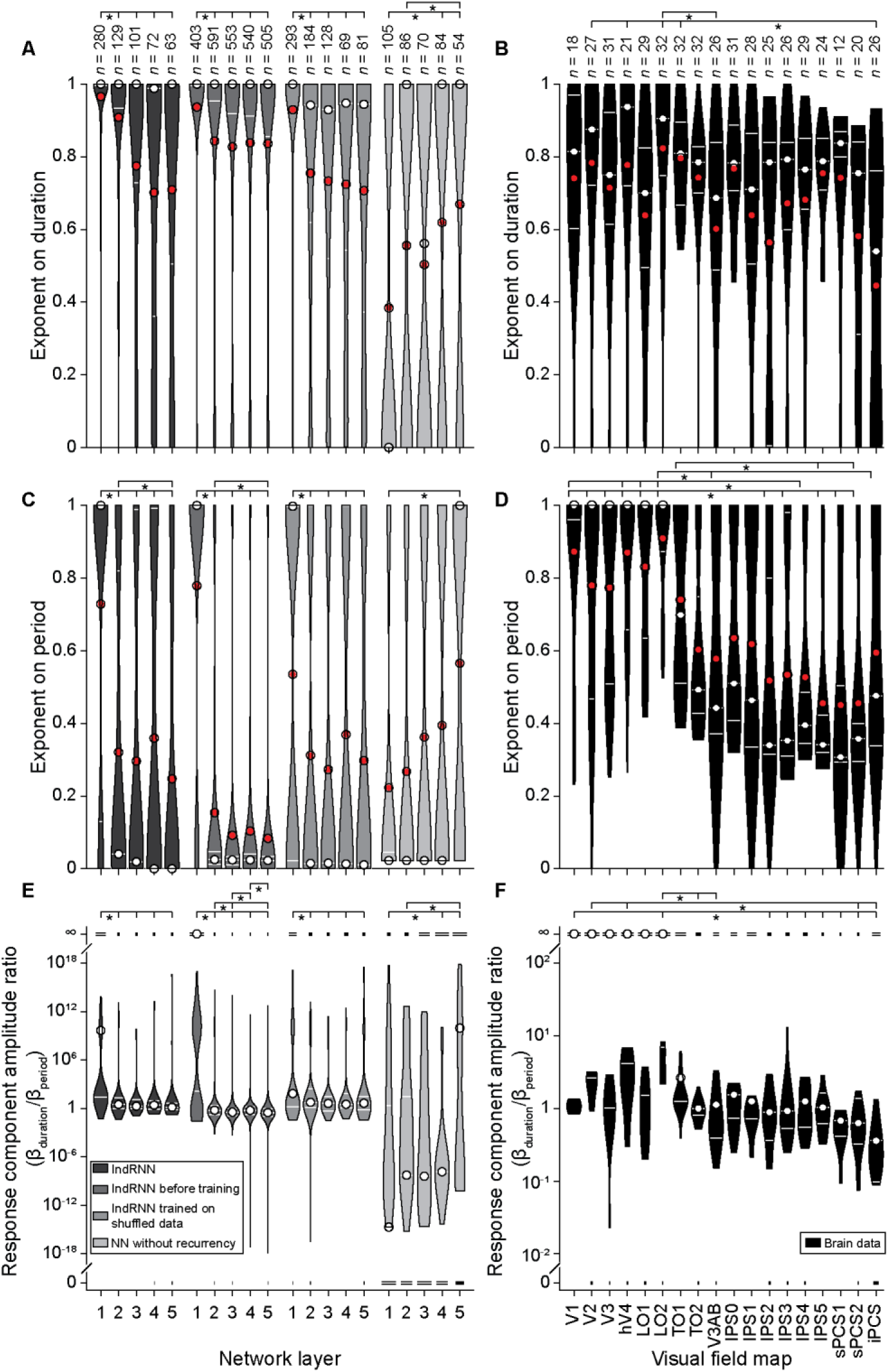
Monotonic parameter progressions in IndRNNs’ monotonic nodes resembled those in the human brain regardless of training regimen. (**A & C**) The exponents on duration (A) and period (C) decreased from the first layer to later layers in all five-layer IndRNNs, but not networks without recurrency. (**B & D**) These exponent similarly decreased through the visual field map hierarchy. (**E**) The ratio between the amplitudes of monotonic responses to duration and period also decrease from the first layer to later layers in all five-layer IndRNNs, but not networks without recurrency. (**F**) This ratio similarly decreased through the visual field map hierarchy. In (E) and (F), the lines at 0 and ∞ represent nodes with responses only driven by period and duration, respectively. The width of corresponding lines and maximum width of the corresponding violin scale with the number of nodes in their respective value range. The maximum width of all (whether of the line or the violin) are kept constant. *n* gives number of included nodes in each network layers, or the number of hemispheres per brain area. Brain data adapted from Hendrikx et al. (2022). Format follows Fig. 2A.

We similarly found significant differences between layers in the exponent on period (Fig. 6C) for IndRNNs trained on repetitive events (*H*_(4)_=152.83, *p*=5×10^-32^, *ε*^2^=0.23), and also IndRNNs before training (*H*_(4)_=511.58, *p*=2×10^-109^, *ε*^2^=0.20) and those trained on shuffled data (*H*_(4)_=69.17, *p*=3×10^-14^, *ε*^2^=0.09). Generally, this exponent decreases from layers one to two in all of these IndRNNs (Tables S8A-C), with a further decrease from layers two to five in IndRNNs trained on repetitive events and those before training. Again, the network without recurrency also showed significant differences in this exponent between network layers (*H*_(4)_=9.64, *p*=0.047, *ε*^2^=0.01, Table S8D), but unlike IndRNNs showed an increase over network layers. Again, in the human brain, the exponent on period of monotonic response functions differed between visual field maps (*H*_(9)_=122.94, *p*=4×10^-18^, *ε*^2^=0.25; Fig. 6D; Table S8E), generally decreasing from earlier to later maps as between IndRNN layers. Exponents on both duration and period were bimodally distributed in all networks (Table S9).

The ratio between the amplitudes of monotonic responses to duration and period also differed between network layers (Fig. 6E) for IndRNNs trained on repetitive events (*H*_(4)_=189.32, *p*=7×10^-40^, *ε*^2^=0.29), those before training (*H*_(4)_=530.04, *p*=2×10^-113^, *ε*^2^=0.20) and those trained on shuffled data (*H*_(4)_=36.85, *p*=2×10^-7^, *ε*^2^=0.04). In all these networks, this ratio switches from strongly favoring the duration component (high ratio) in the first layer to a more equal contribution of the two components (ratio around one) in later layers (Tables S8A-C). Again, the network without recurrency also showed significant differences in this exponent between network layers (*H*_(4)_=23.28, *p*=0.0001, *ε*^2^=0.05, Table S8D), but unlike IndRNNs showed ratios favoring the period component (below one) in the layers one to four to favoring the duration component (above one) in the last layer. Again, in the human brain this ratio differed between visual field maps (*H*_(9)_=90.08, *p*=6×10^-12^, *ε*^2^=0.18; Table S8E; Fig. 6F) generally decreasing from earlier to later maps as between IndRNN layers, though including less extreme ratios.

Therefore, all parameters of monotonic response functions progress through the layers on IndRNNs similarly to the progressions through the brain’s visual field maps regardless of training. Networks without recurrency lack these progressions. This suggests that recurrency, not training, allows the hierarchical transformations of monotonic responses seen through the brain’s visual hierarchy.

### Tuned parameter follow similar progressions to the brain only in IndRNNs trained on repeated events

For the nodes fit best by the tuned model, we only analyzed the parameters of the tuned nodes. Here we excluded mixed nodes because parameters including peak responses outside the used timing range cannot be determined accurately. Here we used the brain’s timing maps (Harvey et al., 2020) (rather than visual field maps) for comparison as these each contain neural populations with a full range of timing preferences. These timing maps show progressive changes in several properties of their tuned neural populations’ response functions in a hierarchy from posterior and inferior to anterior and superior timing maps.

The brain’s timing maps significantly differ in their average preferred durations (*H*_(9)_=28.41, *p*=0.0008, *ε*^2^=0.13; Fig. 7B) and periods (*H*_(9)_=30.46, *p*=0.0004, *ε*^2^=0.14; Fig. 7D), as do the tuned nodes in different layers of IndRNNs trained on repetitive sequences (Preferred duration (Fig. 7A): *H*_(4)_=19.79, *p*=6×10^-4^, *ε*^2^=0.05. Preferred period (Fig. 7C): *H*_(4)_=32.96, *p*=1×10^-6^, *ε*^2^=0.08). Preferred duration and period lack clear and systematic progressions from early to later brain areas, though the intermediate timing map TLS has slightly lower preferred durations and periods than the superior timing map (TPCS) (Table S8F). To an extent, tuned nodes of IndRNNs trained on repetitive sequences show similar differences, with intermediate layers (two and three) having significantly lower preferred durations and periods than later layers (Table S8A). Before training, the network nodes in each layer also showed differences in preferred duration (*H*_(4)_=28.91, *p*=8×10^-6^, *ε*^2^=0.34; Fig 7A) and period (*H*_(4)_=20.23, *p*=0.0005, *ε*^2^=0.22; Fig 7C). However, unlike in the trained IndRNNs and the brain, both preferences increase starting from the first layer (Table S8B). After training on shuffled data, there are no significant differences in duration (*H*_(4)_=1.81, *p*=0.771, *ε*^2^=-0.01; Fig 7A) or period (*H*_(4)_=2.01, *p*=0.735, *ε*^2^=-0.01; Fig 7C) preferences between layers.

**Fig. 7:**
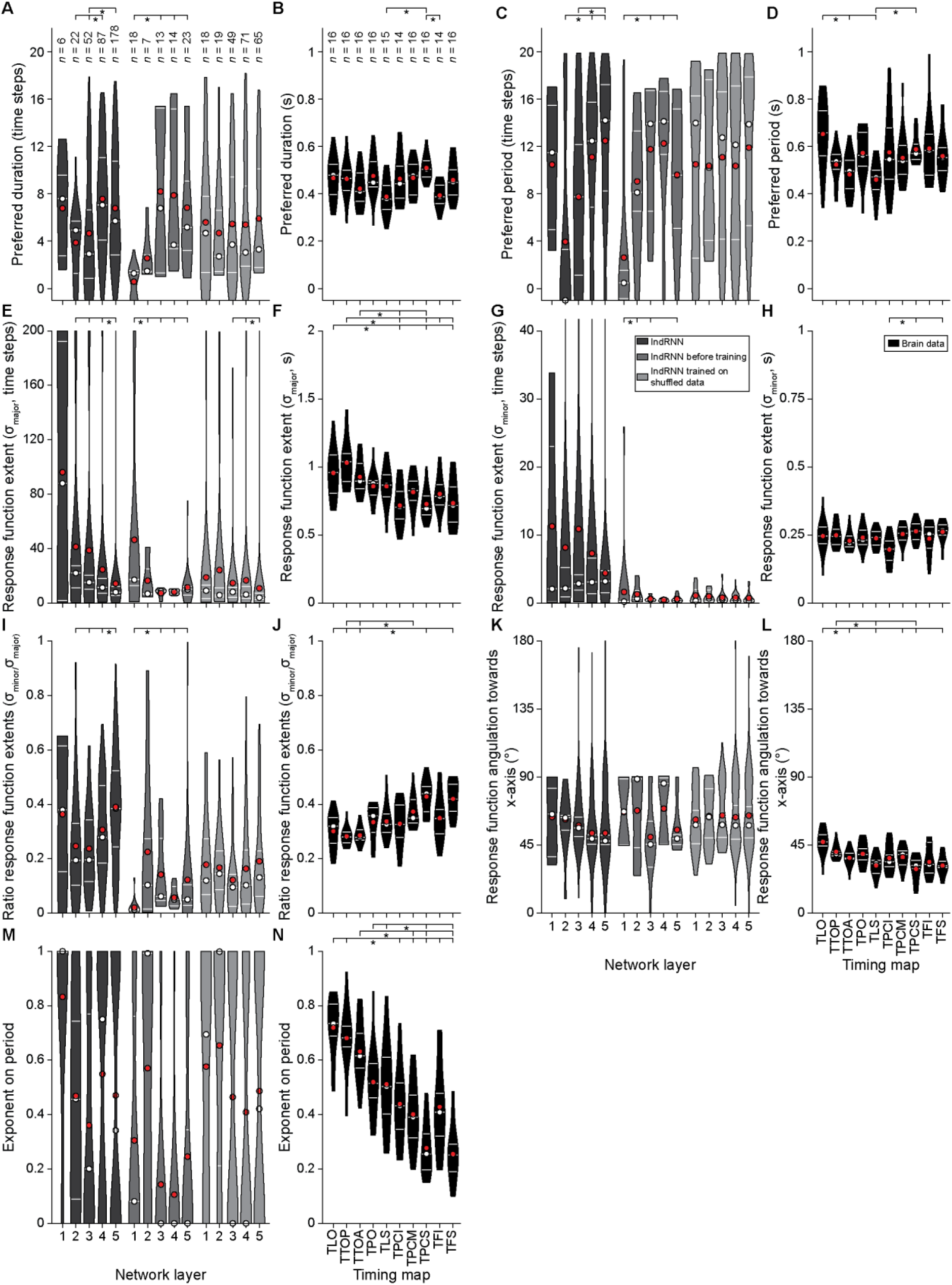
Tuned parameter progressions between layers of IndRNNs trained on repetitive input sequences generally follow progressions through the brain’s visual timing maps and differ between training regimens. (**A-D**) Preferred duration and period in IndRNNs and the brain lack clear and systematic progressions from early to later timing maps. (**E-F**) The response function’s major extent decreased from early to later network layers (E), regardless of training regimen, and also from early to later timing maps (F). (**G-H**) The response function’s minor extent decreased only in IndRNNs before training. (**I-J**) The ratio of the response function extents (the roundness of the response function) increased similarly to the brain’s timing map hierarchy only through the layers of IndRNNs trained on repetitive input sequences. (**K-L**) The angulation of the response function progressed similarly to the brain’s timing map hierarchy only through the layers of IndRNNs trained on repetitive input sequences. (**M-N**) The exponent on event period systematically decreased through the brain’s timing map hierarchy, but became increasingly bimodal through the layers only in IndRNNs trained on repetitive or shuffled input sequences. Brain data adapted from Harvey et al. (2020). Format follows Fig. 2A, except in (K) and (L) where red and white circles are circular means and median, and white lines show the 95% confidence interval of the circular median.

The extents of the response functions along their major axes differed considerably between IndRNN layers (*H*_(4)_=39.62, *p*=5×10^-8^, *ε*^2^=0.10; Fig.7E), decreasing from lower to higher network layers (Table S8A). However, the functions’ extent along their minor axes showed no change between layers (*H*_(4)_=1.63, *p*=0.804, *ε*^2^=-0.01; Fig. 7G). As the major extent differed while the minor extent was similar across layers, their ratio (the roundness of the response function) differed between layers (*H*_(4)_=32.18, *p*=2×10^-6^, *ε*^2^=0.08; Table S8A; Fig. 7I). This ratio increased (towards one) over layers, giving more rounded response functions in higher layers. The brain showed significance differences between timing maps in the response function’s major extent (*H*_(9)_=47.70, *p*=3×10^-7^, *ε*^2^=0.25; Fig. 7F), minor extent (*H*_(9)_=21.08, *p*=0.012, *ε*^2^=0.08; Fig. 7H) and their ratio (*H*_(9)_=60.92, *p*=9×10^-10^, *ε*^2^=0.34; Fig. 7J). As between neural network layers, these timing maps showed a hierarchical decrease in the response functions’ major extents, no systematic change in the minor extents and an increasingly rounded response function (Table S8F). Before training, the IndRNN showed significant differences between layers in the response function’s major extent (*H*_(4)_=26.41, *p*=3×10^-5^, *ε*^2^=0.30; Fig. 7E), minor extent (*H*_(4)_=15.86, *p*=0.003, *ε*^2^=0.16; Fig. 7G) and their ratio (*H*_(4)_=22.87, *p*=0.0001, *ε*^2^=0.25; Fig. 7I). Again, the major extent decreased through the network layers, though less clearly than in trained IndRNNs and the brain’s timing maps. However, unlike in trained IndRNNs and the brain’s timing maps, the minor extent also decreased and the ratio showed no systematic progression through the network layers (Table S8B). After training on shuffled data only the major extent differed significantly between layers (*H*_(4)_=14.53, *p*=0.006, *ε*^2^=0.05; Fig. 7E), again decreasing through the network layers, although less clearly than in trained IndRNNs and the brain’s timing maps. However, unlike in other IndRNNs and the brain’s timing maps, the minor extent (*H*_(4)_=1.51, *p*=0.825, *ε*^2^=-0.01; Fig. 7G) and ratio (*H*_(4)_=5.06, *p*=0.281, *ε*^2^=0.00; Fig 7I) showed no significant differences between layers.

In higher IndRNN layers, the response functions’ major axis was increasingly angulated towards the event duration dimension, though post hoc analyses of the difference between layers did not reach significance (*P*=10.42, *p*=0.034; Fig. 7K; Table S8A). Again, this progression resembled changes through the brain’s timing map hierarchy (*P*=37.00, *p*=3×10^-5^, Fig. 7L)(Table S8F). The nodes’ response function angulations did not significantly differ between layers in IndRNNs before training (*P*=1.62, *p*=0.806; Fig. 7K) or those trained on shuffled data (*P*=0.10, *p*=0.999; Fig. 7K).

The exponent on event period (or frequency) captures a reduction in the response to repeated events in the input sequence. The humans brain’s hierarchy of visual timing maps shows a clear and highly significant decrease in this exponent (*H*_(9)_=93.84, *p*=3×10^-16^, *ε*^2^=0.55; Fig. 7N; Table S8F). Conversely, no IndRNNs show significant differences between exponents in different layers (Fig. 7M), whether trained in repetitive input sequences (*H*_(4)_=6.02, *p*=0.198, *ε*^2^=0.01), shuffled input sequences (*H*_(4)_=4.50, *p*=0.342, *ε*^2^=0.00), or before training (*H*_(4)_=8.75, *p*=0.068, *ε*^2^=0.07). Instead, in IndRNNs show increasingly bimodal distributions of exponents in higher layers when trained on repetitive input sequences (Table S9). As such, the compressive exponents of individual tuned IndRNN nodes do not follow the clear progressions of period exponents seen in the response functions of large grouped neural populations in the brain.

## Discussion

Responses to visual event timing in the human brain gradually transition from monotonic to tuned responses through the visual processing hierarchy (Hendrikx et al., 2022). Through the brain’s visual field map hierarchy, monotonic responses gradually transition from linearly to compressively increasing in amplitude with duration and frequency, and from depending primarily on event duration to being similarly dependent on duration and frequency. Once tuned responses emerge, the preferred timing, extent, shape, and orientation of their response functions progressively change through a hierarchy of timing-responsive brain areas (Harvey et al., 2020; Hendrikx et al., 2022). Here we asked whether these systematic changes between brain areas could result from the computations inherent in hierarchical recurrent neural networks. To assess which neural architectures produce responses tuned to event timing, we built neural networks of different depths and trained these to predict upcoming time steps in input sequences containing repetitive events. Given this training, recurrent networks of all depths performed the task accurately, and recurrent networks with multiple layers reached higher accuracies than a one-layer network. Untrained networks, networks trained on shuffled input sequences and networks without recurrent connection could not accurately predict the timing of repetitive events. Trained recurrent networks showed a gradual transition from monotonic to increasingly tuned and mixed responses across their successive layers, with networks trained on shuffled input sequences showing similar transitions but less clearly. Both training and recurrency were necessary to produce this emergence of tuned responses. Regardless of training, recurrent networks showed monotonic responses whose transitions through network layers resembled transitions of monotonic responses through the brain’s visual field map hierarchy. However, recurrent networks showed tuned response functions whose transitions through network layers resembled transitions of tuned responses through the brain’s visual timing map hierarchy only after training on predictable repetitive input sequences. These results suggest that the emergence of visual timing-tuned responses and the subsequent hierarchical transformations of these responses result from recurrent neural architecture and predictive processing of sensory event timing.

Although we tested networks of up to five layers, networks with at least two layers did not significantly differ in their maximum prediction accuracy if the networks’ complexity (number of free parameters) was matched. However, one-layer networks showed significantly lower accuracy than all these multi-layer networks. Likewise, the proportions of tuned, monotonic and mixed-responses nodes in the final layer of networks from two to five layers did not differ, while one-layer networks showed fewer tuned and mixed response nodes and more monotonic nodes in their (final and only) layer. Therefore, only two network layers seem to be needed to predict upcoming time steps accurately and transition from monotonic to tuned responses. Five-layer networks primarily demonstrate that this transition from monotonic to tuned responses emerges gradually between network layers, as it does in the visual processing hierarchy (Hendrikx et al., 2022).

To separate effects of successive network operations between layers from effects of network complexity, we investigated the response properties of both parameter-matched (i.e. complexity-matched) networks and layer-size-matched networks, in which network complexity increases with network depth. Deeper parameter-matched networks had very few nodes in each layer, so greater variability between repetitions of the training process. Layer-size-matched networks avoided this limitation and so allowed greater statistical power to investigate differences in response properties between the layers of deeper networks. In layer-size-matched networks, the proportion of tuned nodes in the final layer increased with network depth. Likewise, the proportion of monotonic nodes in the final layer decreased with network depth and complexity, while the proportion of mixed response nodes did not differ between multi-layer networks. As these differences between final layers were absent in parameter-matched networks, we attribute them to differences in complexity (i.e. the number of free parameters) between these networks. The human brain has far greater complexity than the networks we tested here. Our findings suggest that this increased complexity would allow emergence of tuned nodes on a larger scale in the brain, where tuned responses dominate higher-level responses to timing (Harvey et al., 2020; Hendrikx et al., 2022).

These progressive transitions from monotonic to tuned responses were absent in networks with the same architecture before training and networks without recurrent connections, suggesting that training and recurrent connections underlie the emergence of timing-tuned responses. Networks before training showed some tuned and mixed-response nodes, but far fewer than after training. Networks trained on shuffled input sequences, where upcoming time steps could not be predicted based on temporal structure but only based on the proportions of on and off states of the sequence input so far, showed transitions from monotonic to tuned responses though less clearly than networks with the intact training regimen.

Both monotonic and tuned responses have previously been shown in biological neurons and neural networks during timing tasks (Bi & Zhou, 2020; Merchant et al., 2011, 2013; Mita et al., 2009). Monotonic responses alone may be enough to allow animals to make predictions and behavior (Bi & Zhou, 2020). However, tuned responses distribute sensory processing across diversely responding nodes. This increases neural networks’ sensory processing and memory capacity compared to monotonically increasing responses (Crespi, 1999; Palm, 1984) which hold similar information in each node. In line with these findings, we see that more complex networks are more accurate and show more tuned and mixed-response nodes. Together with our fMRI data, this parallel increase in tuning and accuracy in more complex networks also suggests tuned responses may have an important role in linking timing perception and prediction for motor planning, with the sensory and motor stages being dominated by monotonic responses (Hendrikx et al., 2022; Merchant et al., 2011, 2013). Nevertheless, we tested for correlations between the proportion of tuned nodes in a training repetition and the accuracy of the resulting network and found no clear relationship. We also tested whether lesioning the (relatively few) tuned nodes in the final layer decreased accuracy more than lesioning other nodes and found no difference. So tuned nodes alone do not seem to specifically underlie accurate task performance. Instead, it may be that interactions between monotonic nodes (at the input and prediction-generative stages) and tuned nodes (at intermediate stages) are required for accurate performance, particularly considering that one-layer networks would not allow such interactions. Alternatively, the large population of mixed-response nodes may support accurate performance even in the absence of strictly tuned nodes.

We see differences between network layers in the parameters of timing-tuned nodes’ response function that often parallel the differences between timing maps in the brain’s visual time processing hierarchy (Harvey et al., 2020). While the direction of changes between network layers is generally consistent with the direction of changes through the brain’s hierarchy, there are clear differences in the magnitude of the parameters involved. For example, the response function’s extent is several times larger and its angulation is closer to the period dimension than in fMRI data. In both cases, these parameters change to more brain-like magnitudes in higher levels of the network, so it may be that the magnitudes seen in the brain simply reflect the fact that the brain is a far deeper and more complex network than we model here.

The compressive exponents on event period showed very clear and progressive decreases through the brain’s hierarchy of visual field maps and timing maps for monotonic and tuned response functions respectively. This is much less clear in our network’s monotonic nodes, where instead we see bimodal distributions dominated by high exponents in the first layer and low exponents in later layers. Likewise for the tuned nodes, we see transitions from high exponents to increasingly bimodal distributions of high and low exponents. Our fMRI data group the responses of the many neurons in each voxel, and then the many voxels in each map, into a single measurement. This is likely to obscure any bimodal distribution and bring the resulting averaged exponent down as the balance of any bimodal distribution shifts from high to low exponents. So, at the level accessible in fMRI data, such a transition from high exponents to bimodal high and low exponents may simply show the decreasing exponent through the hierarchy that we saw.

Conceptually, responses with high exponents (near one) describe independent responses to each event, while low exponents (near zero) describe a single response to the timing of a group of events regardless of their frequency. The top layer of our network was used to generate a timing prediction, so needed specific responses to each event and a high exponent. There was no such task in our fMRI experiments, so we would expect no such responses. The part of the distribution with exponents near zero, on the other hand, may give an abstract timing representation in both cases, which could be used to both understand the events’ timing and to generate input sequence predictions if needed.

In this study we focused on very simple networks to help reveal which neural processes are necessary to yield the emergence of timing-tuned neurons and the transitions of their response properties seen in the human brain’s visual timing map hierarchy. As such, we did not aim to mimic the complex properties of human visual processing in detail. In the brain, visual processing is fundamentally spatial, while here we used an input sequence with no spatial dimension. Embedding recurrent processing in a spatial image representation may reveal further mechanisms of timing-selective responses. For example, our fMRI data showed that early visual monotonic responses are closely tied to the location of the visual input, while tuned components of later responses are location-invariant. Furthermore, embedding recurrent processing in a spatial image representation would allow more biologically plausible neural architectures. The activity in recurrent units of our IndRNN model (Li et al., 2018) is affected by their own activity in previous time step, but biological neurons don’t synapse with themselves. Likewise, in a more common simple RNN (Elman, 1990; Jordan, 1997) the activity in each node is affected by the activity of all nodes in the same layer (including themselves) at the previous time step (akin to a fully connected neural network), but biological neurons also don’t synapse with all neurons in the same brain areas. If embedded in a spatial representation, the recurrent connections could be limit to nearby nodes within the network layer excluding themselves (akin to a convolutional neural network), a closer approximation of biological neural processing. Furthermore, here we model time in discrete steps. However, time progresses continuously, and biological neurons respond (spike) at specific moments in this timing continuum. To investigate responses to event timing in a biologically plausible way, spiking neural networks with spike timing dependent plasticity would reveal further details of the underlying mechanisms. Both of these properties are possible in neural network models and provide important directions for future research.

## Conclusion

Artificial neural networks allow us to investigate the mechanisms underlying neural responses seen in the brain. Their unique ability to test how the network’s performance properties of these responses are affected by training, different training regimens and different architectures reveals which aspects of neural processing are required to produce these responses. In this case, this approach demonstrates that fundamental neural processes of recurrency and training to predict upcoming inputs are sufficient to explain the emergence of timing-tuned neural responses and the transitions in their response properties seen in the human brain.

## Methods

All used input sequences, scripts, and networks will be made available at publication.

### Input sequences

The input sequence followed the event timings used in (Harvey et al., 2020; Hendrikx et al., 2022 van Ackooij et al., 2022), adjusted to simplify training of our network. In Harvey et al. (2020) and Hendrikx et al. (2022) events consisted of black dots appearing and disappearing in random places on a gray background.. In both cases we used repetitive events with durations and periods varying from 50 to 1000 ms. We described fMRI voxels with timing-tuned response profiles and timing preferences within this range, together with timing preferences outside of this range and responses that monotonically varied with event timing.

Our network’s input sequence here simply describes whether the input state was on (value 1, equivalent to a dot being shown) or off (value 0, equivalent to no dot being shown) during each time step (Fig. 1, top row). Each input sequence consisted of 80 time steps. We describe the consecutive time that the input was on as the “on” duration and the consecutive time that the input was off as the “off” duration. We describe the time from the start of one repeating event to the start of the next (i.e. the sum of the time steps in both the on and off states) as the event period (i.e., 1/frequency). In our previous fMRI experiments, the shortest duration or period was 50 ms, and their smallest increment was also 50 ms. Therefore, we see one time step in our input sequence here as equivalent to 50 ms in our fMRI stimuli, though this does not affect the interpretation of our results. We avoided event timings where event duration and period were equal, as this results in all input sequence being “on” regardless of the event period. Situations where duration is longer than period are not possible, as the duration is part of the period. This resulted in 190 combinations of duration and period, with durations ranging between 1 and 19 time steps and periods ranging between 2 and 20 time steps.

This range of periods results in the 80-time step input sequence containing between 4 events (for a period of 20 time steps) and 40 events (for a period of 2 time steps). This could potentially introduce learning biases in favor of short events, as these are input more often into the network. To avoid this, we included all possible phases for each period, so that the time step of the first event could happen anywhere in the period. The number of possible phases corresponded to the period of the repetitive event, so overall we input a similar number of events with each timing into the network. This resulted in 2660 input sequences in total.

#### Shuffled input sequences

In input sequences with a strong imbalance between on and off states, like input sequences with a long “on” duration, upcoming input states can be predicted accurately even by a network with no timing representation. We therefore also trained our networks on input sequences without periodic events: the input sequences previously described with their time steps shuffled. These sequences keep the same imbalance between on and off states, but with no predictable events and so no specific event duration or period.

### Independent recurrent neural networks

For a neural network to predict upcoming repetitive events, it requires information about previous time steps. Recurrent neural networks (RNNs) were specifically designed to combine information about current network inputs with information about the current and previous network state. At time step *t*, a RNN node receives information about the current network input (in layer 1) or network state (in higher layers) through feedforward connections, and information about the network state in the previous time step (*t*-1) through recurrent connections.

In a simple recurrent network (Fig. 8, right)(Elman, 1990; Jordan, 1997), each node has recurrent connections to itself and all other nodes in its layer. The activations of the nodes of a hidden layer can thus be described as:

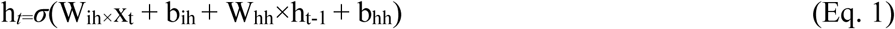

**Fig. 8:**
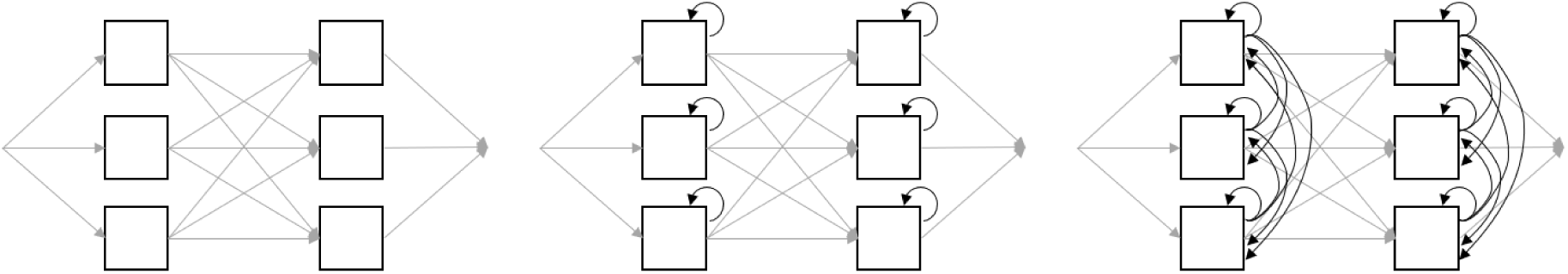
An schematic overview of neural network architectures: a neural network without recurrent connections. (left)**, an IndRNN** (middle)**, and a simple RNN** (right). All examples have two layers with three nodes (represented by squares). Gray arrows indicate between-layer connections and black arrows indicate recurrent connections.

where sigma is the activation function of the nodes in this layer; W_ih_ is the weight matrix between the input nodes and the hidden nodes, which is multiplied by activations of these inputs (x*_t_*) using matrix multiplication; W_hh_ is a matrix with recurrent weights between the hidden nodes in each layer, which is multiplied by the previous activations of these nodes (h*_t-1_*) matrix multiplication; and b_ih_ and b_hh_ are the input and recurrent biases, respectively.

The dense recurrent connections make the nodes’ responses in a layer dependent on each other, making it hard to assess responses in individual nodes within a single layer. Furthermore, the simple RNN architecture has more complex within-layer connections than a biological system does. To illustrate, in the visual processing hierarchy, biological neurons’ activity depend on: 1) excitatory and inhibitory neurons that input into this neuron from previous visual field maps, which can be modeled using between-layer connections; 2) its own polarization that slowly decays unless it is depolarized through an action potential, which can be modeled using recurrency between a node and itself; 3) lateral connections of neurons of the identical receptive field map (which grow over the hierarchy), which can be modeled as lateral non-recurrent connections between nodes within one layer; and 4) top-down connections with inputs from brain areas later in the hierarchy. The third type of connections are significant in spatial location processing, however since we did not vary the spatial location of visual inputs they can be disregarded in our setup. The fourth type of connections were overly complex for the current setup: we will not model influence of attention or other higher-level processes in our networks. A network simulating a biological neural system should then allow complex interactions between layers, but limited interactions within layers.

Therefore, for our predictive task we used a specific version of the RNN: the independent RNN (Fig. 8, middle)(Li et al., 2018). In the IndRNN, each node only has a recurrent connection with its own state at time step *t*-1. This makes the differences in responses of the nodes in different layers more transparent. Each node is also (fully) connected to all nodes in the previous layer (at time step *t*). The activations of the nodes of a hidden layers of the IndRNN we used can be described as follows (Eq. 2):

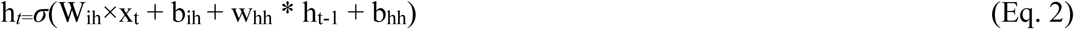

Here, w_hh_ is a vector with recurrent weights from the hidden nodes in the layer back to themselves, that is multiplied by the previous activations of these nodes (h*_t-1_*) using the Hadamard product. We used a sigma activation function for the hidden nodes.

#### Parameters of the IndRNN

In the human brain, tuned responses to event timing gradually emerge through the visual processing hierarchy (Hendrikx et al., 2022). We therefore hypothesized that tuning in IndRNN layers may also emerge gradually, requiring multiple layers for timing tuned responses to develop. Therefore, we built IndRNNs with depths of one to five layers. To allow comparisons between networks of different depths, we focus on a set of networks with similar numbers of free parameters (i.e., the total number of weights and biases) for all network depths. We refer to these as the parameter-matched networks. Here, the one-layer IndRNN had 77 nodes per layer, the two-layer IndRNN had 16 nodes per layer, the three-layer IndRNN had 12 nodes per layer, the four-layer IndRNN had 9 nodes per layer, and the five-layer IndRNN had 8 nodes per layer.

Deeper parameter-matched networks had few nodes in each layer, making it hard to compare the distribution of response properties over the nodes in their layers. To allow such comparisons, we also built a set of networks with 16 nodes per layer regardless of network depth. We refer to these as layer-size-matched networks.

We used a ReLU activation function on the hidden nodes. We used a batch size of 50 input sequences. We initialized the weights of the network using He initialization (He et al., 2015)(He et al., 2015) and the biases at 0. During training we applied Adam optimization with a weight decay of 10^-8^. The initial learning rate was set to 0.002. We used layer normalization for each hidden layer (Ba et al., 2016). We did not apply dropout, since we wanted to reduce the amount of necessary nodes per layer to a minimum and we do not expect the networks to generalize to other tasks.

The target output is binary (“on” or “off”, i.e., 1 or 0), but the network nodes’ activations are continuous and 0 or positive (due to the ReLU activations). In order to transform the network’s output to a value between 0 and 1 we applied a sigmoidal activation function to the output node. Because the target output is binary, we applied a Binary Cross Entropy (BCE) loss function for back propagation. The BCE per batch was the mean loss over all time steps of all 50 input sequences.

### Neural networks without recurrency

To investigate whether recurrency is necessary for predicting event timing, we also built a five-layer network without recurrency (Fig. 8A). As this network cannot access network states from previous time steps, it should be unable to track the timing of its input sequence. We therefore predict such a network will be unable to develop tuned responses to the timing of this input sequence. We kept the parameters and settings for this network as similar as possible to the IndRNNs. Its number of free parameters matched that of the parameter-matched networks, resulting in 9 nodes per layer.

### Training

We trained each network for 10000 epochs using a train-test split of 0.8-0.2. Note that the same timings can be in the train and test set, but the same phases of this timing will never be present in both. This means that networks can be familiar with a specific timing, but their performance on predicting that timing will not hinge on their memory of an exact input sequence.

Our fMRI experiments (Harvey et al., 2020; Hendrikx et al., 2022; van Ackooij et al., 2022), typically vary the timing of stimuli gradually, to reduce the influence of possible adaptation effects. For neural network models, such carry-over effect are easily avoided by resetting the network nodes’ hidden states to 0 before presentation of every new input sequence. We therefore presented input sequences with different timings and phases in a random order.

Due to their random initialization weights, networks with the same architecture vary in training loss, task performance, and node activations. To reliably determine these properties for a particular network architecture, we trained and tested each architecture 50 times with different random initialization weights. We saved these initialization weights and used these same initial weights when training on shuffled input sequences (we used the parameter-matched five-layer network for shuffled sequences). To minimize variability and facilitate comparisons between networks, we trained each network with the same train-test split. During training, each network was presented with a different randomized order of input sequences that was shuffled between epochs.

### Evaluation of timing prediction performance

#### Training loss

For each network we saved its training loss over epochs. We compared the loss on the last epoch of the parameter-matched IndRNNs and that of the layer-size-matched IndRNNs between different network depths using a Kruskal Wallis (scipy.stats; Kruskal & Wallis, 1952) followed by pairwise Dunn’s tests for post hoc comparisons (scipy.stats; Dunn, 1964), with a Holm-Šidák multiple comparisons correction (Šidák, 1967). We used Wilcoxon signed-rank tests (scipy.stats; Wilcoxon, 1945) to compare the loss in the last epoch of the five-layer IndRNN to its initializations trained on shuffled input sequences. We used Mann Whitney U tests (scipy.stats; Mann & Whitney, 1947) to compare the loss in the last epoch of the five-layer IndRNN to the five-layer NN without recurrency. We also used Mann Whitney U tests to compare the loss of the parameter-matched and layer-size matched IndRNNs.

#### Accuracy of timing prediction

We assessed each network’s accuracy at predicting the upcoming time step of the input sequence on the test split of the input sequences. Because the input sequences could have only two states (on and off), here we binarized the networks’ predictions. Our accuracy score per time step is then 1 if this prediction is correct and 0 if it is incorrect. Here we noted that prediction is relatively trivial when the input sequence’s state does not change between time steps, but more difficult when the inputs in the current and upcoming time step differ. We therefore assessed accuracy in two ways. First, per-event accuracy quantifies the average accuracy over the time steps encompassing an entire event, most of which will be relatively trivial in longer events where most of the time steps are unchanged from the previous time steps. Second, per-state-change accuracy quantifies the average accuracy only at the time steps where the input sequence changes state, which then always compares accuracy on two similarly difficult time steps regardless of the event’s timing.

For both accuracy measures, we only assessed accuracy during whole events where the network could theoretically predict the next time step in the input sequence. At the first time step (e.g. time step 1 in Fig. 1), the network has no information about the input sequence’s timing. Time steps where the input changes state reveal the temporal structure of the event. The first state change (e.g., time step 8 in Fig. 1) does not give complete information about timing: it only reveals a lower bound for the length of the initial on or off state. The second state change (e.g., time step 17 in Fig. 1) unambiguously reveals the length of the second on or off state, but still does not reveal the length of the other state. The third state change (e.g., time step 28 in Fig. 1) reveals the length of the initial state. Only at this time step has the network had one whole event input, and so has enough information to make accurate predictions in all further time steps. So, from this third state change on, we can assess the network’s accuracy.

We only assess the accuracy of predictions of whole events. We do this because some parts of the event may be easier to predict than others, which may in turn bias the calculated accuracy score. We define the prediction of whole events as predictions encompassing an entire period, with the full set of the event’s consecutive “on” times steps followed by the full set of its consecutive “off” time steps, or vice versa. The first predicted event for which we assess the accuracy therefore starts at the state change in the *predicted* input sequence that follows the third state change of the input sequence (e.g., time step 36 in Fig. 1). From this predicted event onwards, we assess all following predicted events that still encompass the entire period (e.g., last evaluated time step is time step 75 in Fig. 1).

For each repetition we computed the mean accuracy over all input sequences. We compared per-event and per-state-change accuracy (separately) between networks of different depths using a Kruskal-Wallis test (scipy.stats; Kruskal & Wallis, 1952) followed by post hoc pairwise Dunn’s tests (scipy.stats; Dunn, 1964) with a Holm-Šidák correction (Šidák, 1967). For this and all other Kruskal-Wallis tests we computed effect size *ε*^2^ as *(H-statistic – amount groups + 1) / (total amount datapoints – 1).* To assess differences in maximum performance of the parameter-matched networks, we performed a similar comparison for only the best 25 repetitions of each network depth.

We compared the variance in accuracy over network repetitions between layers separately for the parameter-matched and layer-size-matched networks using a Brown-Forsythe test (scipy.stats.levene; Brown & Forsythe, 1974) followed by posthoc pairwise Brown-Forsythe tests, corrected for multiple comparisons using a Holm-Šidák correction (statsmodels.stats; Šidák, 1967).

We compared the accuracies of the five-layer IndRNN to the same network untrained initializations and those of the five-layer IndRNN to the same network initializations trained on shuffled input sequences using (paired) Wilcoxon signed-rank tests (scipy.stats; Wilcoxon, 1945). We compared the accuracies of the five-layer IndRNN to the five-layer NN and those of the five-layer IndRNN with 8 nodes per layer to the five-layer IndRNN with 16 nodes per layer using a Mann Whitney U test (independent) (scipy.stats; Mann & Whitney, 1947).

#### Stability of accuracy across timings

To assess whether each event timing elicits a similar accuracy pattern or whether there are systematic differences in accuracy between event timings, we computed average per-event and per-state-change accuracies for each timing present in the test set. We compared the variance of these average accuracies (i.e. how much accuracy varies with timing) between network depths, using a Kruskal-Wallis test (scipy.stats; Kruskal & Wallis, 1952) followed by post hoc pairwise Dunn’s tests (scipy.stats; Dunn, 1964) with a Holm-Šidák correction (Šidák, 1967). We again did a similar comparison for only the best 25 repetitions of each network depth.

### Parameterizing node response functions

To parameterize the response functions of the nodes, we computed a per-event response for each timing. We used the same method of selecting the set of time steps as we used for computing the per-event accuracies, selecting only whole, predictable events. For each node we summed the activity across these time steps and divided this by the number of events during these time steps (giving the summed response in the average whole, predictable event). We fit monotonic and tuned response functions to these average per-event responses from all event timings. The response functions we used were the best-fitting response functions in previous studies of the human brain’s sensory event timing-selective responses (Harvey et al., 2020; Hendrikx et al., 2022; van Ackooij et al., 2022).

Here we started with all possible input sequences (from both the training and the test sets used during training). We then split these into two halves, each containing every timing, but with differently shifted phases in each half. We fit each response function’s free parameters on one half of the data and evaluated the resulting function’s fit on the complementary half (using both halves for fitting and both for evaluation).

We fit both functions to each node’s activation using scipy’s curve_fit in Python. Initially, each free parameter of the response function was randomly set within specified bounds (Table S11). These parameter values were optimized to minimize the sum of squared residual to the measured activations. This was done with an *ftol* value (indicating the expected change in the cost function while improving the parameters) of 10^-X^ where X=4 for the first iteration. X was increased by 1 on each iteration until X=14 or no solution could be found for larger values of X. If no solution could be found in the first iteration, this was repeated with a new random value of each free parameter until a solution was found. We used the resulting parameter values to compute the cross-validated variance explained (VE) of the resulting function on the complementary half of the data: the proportion of variance in complementary half of node response that was captured by the resulting response function.

In order for *ftol* to have a similar effect in all nodes, we normalized each node’s response before (re)fitting the response functions. Nodes with the same per-event response for each timing were disregarded in the fitting process, as these responses have no variance for the response function fits to explain.

#### Monotonic response functions

Monotonic response functions have previously been demonstrated to capture effects of event timing on fMRI responses in early visual areas (Hendrikx et al., 2022). Following these results, the monotonic response function we used here has two independent components that scale with the duration of the “on” state of the stimulus sequence and the frequency of event onsets (Stigliani et al., 2017) i.e. 1/period, respectively. These two components each included a free parameters allowing a compressive exponential nonlinearity in the relationship between response amplitude and event duration or frequency (Zhou et al., 2018).

Note that to the network, events being “on” or “off” merely describes a value of the binary input. Therefore, responses could monotonically increase with either the event’s “on” or “off” duration. We therefore fit response functions that increased with either the “on” or “off” duration (as well as increasing with frequency) and chose whichever best predicted the responses under cross-validation. For consistency with the tuned response functions, we describe the responses to each event as a function of its period (i.e. 1/frequency). So, our monotonic response functions can be expressed as:

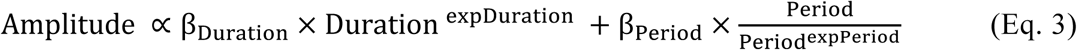

Where Duration can represent the event’s “on” or “off” duration. Here, expDuration and expPeriod are the compressive exponent on the chosen duration and period respectively, in the range 0–1. β_Duration_ and β_Period_ capture the relative amplitudes of the chosen duration and period components, respectively.

The responses of the nodes in a network cannot be negative, as we are applying rectified linear activation functions. Therefore, for consistency with previous work, we did not allow the betas to become negative (though very similar results were found when we allowed this: Fig. S6 & Table S11).

#### Tuned response functions

Tuned response functions have previously been demonstrated to capture effects of event timing on fMRI responses in association cortices (Harvey et al., 2020; Hendrikx et al., 2022; Protopapa et al., 2019). Following these results, the tuned response function we used here is described by a two-dimensional anisotropic Gaussian function of the duration of each event’s “on” state and its period. The function describes the response amplitudes to each event separately. However, when assessing events over the entire input sequence, response amplitudes could also increase with event frequency. Because response amplitudes to each event in the input sequence may also be affected by their frequency (regardless of the parameters of the Gaussian function), we scaled the per-event response by a compressive exponent on frequency (Harvey et al., 2020).

Again, a node’s activations could vary as a Gaussian function of either the event’s “on” or “off” duration. We therefore fit response functions that depended on either “on” or “off” duration (as well as period) and chose whichever best predicted the activations under cross-validation. So, our tuned response functions can be expressed as:

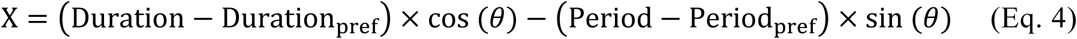

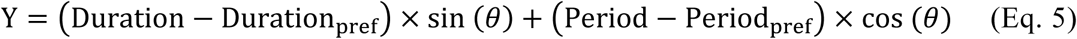

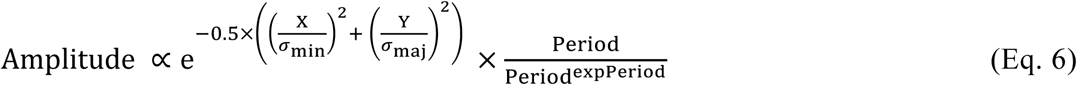

Where Duration could represent the event’s “on” or “off” duration. The six free parameters of the response function are: the preferred duration (Duration_pref_) and the preferred period (Period_pref_) around which the Gaussian function’s mean is centered; the standard deviations along its major and minor axes (*σ*_maj_ and *σ*_min_, in the range 0–10); the angulation of its major axis (*θ*); and the compressive exponent on the period to which the response was scaled (expPeriod) to account for global effects of event frequency.

Again, we do not allow the proportionality between the node activation and the response function (Eq. 6) to become negative (though very similar results were found when we allowed this: Fig. S6 & Table S11).

#### Classification of response function types

After fitting, we classified each node according to the response function with the highest cross-validated VE. For nodes fit best by the tuned response functions, nodes where the fit response function’s standard deviation along both axes reached their maximal allowed values do not show clear evidence of tuned responses as they closely resemble responses with no variance. We therefore did not classify these nodes into either response function category.

Furthermore, nodes where the cross-validated VE of their best fitting response function was very low did not show clear evidence of either type of response function. Following previous fMRI studies (Hendrikx et al., 2022), we used a VE threshold of 0.2, below which we did not classify nodes into either response function category. A higher threshold of 0.8 produced very similar results (Fig. S7 & Table S12).

For nodes best fit by the tuned response function, we classified those with particular combinations of tuned function parameters into a third category: mixed response function. This category was again inspired by fMRI results where many neural populations show characteristics of both monotonic and tuned response functions (Hendrikx et al., 2022). This mixed response function category contained nodes where the peak of the tuned response function (determined with scipy’s minimize) was outside the set of event timings that the network was trained to predict. For example, the responses of such a node may have a clear peak duration within the tested range, but no clear peak period. This can happen in two ways. First, the duration or period of the tuned response function’s peak could be below 1 time step or above 20 time steps, the tested range. Second, the peak state of the tuned response function could be larger than the peak period, an event timing which is impossible because the state is part of the period and so cannot be longer than the period. In both of these cases, we do not see this peak in the fit responses. However, the tuned response function can only outperform the monotonic response function because features of the response amplitudes within the tested range are consistent with a tuned response.

#### Comparing proportions of response function types

For each network repetition we assessed the proportion of nodes best fit by monotonic, mixed, or tuned response functions. Within each network depth, computed each response type as a proportion of nodes each network layer (including nodes that are not classified for any of these response types), and compared the proportions between layers. We did these analyses for the parameter-matched IndRNNs, the IndRNNs trained on shuffled data, the untrained initializations of the IndRNNs and the networks without recurrency. Between network depths, we compared the proportion of nodes in the last layer. We did these analyses for the parameter-matched IndRNNs and the layer-size-matched IndRNNs. For each response function type separately, we performed a Kruskal-Wallis test (scipy.stats; Kruskal & Wallis, 1952), followed by post hoc Dunn’s tests (scipy.stats; Dunn, 1964) with a Holm-Šidák correction (Šidák, 1967) to compare proportions between network layers, and Dunn’s tests (scipy.stats; Dunn, 1964) with a Holm-Šidák correction (Šidák, 1967) to compare proportions between network depths.

### Evaluating properties of the response functions of the nodes

We assessed the parameters of the best-fitting response functions between different layers of the same network depths.

For the monotonic responses, we assessed the compressive exponents on duration and period, and whether the ratio of contribution of duration and period components to the response (β_duration_/β_period_) differed between network depths and layers. In order to make a fair comparison, the betas were rescaled to be fit on normalized response components. Nodes with an exponent on duration of 0 or an exponent on period of 1 create the same activation for all durations or periods, respectively. Therefore, the beta ratio for these nodes was set to 0 or infinite, respectively. Similarly, for nodes with a beta ratio of 0 or infinite, duration and period exponents were set to 0 or 1, respectively. All comparisons were performed using a Kruskal-Wallis (scipy.stats; Kruskal & Wallis, 1952) followed by Dunn’s post hoc (scipy.stats; Dunn, 1964) with Holm-Šidák correction (Šidák, 1967). To compare the patterns in the data, we also performed these tests on the nodes of the layer-size-matched networks before training, layer-size-matched networks trained on shuffled data and parameter-matched networks without recurrency. Furthermore, we performed the same analysis on previously published data about monotonic parameters in visual field maps recorded with fMRI (Hendrikx et al., 2022)

For the tuned responses, we assessed the preferred timings, the extents around these timings, the angulation of this preference towards the x-axis, and the exponent on period between layer-size-matched layers of networks before training, layer-size-matched networks trained on shuffled data and networks trained on repetitive events. For all parameters, except the angulation, we performed using a Kruskal-Wallis (scipy.stats; Kruskal & Wallis, 1952) followed by Dunn’s post hoc (scipy.stats; Dunn, 1964) with Holm-Šidák correction (Šidák, 1967). Furthermore, we performed the same analysis on previously published data about tuned parameters in timing maps recorded with fMRI (Harvey et al., 2020).

For the angulation parameter, we normalized the angulation angles between 0° and 180°. We multiplied the angles by 2 to allow performing a circular common median test (pycircstat.tests.cmtest), followed by posthoc pairwise circular common median tests with Holm-Šidák correction (statsmodels.stats; Šidák, 1967).

For the exponent of the monotonic and tuned response function, we assessed whether the data was bimodally distributed the dip test of unimodality (Hartigan & Hartigan, 1985). For each training regimen, we corrected for each layer’s statistical tests using an FDR correction (Benjamini & Hochberg, 1995).

Note that the statistical tests and included datapoints presented here may slightly differ from the original studies in the brain (Harvey et al., 2020; Hendrikx et al., 2022), to create consistency with the statistical procedures used in the current study. However, the reported trends are very similar.

