## Supplementary materials for "Transitions from monotonic to tuned responses in recurrent neural network models during timing prediction"

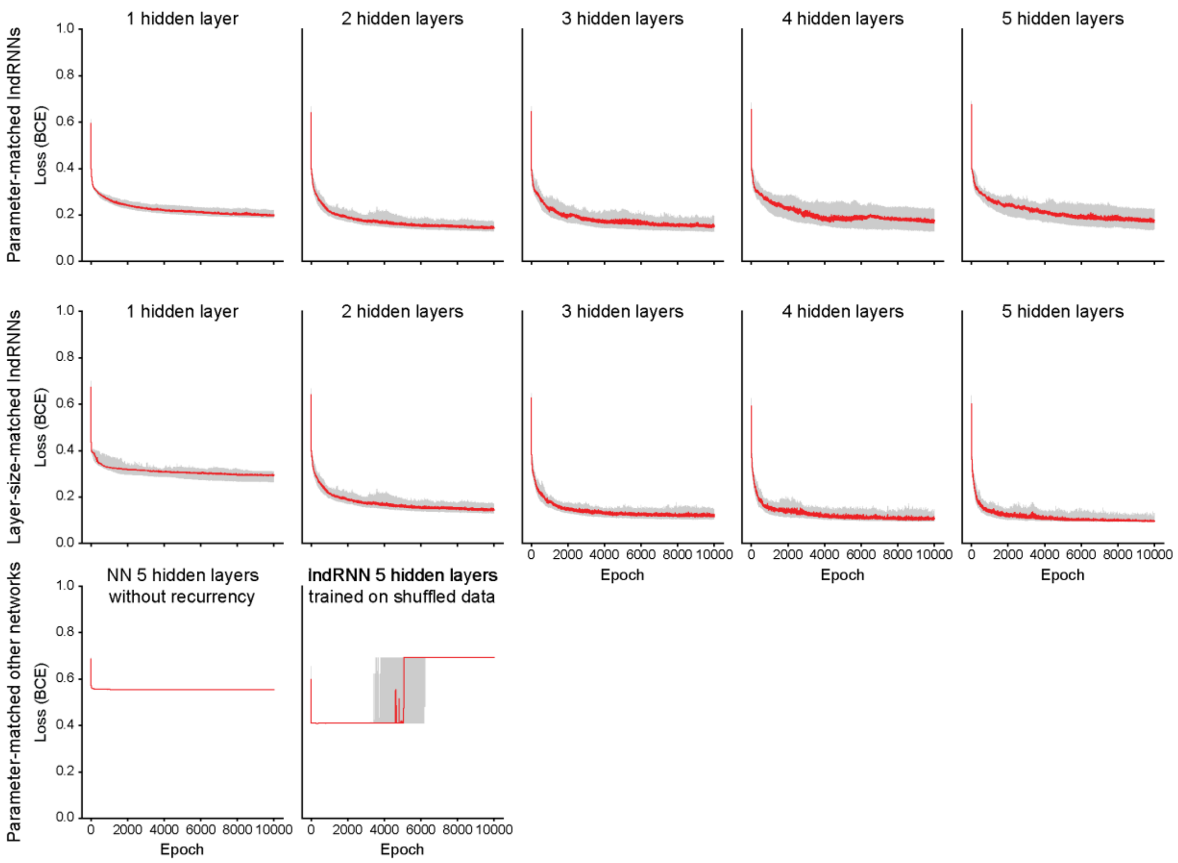


**Supplementary Fig. 1: Loss functions over 10,000 epochs of the parameter-matched IndRNNs, the layer-sized matched IndRNNs, a parameter-matched 5-layer neural network without recurrency, and a parameter-matched 5-layer IndRNN trained on shuffled data.** The IndRNN trained on shuffled data has the same initializations as the one trained on timing data. The 5 layer IndRNN trained on timing data converges on significantly lower loss than the neural network and the IndRNN trained on shuffled data. The 5 layer IndRNN with 16 nodes in each layer (layer-size matched) has a lower loss than the 5 layer IndRNN with 8 nodes in each layer (parameter-matched) (supplementary table 2). The IndRNN trained on shuffled data seems to have 2 strategies and eventually all repetitions fall into the suboptimal minimum over 10,000 epochs.

| | Median [IQR] | Test statistics | | | | |
|---|---|---|---|---|---|---|
| | | 1 layer | 2 layers | 3 layers | 4 layers | 5 layers |
| **1 layer** | 0.20 [0.19, 0.22] | | $1 \times 10^{-9}$ | $9 \times 10^{-6}$ | 0.008 | 0.005 |
| **2 layers** | 0.14 [0.13, 0.17] | -6.44 | | 0.346 | 0.009 | 0.013 |
| **3 layers** | 0.15 [0.13, 0.19] | -4.90 | 1.54 | | 0.346 | 0.346 |
| **4 layers** | 0.18 [0.14,0.22] | -3.26 | 3.18 | 1.64 | | 0.858 |
| **5 layers** | 0.18 [0.14,0.22] | -3.43 | 3.01 | 1.46 | 0.18 | |

**Supplementary Table 1A: Loss of the final epochs is highest for a one-layer parameter-matched network, and lower for a two- than a four- or five-layer parameter-matched network.** Given are the median [IQR] loss over network repetitions and the test statistics comparing these groups. *Z*-values shaded in black, corresponding *p*-values shaded in gray. Significant differences underlined.

| | Median [IQR] | Test statistic | *p*-value |
|---|---|---|---|
| **vs. parameter-matched 5-layer IndRNN** | | | |
| **Parameter-matched 5-layer IndRNN trained on shuffled data** | 0.69 [0.69, 0.69] | $W = 0$ | $2 \times 10^{-15}$ |
| **Parameter-matched 5-layer NN recurrent connections** | 0.55 [0.55, 0.55] | $W = 0$ | $7 \times 10^{-18}$ |
| **Layer-size-matched 5-layer IndRNN** | 0.10 [0.09, 0.11] | $W = 2233$ | $1 \times 10^{-11}$ |
| **vs. parameter-matched 4-layer IndRNN** | | | |
| **Layer-size-matched 4-layer IndRNN** | 0.11 [0.10, 0.13] | $W = 2142$ | $8 \times 10^{-10}$ |
| **vs. parameter-matched 3-layer IndRNN** | | | |
| **Layer-size-matched 3-layer IndRNN** | 0.12 [0.10, 0.14] | $W = 2012$ | $2 \times 10^{-7}$ |
| **vs. parameter-matched 1-layer IndRNN** | | | |
| **Layer-size-matched 1-layer IndRNN** | 0.29 [0.27, 0.31] | $W = 27$ | $4 \times 10^{-17}$ |

**Supplementary Table 1B: Increased complexity decreases the loss in the final epoch, while training on shuffled input sequences and lack of recurrency increase that loss.** Given are the median [IQR] loss over network repetitions of the last epoch. The test statistics comparing the loss to the parameter-matched IndRNNs in the bold single row and the corresponding *p*-values. Significant comparisons underlined.

| | Median [IQR] | Test statistics | | | | |
|---|---|---|---|---|---|---|
| | | **1 layer** | **2 layers** | **3 layers** | **4 layers** | **5 layers** |
| **1 layer** | 0.29 [0.27, 0.31] | | $2 \times 10^{-6}$ | $4 \times 10^{-15}$ | $8 \times 10^{-19}$ | $2 \times 10^{-27}$ |
| **2 layers** | 0.14 [0.13, 0.17] | -5.11 | | 0.010 | $3 \times 10^{-4}$ | $2 \times 10^{-8}$ |
| **3 layers** | 0.12 [0.10, 0.14] | -8.13 | -3.02 | | 0.330 | 0.010 |
| **4 layers** | 0.11 [0.10, 0.13] | -9.10 | -3.99 | -0.97 | | 0.099 |
| **5 layers** | 0.10 [0.09, 0.11] | -11.05 | -5.95 | -2.93 | -1.95 | |

**Supplementary Table 1C: Loss decreases as complexity of the layer-size-matched networks increases.** Given are the median [IQR] loss over network repetitions and the test statistics comparing these groups. *Z*-values shaded in black, corresponding *p*-values shaded in gray. Significant differences underlined.

| **Per-event accuracies** | | | | | | |
|---|---|---|---|---|---|---|
| | **Median [IQR]** | **Test statistics** | | | | |
| | | **1 layer** | **2 layers** | **3 layers** | **4 layers** | **5 layers** |
| **1 layer** | 0.93 [0.92, 0.94] | | $2 \times 10^{-7}$ | $6 \times 10^{-6}$ | 0.029 | 0.013 |
| **2 layers** | 0.96 [0.95, 0.97] | 5.63 | | 0.678 | 0.330 | 0.175 |
| **3 layers** | 0.97 [0.94, 0.97] | 4.97 | -0.66 | | 0.162 | 0.458 |
| **4 layers** | 0.94 [0.92, 0.96] | 2.86 | -2.77 | -2.11 | | 0.678 |
| **5 layers** | 0.95 [0.91, 0.96] | 3.65 | -1.99 | -1.33 | 0.78 | |
| **Per-state change accuracies** | | | | | | |
| **1 layer** | 0.76 [0.69, 0.82] | | 0.0002 | 0.0009 | 0.311 | 0.037 |
| **2 layers** | 0.90 [0.81, 0.94] | 4.33 | | 0.656 | 0.078 | 0.438 |
| **3 layers** | 0.88 [0.76,0.96] | 3.88 | -0.45 | | 0.205 | 0.647 |
| **4 layers** | 0.80 [0.63, 0.93] | 1.80 | -2.53 | -2.08 | | 0.647 |
| **5 layers** | 0.87 [0.65, 0.94] | 2.83 | -1.50 | -1.05 | 1.03 | |

**Supplementary Table 2A: per-event and per-state change accuracies for the parameter-matched networks compared between networks depths.** Given are the median [IQR] accuracies over network repetitions. The test statistics comparing each pair of network depths (post-hoc Dunn's test) are given in black cells, with corresponding corrected *p*-values (Holm-Šidák) of this comparison in the gray cells. Significant comparisons underlined

| Per-event accuracies | | | | | | |
|---|---|---|---|---|---|---|
| | Median [IQR] | Test statistics | | | | |
| | | 1 layer | 2 layers | 3 layers | 4 layers | 5 layers |
| **1 layer** | 0.94 [0.93, 0.94] | | $3 \times 10^{-9}$ | $2 \times 10^{-10}$ | $1 \times 10^{-5}$ | $7 \times 10^{-9}$ |
| **2 layers** | 0.97 [0.96, 0.97] | 6.27 | | 0.928 | 0.513 | 0.928 |
| **3 layers** | 0.97 [0.96, 0.97] | 6.68 | 0.42 | | 0.289 | 0.928 |
| **4 layers** | 0.96 [0.95, 0.97] | 4.77 | -1.50 | -1.92 | | 0.527 |
| **5 layers** | 0.96 [0.96, 0.97] | 6.14 | -0.13 | -0.55 | 1.37 | |
| **Per-state change accuracies** | | | | | | |
| **1 layer** | 0.82 [0.78, 0.85] | | $8 \times 10^{-7}$ | $3 \times 10^{-9}$ | 0.0005 | $4 \times 10^{-7}$ |
| **2 layers** | 0.94 [0.92, 0.95] | 5.32 | | 0.706 | 0.546 | 0.876 |
| **3 layers** | 0.96 [0.93, 0.97] | 6.28 | 0.96 | | 0.120 | 0.706 |
| **4 layers** | 0.93 [0.87, 0.96] | 3.98 | -1.34 | -2.31 | | 0.513 |
| **5 layers** | 0.94 [0.90, 0.97] | 5.48 | 0.16 | -0.81 | 1.50 | |

**Supplementary Table 2B: per-event and per-state change accuracies for the parameter-matched networks compared between networks depths for top 25 repetitions.** Given are the median [IQR] accuracies over network repetitions. The test statistics comparing each pair of network depths (post-hoc Dunn's test) are given in black cells, with corresponding corrected *p*-values (Holm-Šidák) of this comparison in the gray cells. Significant comparisons underlined

| Per-event accuracies | | | | | | |
|---|---|---|---|---|---|---|
| | Median [IQR] | Test statistics | | | | |
| | | 1 layer | 2 layers | 3 layers | 4 layers | 5 layers |
| **1 layer** | 0.87 [0.86, 0.89] | . | $1 \times 10^{-6}$ | $8 \times 10^{-15}$ | $7 \times 10^{-20}$ | $1 \times 10^{-26}$ |
| **2 layers** | 0.96 [0.95, 0.97] | 5.22 | . | 0.016 | $2 \times 10^{-4}$ | $9 \times 10^{-8}$ |
| **3 layers** | 0.97 [0.96, 0.97] | 8.03 | 2.81 | . | 0.234 | 0.016 |
| **4 layers** | 0.97 [0.96, 0.98] | 9.37 | 4.15 | 1.34 | . | 0.234 |
| **5 layers** | 0.98 [0.97, 0.98] | 10.90 | 5.69 | 2.87 | 1.53 | . |
| **Per-state change accuracies** | | | | | | |
| **1 layer** | 0.38 [0.28, 0.45] | . | $9 \times 10^{-7}$ | $2 \times 10^{-14}$ | $1 \times 10^{-20}$ | $2 \times 10^{-26}$ |
| **2 layers** | 0.90 [0.81, 0.94] | 5.25 | . | 0.025 | $8 \times 10^{-5}$ | $2 \times 10^{-7}$ |
| **3 layers** | 0.96 [0.90, 0.99] | 7.89 | 2.64 | . | 0.176 | 0.013 |
| **4 layers** | 0.98 [0.93, 1.00] | 9.57 | 4.32 | 1.68 | . | 0.206 |
| **5 layers** | 0.99 [0.97, 1.00] | 10.83 | 5.59 | 2.95 | 1.26 | . |

**Supplementary Table 2C: per-event and per-state change accuracies for the layer-size-matched networks compared between networks depths.** Given are the median [IQR] accuracies over network repetitions. The test statistics comparing each pair of network depths (post-hoc Dunn's test) are given in black cells, with corresponding corrected *p*-values (Holm-Šidák) of this comparison in the gray cells. Significant comparisons underlined

| Per-event accuracy variances | | | | | |
|---|---|---|---|---|---|
| | 1 layer | 2 layers | 3 layers | 4 layers | 5 layers |
| **1 layer** | | 0.948 | 0.218 | 0.008 | 0.003 |
| **2 layers** | $W = 0.08$ | | 0.315 | 0.019 | 0.008 |
| **3 layers** | $W = 4.32$ | $W = 3.28$ | | 0.587 | 0.510 |
| **4 layers** | $W = 11.39$ | $W = 9.49$ | $W = 1.31$ | | 0.948 |
| **5 layers** | $W = 13.88$ | $W = 11.66$ | $W = 1.97$ | $W = 0.06$ | |
| **Per-state-change accuracy variances** | | | | | |
| **1 layer** | | 0.969 | 0.277 | 0.016 | 0.030 |
| **2 layers** | $W = 0.00$ | | 0.277 | 0.017 | 0.031 |
| **3 layers** | $W = 3.85$ | $W = 3.71$ | | 0.604 | 0.662 |
| **4 layers** | $W = 10.53$ | $W = 10.16$ | $W = 1.62$ | | 0.969 |
| **5 layers** | $W = 8.76$ | $W = 8.47$ | $W = 1.07$ | $W = 0.05$ | |

**Supplementary Table 3: per-event and per-state change accuracy variances between network repetitions for the parameter-matched networks compared between networks depths.** Given are the test statistics of pairwise Levene's tests in the black cells, with corresponding corrected *p*-values (Holm-Šidák) of this comparison in the gray cells. Significant comparisons underlined

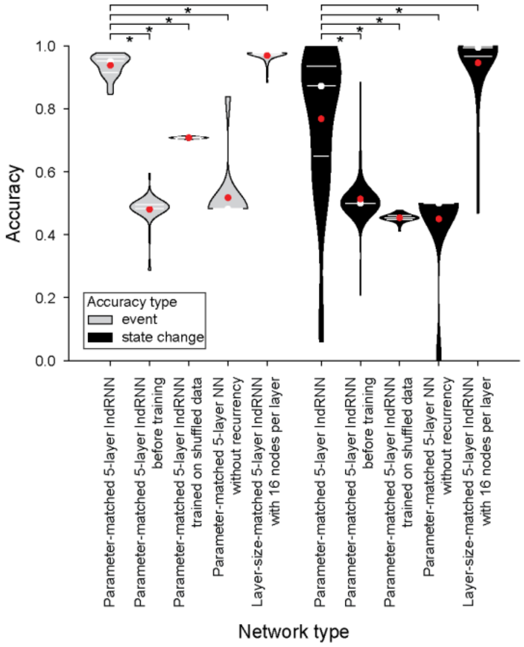


**Supplementary Fig. 2: Target prediction accuracies higher with increased complexity and lower with training on shuffled input sequences and without recurrency**. Stars indicated significant differences between the network types. All network types were only compared to the parameter-matched five-layer IndRNN. Test statistics given in Supplementary Table 4. Format follows Fig. 2A.

| | **Median [IQR]** | **Test statistic** | ***p*-value** |
|---|---|---|---|
| | **Per-event accuracy** | | |
| **Regular IndRNN** | 0.95 [0.91, 0.96] | | |
| **IndRNN before training** | 0.48 [0.48, 0.50] | $W = 0$ | $2 \times 10^{-15}$ |
| **IndRNN trained on shuffled data** | 0.71 [0.71, 0.71] | $W = 0$ | $2 \times 10^{-15}$ |
| **NN without recurrent connections** | 0.48 [0.48, 0.48] | $W = 2500$ | $2 \times 10^{-19}$ |
| **IndRNN with 16 nodes per layer** | 0.98 [0.97, 0.98] | $W = 323$ | $2 \times 10^{-10}$ |
| | **Per-state-change accuracy** | | |
| **Regular IndRNN** | 0.87 [0.65, 0.94] | | |
| **IndRNN before training** | 0.50 [0.50, 0.50] | $W = 107$ | $2 \times 10^{-8}$ |
| **IndRNN trained on shuffled data** | 0.45 [0.45, 0.46] | $W = 57$ | $1 \times 10^{-10}$ |
| **NN without recurrent connections** | 0.50 [0.50, 0.50] | $W = 2275$ | $1 \times 10^{-13}$ |
| **IndRNN with 16 nodes per layer** | 0.99 [0.97, 1.00] | $W = 362$ | $9 \times 10^{-10}$ |

**Supplementary Table 4: Target prediction accuracies higher with increased complexity and lower with training on shuffled input sequences and without recurrency.** Statistics corresponding to Supplementary Fig. 2. Given are the median [IQR] per-event and per-state-change accuracies of each network type. The test statistics (Wilcoxon against shuffled and before training; others Mann Whitney U) comparing the accuracies to the parameter-matched five-layer IndRNN and the corresponding *p*-values. Significant comparisons underlined.

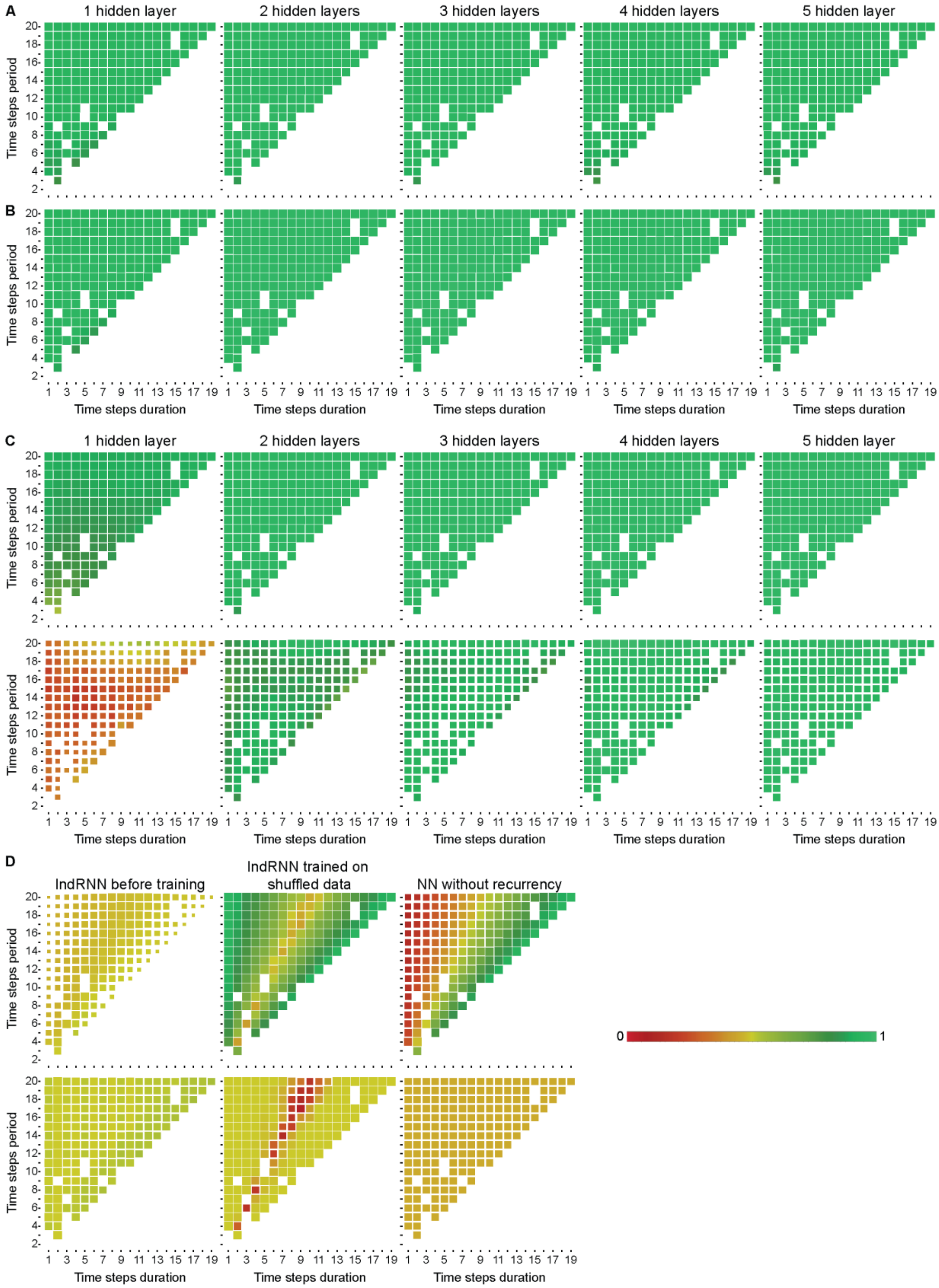


**Supplementary Fig. 3: Consistent per-event target accuracies for the IndRNNs trained on repetitive events, while IndRNNs before training, IndRNNs trained on shuffled data, and the NN without recurrency mimicked accuracy patterns predicted by adhering to specific strategies. (A)** The per-event accuracies of all 50 repetitions of the parameter-matched IndRNNs trained on repetitive events **(B)** The per-event accuracies of the 25 best-performing repetitions of the parameter-matched IndRNNs trained on repetitive events **(C)** The per-event (top) and per-state-change (bottom) accuracies for all 50 repetitions of the layer-size-matched IndRNNs **(D)** The per-event (top) and per-state-change (bottom) accuracies for 5-layer parameter-matched IndRNNs before training, IndRNNs trained on shuffled data, and NNs without recurrency. These network types mimicked patterns produced by random outputs, outputting the proportion of “on” time steps, and outputting only “on” time steps, respectively. The outcomes of these strategies are given in Fig. 2D. Format follows Fig. 2D-F.

| | Median [IQR] | Test statistics | | | | |
|---|---|---|---|---|---|---|
| | | 1 layer | 2 layers | 3 layers | 4 layers | 5 layers |
| **1 layer** | 0.03 [0.03, 0.04] | | <u>0.003</u> | 0.065 | 0.474 | 0.298 |
| **2 layers** | 0.02 [0.01, 0.04] | <u>-3.60</u> | | 0.729 | 0.280 | 0.474 |
| **3 layers** | 0.02 [0.01, 0.04] | -2.68 | 0.93 | | 0.703 | 0.729 |
| **4 layers** | 0.03 [0.02, 0.04] | -1.55 | 2.05 | 1.12 | | 0.729 |
| **5 layers** | 0.02 [0.01, 0.04] | -1.97 | 1.64 | 0.71 | -0.41 | |

**Supplementary Table 5A: Variance between timings lower for the one- than the two-layer parameter-size-matched IndRNNs.** Given are the median [IQR] variance between timings over network repetitions. The test statistics comparing each pair of network depths (post-hoc Dunn's test) are given in black cells, with corresponding corrected *p*-values (Holm-Šidák) of this comparison in the gray cells. Significant comparisons underlined

| | Median [IQR] | Test statistics | | | | |
|---|---|---|---|---|---|---|
| | | 1 layer | 2 layers | 3 layers | 4 layers | 5 layers |
| **1 layer** | 0.03 [0.02, 0.03] | | <u>$3 \times 10^{-7}$</u> | <u>$3 \times 10^{-6}$</u> | <u>0.025</u> | <u>$4 \times 10^{-4}$</u> |
| **2 layers** | 0.01 [0.00, 0.01] | <u>-5.54</u> | | 0.682 | <u>0.049</u> | 0.458 |
| **3 layers** | 0.01 [0.00, 0.01] | <u>-5.13</u> | 0.41 | | 0.124 | 0.566 |
| **4 layers** | 0.02 [0.01, 0.02] | <u>-2.90</u> | <u>2.63</u> | 2.23 | | 0.566 |
| **5 layers** | 0.01 [0.00, 0.02] | <u>-4.07</u> | 1.47 | 1.06 | -1.17 | |

**Supplementary Table 5B: Variance between timings for the top-25 network repetitions of parameter-size-matched IndRNNs are lower for the one-layer network than all mutli-layer networks**. Given are the median [IQR] variance between timings over network repetitions. The test statistics comparing each pair of network depths (post-hoc Dunn's test) are given in black cells, with corresponding corrected *p*-values (Holm-Šidák) of this comparison in the gray cells. Significant comparisons underlined

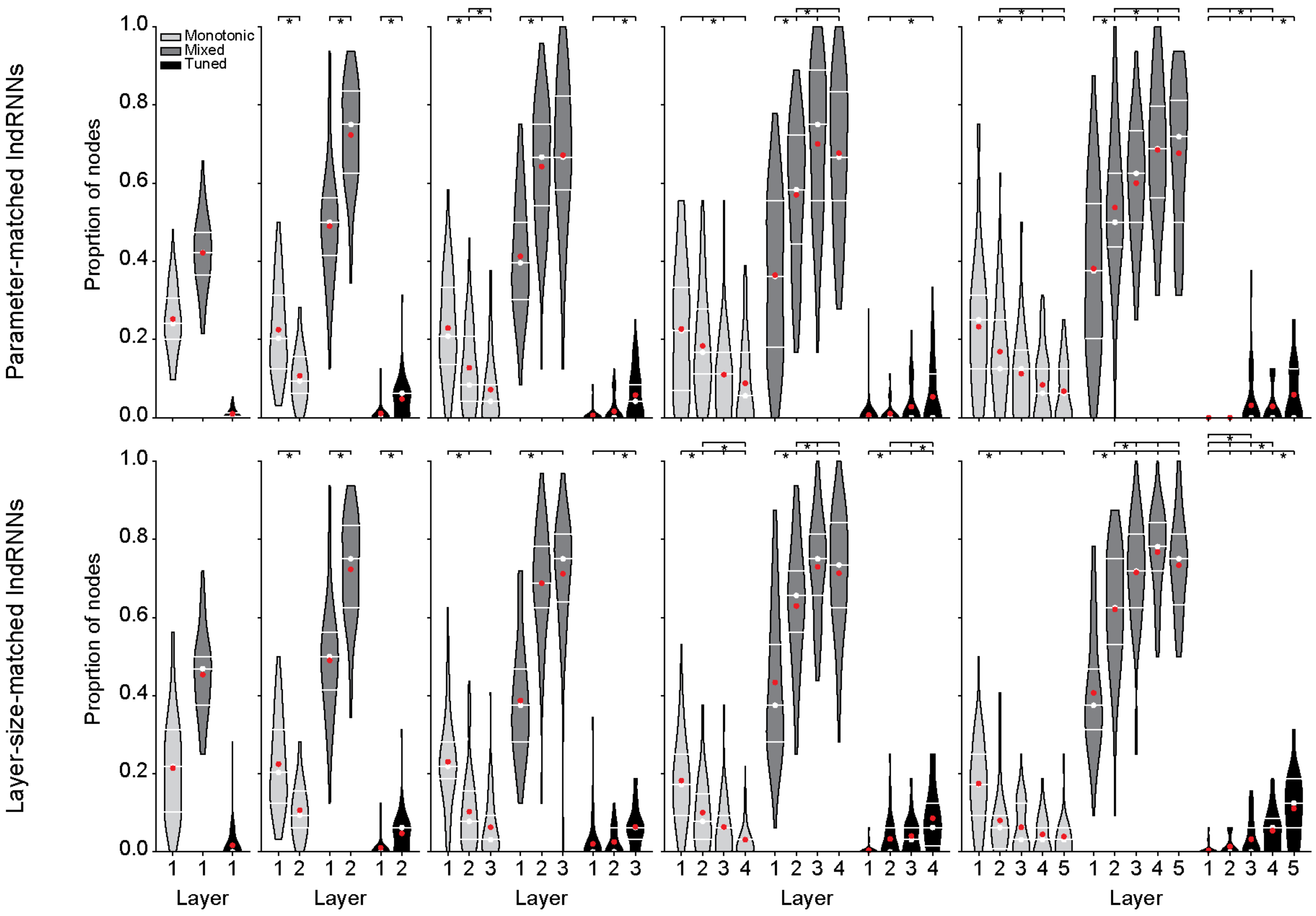


**Supplementary Fig. 4: Progressive transitions from monotonic to mixed and tuned responses across network layers in trained IndRNNs.** Proportions of nodes classified as having monotonic, mixed, or tuned responses in parameter-matched (top) or layer-size-matched (bottom) IndRNNs of different depths Format follows Fig. 2A.

| Network | *H*-statistic | Degrees of freedom | *p*-value | Effect size ($\varepsilon^2$) |
|---|---|---|---|---|
| **Monotonic** | | | | |
| Parameter-matched 2-layer | 27.15 | 1 | $2 \times 10^{-7}$ | 0.26 |
| Parameter-matched 3-layer | 39.31 | 2 | $3 \times 10^{-9}$ | 0.25 |
| Parameter-matched 4-layer | 27.11 | 3 | $6 \times 10^{-6}$ | 0.12 |
| Parameter-matched 5-layer | 44.43 | 4 | $5 \times 10^{-9}$ | 0.16 |
| Layer-size-matched 2-layer | 27.15 | 1 | $2 \times 10^{-7}$ | 0.26 |
| Layer-size-matched 3-layer | 55.36 | 2 | $1 \times 10^{-12}$ | 0.36 |
| Layer-size-matched 4-layer | 63.83 | 3 | $9 \times 10^{-14}$ | 0.31 |
| Layer-size-matched 5-layer | 60.79 | 4 | $2 \times 10^{-12}$ | 0.23 |
| Parameter-matched 5-layer before training | 16.18 | 4 | 0.003 | 0.05 |
| Layer-size-matched 5-layer before training | 25.34 | 4 | $4 \times 10^{-5}$ | 0.09 |
| Parameter-matched 5-layer trained on shuffled data | 39.36 | 4 | $6 \times 10^{-8}$ | 0.14 |
| Layer-size-matched 5-layer trained on shuffled data | 66.36 | 4 | $1 \times 10^{-13}$ | 0.25 |
| Parameter-matched 5-layer without recurrency | 44.94 | 4 | $4 \times 10^{-9}$ | 0.16 |
| Layer-size-matched 5-layer without recurrency | 11.50 | 4 | 0.021 | 0.03 |
| **Mixed** | | | | |
| Parameter-matched 2-layer | 43.90 | 1 | $3 \times 10^{-11}$ | 0.43 |
| Parameter-matched 3-layer | 49.67 | 2 | $2 \times 10^{-11}$ | 0.32 |
| Parameter-matched 4-layer | 57.87 | 3 | $2 \times 10^{-12}$ | 0.28 |
| Parameter-matched 5-layer | 59.52 | 4 | $4 \times 10^{-12}$ | 0.22 |
| Layer-size-matched 2-layer | 43.90 | 1 | $3 \times 10^{-11}$ | 0.43 |
| Layer-size-matched 3-layer | 71.18 | 2 | $4 \times 10^{-16}$ | 0.46 |
| Layer-size-matched 4-layer | 61.71 | 3 | $3 \times 10^{-13}$ | 0.30 |
| Layer-size-matched 5-layer | 101.41 | 4 | $5 \times 10^{-21}$ | 0.39 |
| Parameter-matched 5-layer before training | 3.22 | 4 | 0.521 | -0.00 |
| Layer-size-matched 5-layer before training | 2.20 | 4 | 0.699 | -0.01 |
| Parameter-matched 5-layer trained on shuffled data | 24.23 | 4 | $7 \times 10^{-5}$ | 0.08 |
| Layer-size-matched 5-layer trained on shuffled data | 38.06 | 4 | $1 \times 10^{-7}$ | 0.14 |
| **Tuned** | | | | |
| Parameter-matched 2-layer | 21.90 | 1 | $3 \times 10^{-6}$ | 0.21 |
| Parameter-matched 3-layer | 29.29 | 2 | $4 \times 10^{-7}$ | 0.18 |
| Parameter-matched 4-layer | 19.63 | 3 | $2 \times 10^{-4}$ | 0.08 |
| Parameter-matched 5-layer | 48.41 | 4 | $8 \times 10^{-10}$ | 0.18 |
| Layer-size-matched 2-layer | 21.90 | 1 | $3 \times 10^{-6}$ | 0.21 |
| Layer-size-matched 3-layer | 32.95 | 2 | $7 \times 10^{-8}$ | 0.21 |
| Layer-size-matched 4-layer | 55.43 | 3 | $6 \times 10^{-12}$ | 0.26 |
| Layer-size-matched 5-layer | 94.31 | 4 | $2 \times 10^{-19}$ | 0.36 |
| Parameter-matched 5-layer before training | 1.05 | 4 | 0.902 | -0.01 |
| Layer-size-matched 5-layer before training | 3.09 | 4 | 0.542 | -0.00 |
| Parameter-matched 5-layer trained on shuffled data | 8.33 | 4 | 0.080 | 0.02 |
| Layer-size-matched 5-layer trained on shuffled data | 24.17 | 4 | $7 \times 10^{-5}$ | 0.08 |

**Supplementary Table 6A: Differences in proportion of response types between network layers.** Given are the test statistics of the Kruskal-Wallis tests between network layers. Post-hoc statistics are given in Table 6B-I. Significant differences underlined

| | Median [IQR] | Test statistic | | | | |
|---|---|---|---|---|---|---|
| | | Layer 1 | Layer 2 | Layer 3 | Layer 4 | Layer 5 |
| **Monotonic** | | | | | | |
| **1-layer network** | | | | | | |
| **Layer 1** | 0.24 [0.20, 0.31] | | | | | |
| **2-layer network** | | | | | | |
| **Layer 1** | 0.20 [0.12, 0.31] | | | | | |
| **Layer 2** | 0.09 [0.06, 0.16] | | | | | |
| **3-layer network** | | | | | | |
| **Layer 1** | 0.21 [0.14, 0.33] | | 0.001 | $1 \times 10^{-9}$ | | |
| **Layer 2** | 0.08 [0.04, 0.21] | -3.62 | | 0.009 | | |
| **Layer 3** | 0.04 [0.00, 0.08] | -6.24 | -2.63 | | | |
| **4-layer network** | | | | | | |
| **Layer 1** | 0.22 [0.07, 0.33] | | 0.480 | 0.002 | $3 \times 10^{-5}$ | |
| **Layer 2** | 0.17 [0.11, 0.28] | -0.92 | | 0.033 | 0.001 | |
| **Layer 3** | 0.11 [0.00, 0.17] | -3.46 | -2.54 | | 0.480 | |
| **Layer 4** | 0.06 [0.00, 0.17] | -4.54 | -3.62 | -1.08 | | |
| **5-layer network** | | | | | | |
| **Layer 1** | 0.25 [0.12, 0.31] | | 0.267 | 0.001 | $6 \times 10^{-6}$ | $2 \times 10^{-7}$ |
| **Layer 2** | 0.12 [0.12, 0.25] | -1.65 | | 0.129 | 0.006 | $5 \times 10^{-4}$ |
| **Layer 3** | 0.12 [0.00, 0.17] | -3.86 | -2.21 | | 0.471 | 0.258 |
| **Layer 4** | 0.06 [0.00, 0.12] | -4.96 | -3.30 | -1.10 | | 0.482 |
| **Layer 5** | 0.06 [0.00, 0.12] | -5.66 | -4.01 | -1.80 | -0.70 | |
| **Mixed** | | | | | | |
| **1-layer network** | | | | | | |
| **Layer 1** | 0.42 [0.37, 0.47] | | | | | |
| **2-layer network** | | | | | | |
| **Layer 1** | 0.50 [0.41, 0.56] | | | | | |
| **Layer 2** | 0.75 [0.62, 0.84] | | | | | |
| **3-layer network** | | | | | | |
| **Layer 1** | 0.40 [0.30, 0.50] | | $2 \times 10^{-8}$ | $4 \times 10^{-10}$ | | |
| **Layer 2** | 0.67 [0.54, 0.75] | 5.75 | | 0.510 | | |
| **Layer 3** | 0.67 [0.58, 0.82] | 6.41 | 0.66 | | | |
| **4-layer network** | | | | | | |
| **Layer 1** | 0.36 [0.18, 0.56] | | $3 \times 10^{-4}$ | $5 \times 10^{-11}$ | $2 \times 10^{-9}$ | |
| **Layer 2** | 0.58 [0.44, 0.72] | 3.97 | | 0.012 | 0.043 | |
| **Layer 3** | 0.75 [0.56, 0.89] | 6.84 | 2.87 | | 0.564 | |
| **Layer 4** | 0.67 [0.56, 0.83] | 6.27 | 2.30 | -0.58 | | |
| **5-layer network** | | | | | | |
| **Layer 1** | 0.38 [0.20, 0.55] | | 0.015 | $4 \times 10^{-5}$ | $5 \times 10^{-10}$ | $1 \times 10^{-9}$ |
| **Layer 2** | 0.50 [0.44, 0.62] | 2.98 | | 0.221 | 0.002 | 0.003 |
| **Layer 3** | 0.62 [0.50, 0.73] | 4.54 | 1.57 | | 0.163 | 0.165 |
| **Layer 4** | 0.69 [0.56, 0.80] | 6.56 | 3.58 | 2.02 | | 0.900 |
| **Layer 5** | 0.72 [0.50, 0.81] | 6.43 | 3.46 | 1.89 | -0.13 | |
| **Tuned** | | | | | | |
| **1-layer network** | | | | | | |
| **Layer 1** | 0.00 [0.00, 0.01] | | | | | |
| **2-layer network** | | | | | | |
| **Layer 1** | 0.00 [0.00, 0.00] | | | | | |
| **Layer 2** | 0.06 [0.00, 0.06] | | | | | |
| **3-layer network** | | | | | | |
| **Layer 1** | 0.00 [0.00, 0.00] | | 0.197 | $6 \times 10^{-7}$ | | |
| **Layer 2** | 0.00 [0.00, 0.00] | 1.29 | | $2 \times 10^{-4}$ | | |
| **Layer 3** | 0.04 [0.00, 0.08] | 5.20 | 3.91 | | | |
| **4-layer network** | | | | | | |
| **Layer 1** | 0.00 [0.00, 0.00] | | 0.289 | 0.051 | $2 \times 10^{-4}$ | |
| **Layer 2** | 0.00 [0.00, 0.00] | 1.06 | | 0.285 | 0.009 | |
| **Layer 3** | 0.00 [0.00, 0.00] | 2.48 | 1.42 | | 0.250 | |
| **Layer 4** | 0.00 [0.00, 0.11] | 4.17 | 3.11 | 1.69 | | |
| **5-layer network** | | | | | | |
| **Layer 1** | 0.00 [0.00, 0.00] | | 1.000 | 0.021 | 0.017 | $4 \times 10^{-8}$ |
| **Layer 2** | 0.00 [0.00, 0.00] | -0.00 | | 0.021 | 0.017 | $4 \times 10^{-8}$ |
| **Layer 3** | 0.00 [0.00, 0.00] | 2.86 | 2.86 | | 0.971 | 0.017 |
| **Layer 4** | 0.00 [0.00, 0.00] | 3.07 | 3.07 | 0.22 | | 0.021 |
| **Layer 5** | 0.00 [0.00, 0.12] | 5.87 | 5.87 | 3.01 | 2.80 | |

**Supplementary Table 6B: Gradual transition from monotonic to mixed and tuned responses over parameter-matched IndRNN layers.** Given are the median [IQR] proportion of classified nodes over network repetitions. The test statistics comparing each pair of network depths (post-hoc Dunn's test) are given in black cells, with corresponding corrected *p*-values (Holm-Šidák) of this comparison in the gray cells. Significant comparisons underlined

| | Median [IQR] | Test statistic | | | | |
|---|---|---|---|---|---|---|
| | | **Layer 1** | **Layer 2** | **Layer 3** | **Layer 4** | **Layer 5** |
| | | | **Monotonic** | | | |
| **1-layer network** | | | | | | |
| **Layer 1** | 0.22 [0.10, 0.31] | | | | | |
| **2-layer network** | | | | | | |
| **Layer 1** | 0.20 [0.12, 0.31] | | | | | |
| **Layer 2** | 0.09 [0.06, 0.16] | | | | | |
| **3-layer network** | | | | | | |
| **Layer 1** | 0.22 [0.19, 0.28] | | $3 \times 10^{-7}$ | $2 \times 10^{-12}$ | | |
| **Layer 2** | 0.08 [0.03, 0.16] | -5.25 | | 0.052 | | |
| **Layer 3** | 0.03 [0.00, 0.09] | -7.19 | -1.94 | | | |
| **4-layer network** | | | | | | |
| **Layer 1** | 0.17 [0.09, 0.25] | | 0.002 | $2 \times 10^{-7}$ | $1 \times 10^{-13}$ | |
| **Layer 2** | 0.08 [0.03, 0.15] | -3.38 | | 0.065 | $8 \times 10^{-5}$ | |
| **Layer 3** | 0.06 [0.00, 0.09] | -5.52 | -2.13 | | 0.065 | |
| **Layer 4** | 0.03 [0.00, 0.03] | -7.65 | -4.27 | -2.13 | | |
| **5-layer network** | | | | | | |
| **Layer 1** | 0.17 [0.09, 0.25] | | $7 \times 10^{-5}$ | $8 \times 10^{-7}$ | $2 \times 10^{-9}$ | $3 \times 10^{-11}$ |
| **Layer 2** | 0.06 [0.01, 0.09] | -4.41 | | 0.669 | 0.238 | 0.060 |
| **Layer 3** | 0.03 [0.00, 0.12] | -5.32 | -0.92 | | 0.669 | 0.339 |
| **Layer 4** | 0.03 [0.00, 0.06] | -6.34 | -1.94 | -1.02 | | 0.669 |
| **Layer 5** | 0.03 [0.00, 0.06] | -6.98 | -2.57 | -1.65 | -0.63 | |
| | | | **Mixed** | | | |
| **1-layer network** | | | | | | |
| **Layer 1** | 0.47 [0.38, 0.50] | | | | | |
| **2-layer network** | | | | | | |
| **Layer 1** | 0.50 [0.41, 0.56] | | | | | |
| **Layer 2** | 0.75 [0.62, 0.84] | | | | | |
| **3-layer network** | | | | | | |
| **Layer 1** | 0.38 [0.28, 0.47] | | $2 \times 10^{-11}$ | $4 \times 10^{-14}$ | | |
| **Layer 2** | 0.69 [0.62, 0.78] | 6.82 | | 0.376 | | |
| **Layer 3** | 0.75 [0.64, 0.81] | 7.71 | 0.89 | | | |
| **4-layer network** | | | | | | |
| **Layer 1** | 0.38 [0.28, 0.53] | | $1 \times 10^{-4}$ | $2 \times 10^{-11}$ | $3 \times 10^{-10}$ | |
| **Layer 2** | 0.66 [0.56, 0.72] | 4.20 | | 0.015 | 0.035 | |
| **Layer 3** | 0.75 [0.66, 0.81] | 7.00 | 2.80 | | 0.668 | |
| **Layer 4** | 0.73 [0.62, 0.84] | 6.57 | 2.37 | -0.43 | | |
| **5-layer network** | | | | | | |
| **Layer 1** | 0.38 [0.31, 0.47] | | $3 \times 10^{-5}$ | $4 \times 10^{-12}$ | $2 \times 10^{-18}$ | $2 \times 10^{-13}$ |
| **Layer 2** | 0.62 [0.53, 0.75] | 4.62 | | 0.036 | $7 \times 10^{-5}$ | 0.013 |
| **Layer 3** | 0.72 [0.62, 0.81] | 7.23 | 2.61 | | 0.206 | 0.697 |
| **Layer 4** | 0.78 [0.72, 0.84] | 9.02 | 4.40 | 1.79 | | 0.298 |
| **Layer 5** | 0.75 [0.63, 0.81] | 7.62 | 3.00 | 0.39 | -1.40 | |
| | | | **Tuned** | | | |
| **1-layer network** | | | | | | |
| **Layer 1** | 0.00 [0.00, 0.00] | | | | | |
| **2-layer network** | | | | | | |
| **Layer 1** | 0.00 [0.00, 0.00] | | | | | |
| **Layer 2** | 0.06 [0.00, 0.06] | | | | | |
| **3-layer network** | | | | | | |
| **Layer 1** | 0.00 [0.00, 0.00] | | 0.119 | $8 \times 10^{-8}$ | | |
| **Layer 2** | 0.00 [0.00, 0.06] | 1.56 | | $1 \times 10^{-4}$ | | |
| **Layer 3** | 0.06 [0.03, 0.06] | 5.56 | 4.01 | | | |
| **4-layer network** | | | | | | |
| **Layer 1** | 0.00 [0.00, 0.00] | | 0.004 | $7 \times 10^{-5}$ | $1 \times 10^{-12}$ | |
| **Layer 2** | 0.00 [0.00, 0.06] | 3.18 | | 0.240 | $1 \times 10^{-4}$ | |
| **Layer 3** | 0.03 [0.00, 0.06] | 4.35 | 1.17 | | 0.005 | |
| **Layer 4** | 0.06 [0.02, 0.12] | 7.35 | 4.17 | 3.00 | | |
| **5-layer network** | | | | | | |
| **Layer 1** | 0.00 [0.00, 0.00] | | 0.213 | 0.012 | $3 \times 10^{-7}$ | $1 \times 10^{-16}$ |
| **Layer 2** | 0.00 [0.00, 0.00] | 1.25 | | 0.160 | $2 \times 10^{-4}$ | $2 \times 10^{-12}$ |
| **Layer 3** | 0.00 [0.00, 0.06] | 2.98 | 1.73 | | 0.039 | $2 \times 10^{-7}$ |
| **Layer 4** | 0.06 [0.00, 0.09] | 5.45 | 4.21 | 2.48 | | 0.009 |
| **Layer 5** | 0.12 [0.06, 0.19] | 8.58 | 7.33 | 5.60 | 3.13 | |

**Supplementary Table 6C: Gradual transition from monotonic to mixed and tuned responses over layer-size-matched IndRNN layers.** Like Supplementary Table 6A, but for layer-size-matched networks

| | Median [IQR] | Test statistic | | | | |
|---|---|---|---|---|---|---|
| | | Layer 1 | Layer 2 | Layer 3 | Layer 4 | Layer 5 |
| | | **Monotonic** | | | | |
| **Layer 1** | 0.25 [0.12, 0.30] | | 0.050 | 0.007 | $4 \times 10^{-8}$ | $3 \times 10^{-5}$ |
| **Layer 2** | 0.12 [0.06, 0.23] | -2.63 | | 0.484 | 0.009 | 0.172 |
| **Layer 3** | 0.12 [0.02, 0.25] | -3.33 | -0.70 | | 0.055 | 0.480 |
| **Layer 4** | 0.06 [0.00, 0.12] | -5.87 | -3.23 | -2.53 | | 0.480 |
| **Layer 5** | 0.06 [0.00, 0.17] | -4.63 | -1.99 | -1.29 | 1.24 | |
| | | **Mixed** | | | | |
| **Layer 1** | 0.38 [0.25, 0.50] | | 0.293 | 0.004 | $2 \times 10^{-4}$ | 0.001 |
| **Layer 2** | 0.44 [0.38, 0.62] | 1.89 | | 0.372 | 0.130 | 0.293 |
| **Layer 3** | 0.53 [0.44, 0.62] | 3.49 | 1.60 | | 0.845 | 0.893 |
| **Layer 4** | 0.62 [0.44, 0.69] | 4.23 | 2.33 | 0.73 | | 0.893 |
| **Layer 5** | 0.56 [0.39, 0.62] | 3.80 | 1.91 | 0.31 | -0.42 | |
| | | **Tuned** | | | | |
| **Layer 1** | 0.00 [0.00, 0.00] | Kruskal-Wallis not significant | | | | |
| **Layer 2** | 0.00 [0.00, 0.00] | | | | | |
| **Layer 3** | 0.00 [0.00, 0.00] | | | | | |
| **Layer 4** | 0.00 [0.00, 0.00] | | | | | |
| **Layer 5** | 0.00 [0.00, 0.00] | | | | | |

**Supplementary Table 6D: Gradual transition from monotonic to mixed and tuned responses over IndRNN layers of parameter-matched IndRNNs trained on shuffled data.** Like Supplementary Table 6A, but for the parameter-matched five-layer IndRNN

| | Median [IQR] | Test statistic | | | | |
|---|---|---|---|---|---|---|
| | | Layer 1 | Layer 2 | Layer 3 | Layer 4 | Layer 5 |
| | | **Monotonic** | | | | |
| **Layer 1** | 0.19 [0.09, 0.25] | | 0.017 | $6 \times 10^{-5}$ | $9 \times 10^{-10}$ | $9 \times 10^{-12}$ |
| **Layer 2** | 0.09 [0.06, 0.18] | -2.92 | | 0.226 | 0.002 | $2 \times 10^{-4}$ |
| **Layer 3** | 0.06 [0.03, 0.12] | -4.48 | -1.55 | | 0.131 | 0.030 |
| **Layer 4** | 0.06 [0.00, 0.06] | -6.47 | -3.55 | -2.00 | | 0.504 |
| **Layer 5** | 0.03 [0.00, 0.06] | -7.14 | -4.22 | -2.67 | -0.67 | |
| | | **Mixed** | | | | |
| **Layer 1** | 0.38 [0.28, 0.47] | | $2 \times 10^{-5}$ | $3 \times 10^{-7}$ | $3 \times 10^{-5}$ | $4 \times 10^{-4}$ |
| **Layer 2** | 0.56 [0.38, 0.69] | 4.76 | | 0.908 | 0.908 | 0.908 |
| **Layer 3** | 0.58 [0.45, 0.69] | 5.52 | 0.76 | | 0.897 | 0.598 |
| **Layer 4** | 0.55 [0.44, 0.62] | 4.61 | -0.15 | -0.90 | | 0.908 |
| **Layer 5** | 0.50 [0.41, 0.59] | 4.04 | -0.71 | -1.47 | -0.57 | |
| | | **Tuned** | | | | |
| **Layer 1** | 0.00 [0.00, 0.00] | | 0.969 | 0.035 | 0.001 | 0.023 |
| **Layer 2** | 0.00 [0.00, 0.00] | 0.22 | | 0.048 | 0.003 | 0.035 |
| **Layer 3** | 0.02 [0.00, 0.06] | 2.80 | 2.58 | | 0.769 | 0.969 |
| **Layer 4** | 0.03 [0.00, 0.06] | 3.83 | 3.60 | 1.02 | | 0.779 |
| **Layer 5** | 0.00 [0.00, 0.09] | 2.98 | 2.75 | 0.17 | -0.85 | |

**Supplementary Table 6E: Gradual transition from monotonic to mixed and tuned responses over IndRNN layers of layer-size-matched IndRNNs trained on shuffled data.** Like Supplementary Table 6C, but for the layer-size-matched IndRNN

| | Median [IQR] | Test statistic | | | | |
|---|---|---|---|---|---|---|
| | | Layer 1 | Layer 2 | Layer 3 | Layer 4 | Layer 5 |
| | | **Monotonic** | | | | |
| **Layer 1** | 0.25 [0.14, 0.38] | | 0.068 | 0.001 | 0.108 | 0.422 |
| **Layer 2** | 0.38 [0.25, 0.48] | 2.66 | | 0.651 | 0.835 | 0.714 |
| **Layer 3** | 0.38 [0.31, 0.50] | 3.86 | 1.20 | | 0.582 | 0.201 |
| **Layer 4** | 0.38 [0.25, 0.48] | 2.45 | -0.21 | -1.40 | | 0.714 |
| **Layer 5** | 0.31 [0.25, 0.44] | 1.71 | -0.95 | -2.15 | -0.74 | |
| | | **Mixed** | | | | |
| **Layer 1** | 0.12 [0.08, 0.42] | Kruskal-Wallis not significant | | | | |
| **Layer 2** | 0.19 [0.08, 0.48] | | | | | |
| **Layer 3** | 0.12 [0.06, 0.31] | | | | | |
| **Layer 4** | 0.12 [0.06, 0.30] | | | | | |
| **Layer 5** | 0.19 [0.06, 0.31] | | | | | |
| | | **Tuned** | | | | |
| **Layer 1** | 0.00 [0.00, 0.00] | Kruskal-Wallis not significant | | | | |
| **Layer 2** | 0.00 [0.00, 0.00] | | | | | |
| **Layer 3** | 0.00 [0.00, 0.00] | | | | | |
| **Layer 4** | 0.00 [0.00, 0.00] | | | | | |
| **Layer 5** | 0.00 [0.00, 0.00] | | | | | |

**Supplementary Table 6F: No gradual transition from monotonic to mixed and tuned responses over IndRNN layers for the parameter-matched IndRNNs before training.** Like Supplementary Table 6A, but for the parameter-matched five-layer IndRNN.

| | Median [IQR] | Test statistic | | | | |
|---|---|---|---|---|---|---|
| | | Layer 1 | Layer 2 | Layer 3 | Layer 4 | Layer 5 |
| **Monotonic** | | | | | | |
| **Layer 1** | 0.25 [0.19, 0.31] | | $3 \times 10^{-5}$ | 0.002 | 0.008 | 0.080 |
| **Layer 2** | 0.41 [0.31, 0.44] | 4.69 | | 0.725 | 0.594 | 0.165 |
| **Layer 3** | 0.34 [0.26, 0.47] | 3.76 | -0.94 | | 0.725 | 0.620 |
| **Layer 4** | 0.34 [0.25, 0.43] | 3.31 | -1.39 | -0.45 | | 0.725 |
| **Layer 5** | 0.31 [0.22, 0.44] | 2.52 | -2.18 | -1.24 | -0.79 | |
| **Mixed** | | | | | | |
| **Layer 1** | 0.19 [0.06, 0.34] | Kruskal-Wallis not significant | | | | |
| **Layer 2** | 0.20 [0.09, 0.46] | | | | | |
| **Layer 3** | 0.30 [0.09, 0.46] | | | | | |
| **Layer 4** | 0.28 [0.14, 0.41] | | | | | |
| **Layer 5** | 0.25 [0.12, 0.43] | | | | | |
| **Tuned** | | | | | | |
| **Layer 1** | 0.00 [0.00, 0.00] | Kruskal-Wallis not significant | | | | |
| **Layer 2** | 0.00 [0.00, 0.00] | | | | | |
| **Layer 3** | 0.00 [0.00, 0.00] | | | | | |
| **Layer 4** | 0.00 [0.00, 0.00] | | | | | |
| **Layer 5** | 0.00 [0.00, 0.00] | | | | | |

**Supplementary Table 6G: No gradual transition from monotonic to mixed and tuned responses over IndRNN layers for the layer-size-matched IndRNNs before training.** Like Supplementary Table 6E, but for the layer-size-matched IndRNN before training.

| | Median [IQR] | Test statistic | | | | |
|---|---|---|---|---|---|---|
| | | Layer 1 | Layer 2 | Layer 3 | Layer 4 | Layer 5 |
| **Monotonic** | | | | | | |
| **Layer 1** | 0.11 [0.11, 0.22] | | 0.179 | $2 \times 10^{-4}$ | 0.087 | $5 \times 10^{-9}$ |
| **Layer 2** | 0.11 [0.00, 0.15] | -1.98 | | 0.096 | 0.648 | $2 \times 10^{-4}$ |
| **Layer 3** | 0.00 [0.00, 0.11] | -4.31 | -2.33 | | 0.179 | 0.179 |
| **Layer 4** | 0.11 [0.00, 0.11] | -2.43 | -0.46 | 1.87 | | 0.001 |
| **Layer 5** | 0.00 [0.00, 0.00] | -6.23 | -4.25 | -1.92 | -3.79 | |
| **Mixed** | | | | | | |
| **Layer 1** | 0.00 [0.00, 0.00] | Kruskal-Wallis not significant | | | | |
| **Layer 2** | 0.00 [0.00, 0.00] | | | | | |
| **Layer 3** | 0.00 [0.00, 0.00] | | | | | |
| **Layer 4** | 0.00 [0.00, 0.00] | | | | | |
| **Layer 5** | 0.00 [0.00, 0.00] | | | | | |
| **Tuned** | | | | | | |
| **Layer 1** | 0.00 [0.00, 0.00] | Kruskal-Wallis not significant | | | | |
| **Layer 2** | 0.00 [0.00, 0.00] | | | | | |
| **Layer 3** | 0.00 [0.00, 0.00] | | | | | |
| **Layer 4** | 0.00 [0.00, 0.00] | | | | | |
| **Layer 5** | 0.00 [0.00, 0.00] | | | | | |

**Supplementary Table 6H: Predominantly monotonic responses in parameter-matched neural networks without recurrency.** Given are the median [IQR] proportion of classified nodes over network repetitions. The test statistics comparing each pair of network depths (post-hoc Dunn's test) are given in black cells, with corresponding corrected *p*-values (Holm-Šidák) of this comparison in the gray cells. Significant comparisons underlined.

| | Median [IQR] | Test statistic | | | | |
|---|---|---|---|---|---|---|
| | | **Layer 1** | **Layer 2** | **Layer 3** | **Layer 4** | **Layer 5** |
| **Monotonic** | | | | | | |
| **Layer 1** | 0.06 [0.01, 0.12] | | 0.753 | 0.252 | 0.560 | 0.012 |
| **Layer 2** | 0.06 [0.00, 0.06] | -1.17 | | 0.753 | 0.781 | 0.265 |
| **Layer 3** | 0.06 [0.00, 0.06] | -2.15 | -0.98 | | 0.781 | 0.753 |
| **Layer 4** | 0.06 [0.00, 0.06] | -1.52 | -0.36 | 0.63 | | 0.465 |
| **Layer 5** | 0.00 [0.00, 0.06] | -3.24 | -2.08 | -1.09 | -1.72 | |
| **Mixed** | | | | | | |
| **Layer 1** | 0.00 [0.00, 0.00] | Kruskal-Wallis not significant | | | | |
| **Layer 2** | 0.00 [0.00, 0.00] | | | | | |
| **Layer 3** | 0.00 [0.00, 0.00] | | | | | |
| **Layer 4** | 0.00 [0.00, 0.00] | | | | | |
| **Layer 5** | 0.00 [0.00, 0.00] | | | | | |
| **Tuned** | | | | | | |
| **Layer 1** | 0.00 [0.00, 0.00] | Kruskal-Wallis not significant | | | | |
| **Layer 2** | 0.00 [0.00, 0.00] | | | | | |
| **Layer 3** | 0.00 [0.00, 0.00] | | | | | |
| **Layer 4** | 0.00 [0.00, 0.00] | | | | | |
| **Layer 5** | 0.00 [0.00, 0.00] | | | | | |

**Supplementary Table 6I: Predominantly monotonic responses in layer-size-matched neural networks without recurrency.** Given are the median [IQR] proportion of classified nodes over network repetitions. The test statistics comparing each pair of network depths (post-hoc Dunn's test) are given in black cells, with corresponding corrected *p*-values (Holm-Šidák) of this comparison in the gray cells. Significant comparisons underlined.

| | Median [IQR] | Test statistic | | | | |
|---|---|---|---|---|---|---|
| | | Layer 1 | Layer 2 | Layer 3 | Layer 4 | Layer 5 |
| **Monotonic** | | | | | | |
| 1 layers | 0.24 [0.20, 0.31] | | $2 \times 10^{-7}$ | $2 \times 10^{-14}$ | $1 \times 10^{-12}$ | $2 \times 10^{-14}$ |
| 2 layers | 0.09 [0.06, 0.16] | -5.58 | | 0.104 | 0.265 | 0.104 |
| 3 layers | 0.04 [0.00, 0.08] | -7.94 | -2.36 | | 0.917 | 0.975 |
| 4 layers | 0.06 [0.00, 0.17] | -7.36 | -1.79 | 0.58 | | 0.917 |
| 5 layers | 0.06 [0.00, 0.12] | -7.91 | -2.33 | 0.03 | -0.55 | |
| **Mixed** | | | | | | |
| 1 layers | 0.42 [0.37, 0.47] | | $3 \times 10^{-13}$ | $1 \times 10^{-9}$ | $1 \times 10^{-9}$ | $1 \times 10^{-9}$ |
| 2 layers | 0.75 [0.62, 0.84] | 7.60 | | 0.793 | 0.793 | 0.793 |
| 3 layers | 0.67 [0.58, 0.82] | 6.41 | -1.20 | | 1.000 | 1.000 |
| 4 layers | 0.67 [0.56, 0.83] | 6.44 | -1.16 | 0.04 | | 1.000 |
| 5 layers | 0.72 [0.50, 0.81] | 6.45 | -1.15 | 0.05 | 0.01 | |
| **Tuned** | | | | | | |
| 1 layers | 0.00 [0.00, 0.01] | | 0.084 | 0.060 | 0.706 | 0.164 |
| 2 layers | 0.06 [0.00, 0.06] | 2.59 | | 0.957 | 0.688 | 0.957 |
| 3 layers | 0.04 [0.00, 0.08] | 2.74 | 0.15 | | 0.629 | 0.957 |
| 4 layers | 0.00 [0.00, 0.11] | 1.23 | -1.35 | -1.51 | | 0.749 |
| 5 layers | 0.00 [0.00, 0.12] | 2.29 | -0.30 | -0.45 | 1.05 | |

**Supplementary Table 7A: More monotonic and less mixed and tuned responses in the last layer of the 1-layer parameter-matched IndRNN.** Given are the median [IQR] proportion of classified nodes over network repetitions. The test statistics comparing each pair of network depths (post-hoc Dunn's test) are given in black cells, with corresponding corrected *p*-values (Holm-Šidák) of this comparison in the gray cells. Significant comparisons underlined

| | Median [IQR] | Test statistic | | | | |
|---|---|---|---|---|---|---|
| | | Layer 1 | Layer 2 | Layer 3 | Layer 4 | Layer 5 |
| **Monotonic** | | | | | | |
| 1 layers | 0.22 [0.10, 0.31] | | 0.006 | $4 \times 10^{-10}$ | $2 \times 10^{-16}$ | $5 \times 10^{-14}$ |
| 2 layers | 0.09 [0.06, 0.16] | -3.17 | | 0.004 | $7 \times 10^{-7}$ | $2 \times 10^{-5}$ |
| 3 layers | 0.03 [0.00, 0.09] | -6.56 | -3.38 | | 0.149 | 0.373 |
| 4 layers | 0.03 [0.00, 0.03] | -8.50 | -5.32 | -1.94 | | 0.496 |
| 5 layers | 0.03 [0.00, 0.06] | -7.82 | -4.64 | -1.26 | 0.68 | |
| **Mixed** | | | | | | |
| 1 layers | 0.47 [0.38, 0.50] | | $5 \times 10^{-13}$ | $1 \times 10^{-12}$ | $3 \times 10^{-12}$ | $2 \times 10^{-13}$ |
| 2 layers | 0.75 [0.62, 0.84] | 7.51 | | 1.000 | 1.000 | 1.000 |
| 3 layers | 0.75 [0.64, 0.81] | 7.40 | -0.12 | | 1.000 | 1.000 |
| 4 layers | 0.73 [0.62, 0.84] | 7.26 | -0.25 | -0.14 | | 0.999 |
| 5 layers | 0.75 [0.63, 0.81] | 7.67 | 0.16 | 0.27 | 0.41 | |
| **Tuned** | | | | | | |
| 1 layers | 0.00 [0.00, 0.00] | | 0.009 | $2 \times 10^{-5}$ | $6 \times 10^{-8}$ | $8 \times 10^{-13}$ |
| 2 layers | 0.06 [0.00, 0.06] | 3.17 | | 0.260 | 0.034 | $1 \times 10^{-4}$ |
| 3 layers | 0.06 [0.03, 0.06] | 4.72 | 1.56 | | 0.283 | 0.030 |
| 4 layers | 0.06 [0.02, 0.12] | 5.80 | 2.63 | 1.07 | | 0.260 |
| 5 layers | 0.12 [0.06, 0.19] | 7.46 | 4.30 | 2.74 | 1.67 | |

**Supplementary Table 7B: Decreasing proportions of monotonic and increasing proportions of tuned responses in the last layer with increasing network complexity.** Given are the median [IQR] proportion of classified nodes over network repetitions of the layer-size-matched IndRNNs. The test statistics comparing each pair of network depths (post-hoc Dunn's test) are given in black cells, with corresponding corrected *p*-values (Holm-Šidák) of this comparison in the gray cells. Significant comparisons underlined

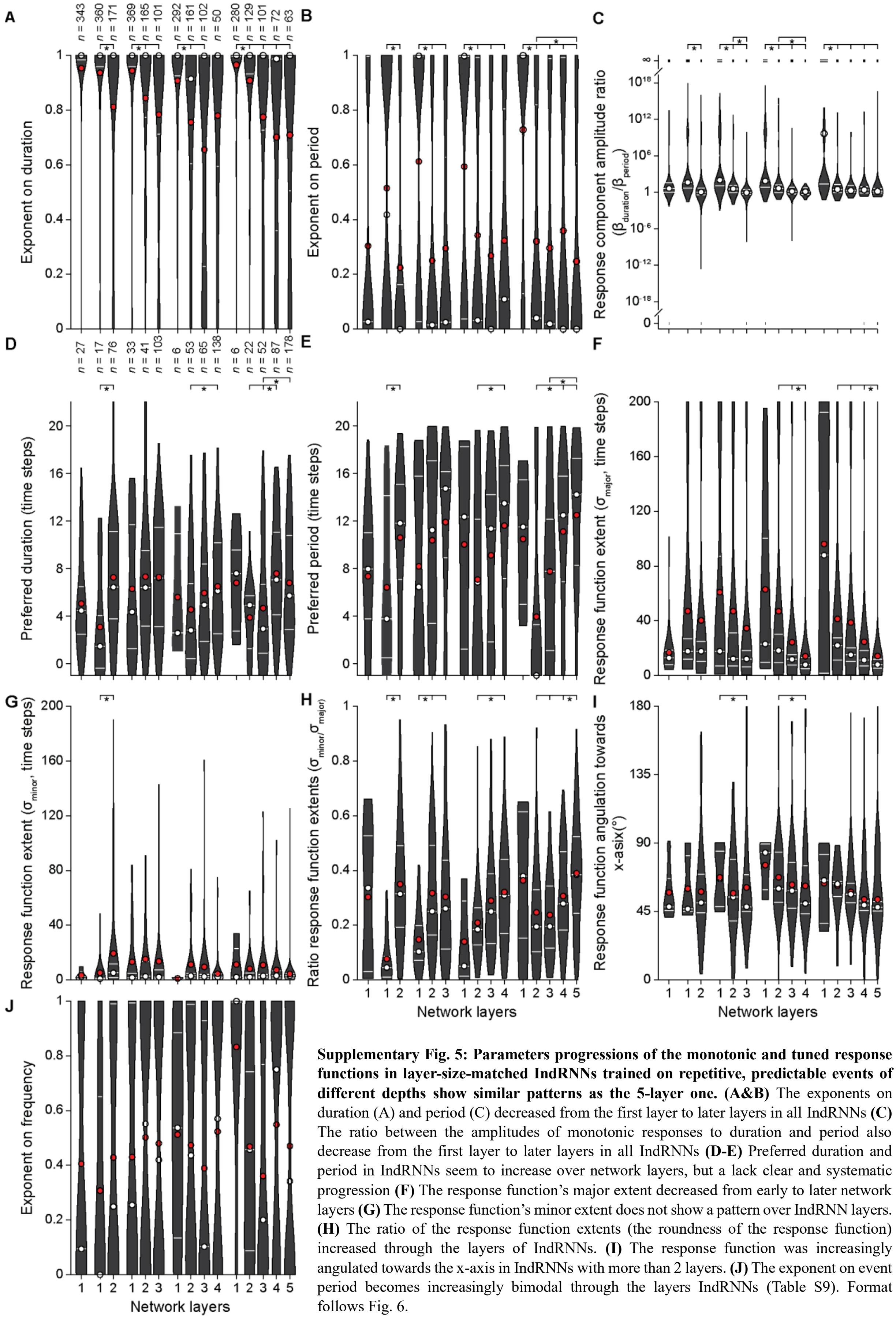

**Supplementary Fig. 5: Parameters progressions of the monotonic and tuned response functions in layer-size-matched IndRNNs trained on repetitive, predictable events of different depths show similar patterns as the 5-layer one. (A&B)** The exponents on duration (A) and period (C) decreased from the first layer to later layers in all IndRNNs **(C)** The ratio between the amplitudes of monotonic responses to duration and period also decrease from the first layer to later layers in all IndRNNs **(D-E)** Preferred duration and period in IndRNNs seem to increase over network layers, but a lack clear and systematic progression **(F)** The response function's major extent decreased from early to later network layers **(G)** The response function's minor extent does not show a pattern over IndRNN layers. **(H)** The ratio of the response function extents (the roundness of the response function) increased through the layers of IndRNNs. **(I)** The response function was increasingly angulated towards the x-axis in IndRNNs with more than 2 layers. **(J)** The exponent on event period becomes increasingly bimodal through the layers IndRNNs (Table S9). Format follows Fig. 6.

| | Median [IQR] | Layer 1 | Layer 2 | Layer 3 | Layer 4 | Layer 5 |
|---|---|---|---|---|---|---|
| **Monotonic: exponent on duration** | | | | | | |
| **Layer 1** | 1.00 [1.00, 1.00] | | $4 \times 10^{-6}$ | $9 \times 10^{-10}$ | $9 \times 10^{-8}$ | $3 \times 10^{-6}$ |
| **Layer 2** | 1.00 [0.93, 1.00] | -5.01 | | 0.460 | 0.490 | 0.671 |
| **Layer 3** | 1.00 [0.73, 1.00] | -6.49 | -1.66 | | 0.991 | 0.991 |
| **Layer 4** | 0.99 [0.36, 1.00] | -5.74 | -1.53 | -0.03 | | 0.991 |
| **Layer 5** | 1.00 [0.51, 1.00] | -5.11 | -1.17 | 0.25 | 0.26 | |
| **Monotonic: exponent on period** | | | | | | |
| **Layer 1** | 1.00 [0.13, 1.00] | | $1 \times 10^{-12}$ | $1 \times 10^{-15}$ | $6 \times 10^{-14}$ | $3 \times 10^{-17}$ |
| **Layer 2** | 0.04 [0.00, 0.82] | -7.35 | | 0.447 | 0.392 | 0.030 |
| **Layer 3** | 0.02 [0.00, 0.99] | -8.28 | -1.34 | | 0.663 | 0.392 |
| **Layer 4** | 0.00 [0.00, 0.99] | -7.78 | -1.67 | -0.44 | | 0.489 |
| **Layer 5** | 0.00 [0.00, 0.61] | -8.70 | -2.80 | -1.57 | -1.07 | |
| **Monotonic: ratio between duration and period component** | | | | | | |
| **Layer 1** | $5 \times 10^{9}$ [23.32, Inf] | | $6 \times 10^{-21}$ | $6 \times 10^{-19}$ | $2 \times 10^{-15}$ | $1 \times 10^{-18}$ |
| **Layer 2** | 3.06 [0.89, 19.84] | -9.63 | | 0.969 | 0.969 | 0.562 |
| **Layer 3** | 1.93 [1.01, 13.5] | -9.13 | -0.26 | | 0.969 | 0.703 |
| **Layer 4** | 2.54 [0.76, 8.19] | -8.20 | -0.40 | -0.16 | | 0.776 |
| **Layer 5** | 1.43 [0.69, 4.31] | -9.02 | -1.52 | -1.24 | -1.01 | |
| **Tuned: preferred duration** | | | | | | |
| **Layer 1** | 0.38 [0.14, 0.48] | | 0.658 | 0.658 | 0.983 | 0.983 |
| **Layer 2** | 0.25 [0.06, 0.28] | -1.30 | | 0.983 | 0.019 | 0.099 |
| **Layer 3** | 0.15 [0.04, 0.33] | -1.20 | 0.33 | | 0.002 | 0.024 |
| **Layer 4** | 0.35 [0.20, 0.55] | 0.32 | 3.08 | 3.71 | | 0.649 |
| **Layer 5** | 0.29 [0.14, 0.54] | -0.12 | 2.44 | 2.96 | -1.40 | |
| **Tuned: preferred period** | | | | | | |
| **Layer 1** | 0.57 [0.25, 0.77] | | 0.546 | 0.798 | 0.798 | 0.798 |
| **Layer 2** | -0.05 [-0.05, 0.16] | -1.45 | | 0.554 | 0.009 | $8 \times 10^{-5}$ |
| **Layer 3** | 0.39 [0.06, 0.61] | -0.77 | 1.33 | | 0.082 | $2 \times 10^{-4}$ |
| **Layer 4** | 0.62 [0.34, 0.79] | 0.26 | 3.26 | 2.51 | | 0.383 |
| **Layer 5** | 0.71 [0.41, 0.86] | 0.82 | 4.47 | 4.26 | 1.77 | |
| **Tuned: major extent** | | | | | | |
| **Layer 1** | 4.40 [0.09, 9.62] | | 0.770 | 0.797 | 0.877 | 0.797 |
| **Layer 2** | 1.10 [0.56, 1.37] | 1.14 | | 0.816 | 0.324 | $3 \times 10^{-4}$ |
| **Layer 3** | 0.76 [0.51, 1.01] | 0.88 | -0.57 | | 0.363 | $6 \times 10^{-6}$ |
| **Layer 4** | 0.57 [0.36, 0.91] | 0.15 | -1.92 | -1.80 | | 0.003 |
| **Layer 5** | 0.40 [0.27, 0.60] | -0.98 | -4.11 | -4.98 | -3.60 | |
| **Tuned: minor extent** | | | | | | |
| **Layer 1** | 0.10 [0.01, 1.15] | Kruskal-Wallis not significant | | | | |
| **Layer 2** | 0.11 [0.05, 0.26] | | | | | |
| **Layer 3** | 0.14 [0.09, 0.21] | | | | | |
| **Layer 4** | 0.15 [0.07, 0.33] | | | | | |
| **Layer 5** | 0.16 [0.07, 0.24] | | | | | |
| **Tuned: ratio between extents** | | | | | | |
| **Layer 1** | 0.38 [0.15, 0.62] | | 0.589 | 0.589 | 0.928 | 0.928 |
| **Layer 2** | 0.19 [0.10, 0.33] | -1.33 | | 0.964 | 0.499 | 0.006 |
| **Layer 3** | 0.20 [0.12, 0.34] | -1.40 | 0.05 | | 0.217 | $1 \times 10^{-5}$ |
| **Layer 4** | 0.28 [0.18, 0.47] | -0.55 | 1.60 | 2.12 | | 0.024 |
| **Layer 5** | 0.38 [0.24, 0.52] | 0.38 | 3.41 | 4.81 | 2.96 | |
| **Tuned: angulation towards x-axis** | | | | | | |
| **Layer 1** | 65.54 [37.42, 82.68] | | 1.000 | 1.000 | 1.000 | 1.000 |
| **Layer 2** | 63.11 [54.80, 64.85] | 0.00 | | 0.257 | 0.178 | 0.195 |
| **Layer 3** | 56.39 [50.68, 63.53] | 0.00 | 4.14 | | 0.257 | 0.199 |
| **Layer 4** | 49.25 [45.59, 52.06] | 0.00 | 5.47 | 4.15 | | 0.980 |
| **Layer 5** | 47.86 [45.43, 51.69] | 0.00 | 5.11 | 4.87 | 0.37 | |
| **Tuned: exponent on frequency** | | | | | | |
| **Layer 1** | 1.00 [1.00, 1.00] | Kruskal-Wallis not significant | | | | |
| **Layer 2** | 0.46 [0.09, 0.74] | | | | | |
| **Layer 3** | 0.20 [0.00, 0.77] | | | | | |
| **Layer 4** | 0.75 [0.00, 1.00] | | | | | |
| **Layer 5** | 0.34 [0.00, 1.00] | | | | | |

**Supplementary Table 8A: Some parameters of the monotonic and tuned response functions change over network layers of the layer-size-matched five-layer indRNN trained on repetitive events.** For all parameters, except the angulation towards the x-axis, given are the median [IQR] parameter values of the classified nodes for the monotonic and tuned response functions. Note that the tuned response function preferences and extents are expressed in seconds, where each time step represented 50 ms. The test statistics comparing each pair of network layers (post-hoc Dunn's test) are given in black cells, with corresponding corrected *p*-values (Holm-Šidák) of this comparison in the gray cells. For the angulation towards the x-axis, given are the circular median [95% ci of the circular median] for classified nodes classified as tuned. Here, the test statistics are pairwise common median tests, with corresponding corrected *p*-values (Holm-Šidák).Significant comparisons underlined.

| | Median [IQR] | Layer 1 | Layer 2 | Layer 3 | Layer 4 | Layer 5 |
|---|---|---|---|---|---|---|
| **Monotonic: Exponent on duration** | | | | | | |
| **Layer 1** | 1.00 [1.00, 1.00] | | $2 \times 10^{-23}$ | $1 \times 10^{-29}$ | $2 \times 10^{-33}$ | $2 \times 10^{-30}$ |
| **Layer 2** | 1.00 [0.95, 1.00] | -10.17 | | 0.356 | 0.069 | 0.214 |
| **Layer 3** | 1.00 [0.92, 1.00] | -11.50 | -1.62 | | 0.755 | 0.866 |
| **Layer 4** | 1.00 [0.91, 1.00] | -12.26 | -2.52 | -0.89 | | 0.866 |
| **Layer 5** | 1.00 [0.86, 1.00] | -11.64 | -1.99 | -0.39 | 0.48 | |
| **Monotonic: Exponent on period** | | | | | | |
| **Layer 1** | 1.00 [1.00, 1.00] | | $4 \times 10^{-63}$ | $1 \times 10^{-77}$ | $4 \times 10^{-75}$ | $2 \times 10^{-85}$ |
| **Layer 2** | 0.02 [0.01, 0.05] | -16.89 | | 0.093 | 0.143 | 0.001 |
| **Layer 3** | 0.02 [0.01, 0.03] | -18.77 | -2.34 | | 0.809 | 0.285 |
| **Layer 4** | 0.02 [0.00, 0.04] | -18.45 | -2.08 | 0.24 | | 0.267 |
| **Layer 5** | 0.02 [0.00, 0.03] | -19.72 | -3.73 | -1.42 | -1.65 | |
| **Monotonic: ratio between duration and period component** | | | | | | |
| **Layer 1** | Inf [135.42, Inf] | | $2 \times 10^{-59}$ | $4 \times 10^{-88}$ | $9 \times 10^{-66}$ | $3 \times 10^{-90}$ |
| **Layer 2** | 0.60 [0.17, 2.17] | -16.37 | | $9 \times 10^{-5}$ | 0.341 | $6 \times 10^{-6}$ |
| **Layer 3** | 0.34 [0.13, 0.99] | -20.01 | -4.28 | | 0.011 | 0.489 |
| **Layer 4** | 0.57 [0.17, 1.58] | -17.25 | -1.32 | 2.89 | | 0.002 |
| **Layer 5** | 0.31 [0.11, 1.02] | -20.26 | -4.88 | -0.69 | -3.51 | |
| **Tuned: preferred duration** | | | | | | |
| **Layer 1** | 0.06 [-0.05, 0.08] | | 0.509 | 0.001 | $1 \times 10^{-4}$ | $8 \times 10^{-5}$ |
| **Layer 2** | 0.07 [0.07, 0.14] | 1.39 | | 0.480 | 0.242 | 0.347 |
| **Layer 3** | 0.34 [0.07, 0.76] | 3.75 | 1.54 | | 0.882 | 0.882 |
| **Layer 4** | 0.18 [0.17, 0.76] | 4.37 | 2.07 | 0.66 | | 0.882 |
| **Layer 5** | 0.26 [0.16, 0.45] | 4.47 | 1.82 | 0.21 | -0.53 | |
| **Tuned: preferred period** | | | | | | |
| **Layer 1** | 0.02 [-0.04, 0.08] | | 0.480 | 0.002 | 0.002 | 0.021 |
| **Layer 2** | 0.41 [0.33, 0.67] | 1.70 | | 0.728 | 0.728 | 0.887 |
| **Layer 3** | 0.70 [0.33, 0.84] | 3.79 | 1.28 | | 0.971 | 0.728 |
| **Layer 4** | 0.71 [0.57, 0.83] | 3.75 | 1.30 | 0.04 | | 0.728 |
| **Layer 5** | 0.48 [0.25, 0.76] | 3.00 | 0.43 | -1.20 | -1.21 | |
| **Tuned: major extent** | | | | | | |
| **Layer 1** | 0.85 [0.64, 0.88] | | 0.044 | $6 \times 10^{-5}$ | 0.001 | 0.005 |
| **Layer 2** | 0.34 [0.32, 1.25] | -2.72 | | 0.859 | 0.938 | 0.938 |
| **Layer 3** | 0.43 [0.26, 0.45] | -4.53 | -0.86 | | 0.938 | 0.515 |
| **Layer 4** | 0.42 [0.33, 0.47] | -4.00 | -0.52 | 0.41 | | 0.799 |
| **Layer 5** | 0.50 [0.29, 0.60] | -3.42 | 0.32 | 1.58 | 1.09 | |
| **Tuned: minor extent** | | | | | | |
| **Layer 1** | 0.01 [0.01, 0.01] | | 0.044 | 0.005 | 0.098 | 0.044 |
| **Layer 2** | 0.03 [0.01, 0.09] | 2.81 | | 1.000 | 0.893 | 0.893 |
| **Layer 3** | 0.02 [0.02, 0.03] | 3.48 | -0.02 | | 0.893 | 0.869 |
| **Layer 4** | 0.02 [0.01, 0.03] | 2.44 | -0.77 | -0.91 | | 1.000 |
| **Layer 5** | 0.02 [0.01, 0.04] | 2.80 | -0.86 | -1.06 | -0.03 | |
| **Tuned: ratio between extents** | | | | | | |
| **Layer 1** | 0.01 [0.01, 0.02] | | 0.052 | $2 \times 10^{-4}$ | 0.025 | 0.002 |
| **Layer 2** | 0.10 [0.02, 0.27] | 2.67 | | 0.910 | 0.985 | 0.987 |
| **Layer 3** | 0.06 [0.05, 0.28] | 4.31 | 0.75 | | 0.791 | 0.827 |
| **Layer 4** | 0.05 [0.03, 0.10] | 2.95 | -0.25 | -1.20 | | 0.985 |
| **Layer 5** | 0.05 [0.03, 0.10] | 3.75 | -0.02 | -1.05 | 0.31 | |
| **Tuned: angulation towards x-axis** | | | | | | |
| **Layer 1** | 67.19 [44.92, 88.82] | Common median test not significant | | | | |
| **Layer 2** | 88.74 [43.37, 88.85] | | | | | |
| **Layer 3** | 45.54 [30.57, 60.82] | | | | | |
| **Layer 4** | 85.90 [45.65, 89.38] | | | | | |
| **Layer 5** | 49.36 [45.48, 51.95] | | | | | |
| **Tuned: exponent on frequency** | | | | | | |
| **Layer 1** | 0.08 [0.00, 0.76] | Kruskal-Wallis not significant | | | | |
| **Layer 2** | 0.99 [0.00, 1.00] | | | | | |
| **Layer 3** | 0.00 [0.00, 0.00] | | | | | |
| **Layer 4** | 0.00 [0.00, 0.00] | | | | | |
| **Layer 5** | 0.00 [0.00, 0.34] | | | | | |

**Supplementary Table 8B: Some parameters of the monotonic and tuned response functions change over network layers of the layer-size-matched five-layer indRNN before training.** Like Supplementary Table 8A, but for the layer-size-matched five-layer indRNN trained on repetitive events

| | Median [IQR] | Layer 1 | Layer 2 | Layer 3 | Layer 4 | Layer 5 |
|---|---|---|---|---|---|---|
| **Monotonic: exponent on duration** | | | | | | |
| **Layer 1** | 1.00 [1.00, 1.00] | | $8 \times 10^{-17}$ | $7 \times 10^{-11}$ | $6 \times 10^{-5}$ | $6 \times 10^{-7}$ |
| **Layer 2** | 0.94 [0.62, 1.00] | -8.60 | | 0.845 | 0.316 | 0.845 |
| **Layer 3** | 0.93 [0.52, 1.00] | -6.84 | 0.73 | | 0.755 | 0.976 |
| **Layer 4** | 0.95 [0.54, 1.00] | -4.46 | 1.87 | 1.16 | | 0.795 |
| **Layer 5** | 0.94 [0.37, 1.00] | -5.38 | 0.63 | 0.03 | -0.98 | |
| **Monotonic: exponent on period** | | | | | | |
| **Layer 1** | 1.00 [0.02, 1.00] | | $1 \times 10^{-9}$ | $2 \times 10^{-8}$ | $4 \times 10^{-6}$ | $2 \times 10^{-5}$ |
| **Layer 2** | 0.01 [0.00, 1.00] | -6.42 | | 1.000 | 1.000 | 1.000 |
| **Layer 3** | 0.02 [0.00, 0.65] | -6.01 | -0.29 | | 1.000 | 1.000 |
| **Layer 4** | 0.01 [0.00, 1.00] | -5.01 | -0.18 | 0.06 | | 1.000 |
| **Layer 5** | 0.01 [0.00, 1.00] | -4.69 | -0.17 | 0.06 | 0.01 | |
| **Monotonic: ratio between duration and period component** | | | | | | |
| **Layer 1** | 69.55 [1.64, Inf] | | 0.002 | $5 \times 10^{-6}$ | 0.009 | 0.001 |
| **Layer 2** | 5.67 [1.33, 263.75] | -3.65 | | 0.467 | 0.874 | 0.713 |
| **Layer 3** | 4.17 [0.48, 31.88] | -5.03 | -1.65 | | 0.842 | 0.911 |
| **Layer 4** | 3.24 [1.06, 73.42] | -3.23 | -0.46 | 0.90 | | 0.874 |
| **Layer 5** | 4.47 [0.73, 29.11] | -3.86 | -1.22 | 0.11 | -0.68 | |
| **Tuned: preferred duration** | | | | | | |
| **Layer 1** | 0.23 [0.07, 0.39] | Kruskal-Wallis not significant | | | | |
| **Layer 2** | 0.14 [0.07, 0.33] | | | | | |
| **Layer 3** | 0.19 [0.07, 0.47] | | | | | |
| **Layer 4** | 0.15 [0.09, 0.43] | | | | | |
| **Layer 5** | 0.17 [0.09, 0.51] | | | | | |
| **Tuned: preferred period** | | | | | | |
| **Layer 1** | 0.7 [0.13, 0.81] | Kruskal-Wallis not significant | | | | |
| **Layer 2** | 0.51 [0.20, 0.88] | | | | | |
| **Layer 3** | 0.64 [0.21, 0.83] | | | | | |
| **Layer 4** | 0.61 [0.21, 0.86] | | | | | |
| **Layer 5** | 0.69 [0.27, 0.89] | | | | | |
| **Tuned: major extent** | | | | | | |
| **Layer 1** | 0.45 [0.17, 0.66] | . | 0.999 | 0.999 | 0.999 | 0.253 |
| **Layer 2** | 0.29 [0.22, 0.55] | -0.01 | . | 0.999 | 0.999 | 0.253 |
| **Layer 3** | 0.41 [0.21, 0.57] | 0.29 | 0.32 | . | 0.977 | 0.008 |
| **Layer 4** | 0.31 [0.18, 0.60] | -0.21 | -0.19 | -0.73 | . | 0.033 |
| **Layer 5** | 0.20 [0.13, 0.35] | -2.07 | -2.10 | -3.34 | -2.90 | . |
| **Tuned: minor extent** | | | | | | |
| **Layer 1** | 0.02 [0.01, 0.10] | Kruskal-Wallis not significant | | | | |
| **Layer 2** | 0.03 [0.01, 0.10] | | | | | |
| **Layer 3** | 0.03 [0.01, 0.04] | | | | | |
| **Layer 4** | 0.03 [0.01, 0.06] | | | | | |
| **Layer 5** | 0.02 [0.01, 0.04] | | | | | |
| **Tuned: ratio between extents** | | | | | | |
| **Layer 1** | 0.12 [0.07, 0.27] | Kruskal-Wallis not significant | | | | |
| **Layer 2** | 0.15 [0.09, 0.23] | | | | | |
| **Layer 3** | 0.10 [0.03, 0.14] | | | | | |
| **Layer 4** | 0.10 [0.03, 0.23] | | | | | |
| **Layer 5** | 0.13 [0.06, 0.23] | | | | | |
| **Tuned: exponent on frequency** | | | | | | |
| **Layer 1** | 0.69 [0.00, 1.00] | Kruskal-Wallis not significant | | | | |
| **Layer 2** | 1.00 [0.21, 1.00] | | | | | |
| **Layer 3** | 0.00 [0.00, 1.00] | | | | | |
| **Layer 4** | 0.00 [0.00, 1.00] | | | | | |
| **Layer 5** | 0.42 [0.00, 1.00] | | | | | |
| **Tuned: angulation towards x-axis** | | | | | | |
| **Layer 1** | 58.44 [45.90, 67.87] | Common median test not significant | | | | |
| **Layer 2** | 63.62 [49.91, 82.43] | | | | | |
| **Layer 3** | 58.52 [50.11, 66.86] | | | | | |
| **Layer 4** | 58.06 [48.92, 71.26] | | | | | |
| **Layer 5** | 58.10 [49.93, 70.47] | | | | | |
| **Tuned: exponent on frequency** | | | | | | |
| **Layer 1** | 0.69 [0.00, 1.00] | Kruskal-Wallis not significant | | | | |
| **Layer 2** | 1.00 [0.21, 1.00] | | | | | |
| **Layer 3** | 0.00 [0.00, 1.00] | | | | | |
| **Layer 4** | 0.00 [0.00, 1.00] | | | | | |
| **Layer 5** | 0.42 [0.00, 1.00] | | | | | |

**Supplementary Table 8C: Some parameters of the monotonic and tuned response functions change over network layers of the layer-size-matched five-layer indRNN trained on shuffled data.** Like Supplementary Table 8A, but for the layer-size-matched five-layer indRNN trained on shuffled data

| | Median [IQR] | Layer 1 | Layer 2 | Layer 3 | Layer 4 | Layer 5 |
|---|---|---|---|---|---|---|
| **Monotonic: Exponent on duration** | | | | | | |
| **Layer 1** | 0.00 [0.0, 1.00] | | 0.362 | 0.163 | 0.005 | $2 \times 10^{-4}$ |
| **Layer 2** | 1.00 [0.0, 1.00] | 1.61 | | 0.553 | 0.325 | 0.045 |
| **Layer 3** | 0.56 [0.0, 1.00] | 2.24 | 0.69 | | 0.553 | 0.226 |
| **Layer 4** | 1.00 [0.0, 1.00] | 3.47 | 1.78 | 1.00 | | 0.553 |
| **Layer 5** | 1.00 [0.0, 1.00] | 4.27 | 2.76 | 2.04 | 1.19 | |
| **Monotonic: Exponent on period** | | | | | | |
| **Layer 1** | 0.02 [0.02, 0.05] | | 0.917 | 0.667 | 0.667 | 0.042 |
| **Layer 2** | 0.02 [0.02, 0.49] | 0.37 | | 0.744 | 0.744 | 0.123 |
| **Layer 3** | 0.02 [0.02, 1.00] | 1.46 | 1.06 | | 0.930 | 0.667 |
| **Layer 4** | 0.02 [0.02, 1.00] | 1.44 | 1.02 | -0.09 | | 0.654 |
| **Layer 5** | 1.00s [0.02, 1.00] | 2.86 | 2.45 | 1.40 | 1.54 | |
| **Monotonic: ratio between duration and period component** | | | | | | |
| **Layer 1** | $2 \times 10^{-15}$ [0, 2.08] | | 0.348 | 0.168 | 0.005 | $1 \times 10^{-4}$ |
| **Layer 2** | $5 \times 10^{-9}$ [0, 28.60] | 1.66 | | 0.524 | 0.348 | 0.039 |
| **Layer 3** | $4 \times 10^{-9}$ [0, Inf] | 2.23 | 0.64 | | 0.524 | 0.186 |
| **Layer 4** | $1 \times 10^{-8}$ [0, Inf] | 3.47 | 1.74 | 1.02 | | 0.500 |
| **Layer 5** | $9 \times 10^{9}$ [$3 \times 10^{-18}$, Inf] | 4.35 | 2.81 | 2.12 | 1.26 | |

**Supplementary Table 8D. Parameters of the monotonic response function change over network layers of the layer-size-matched five-layer neural network without recurrency.** Given are the median [IQR] parameter values of the classified nodes for the monotonic response function. The test statistics comparing each pair of network layers (post-hoc Dunn's test) are given in black cells, with corresponding corrected *p*-values (Holm-Šidák) of this comparison in the gray cells. Significant comparisons underlined

| | Median [IQR] | V1 | V2 | V3 | hV4 | LO1 | LO2 | TO1 | TO2 | V3AB | IPS0 | IPS1 | IPS2 | IPS3 | IPS4 | IPS5 | sPCS1 | sPCS2 | iPCS |
|---|---|---|---|---|---|---|---|---|---|---|---|---|---|---|---|---|---|---|---|
| **Monotonic: exponent on duration** | | | | | | | | | | | | | | | | | | | |
| **V1** | 0.81 [0.6, 0.97] | | 1.000 | 1.000 | 1.000 | 1.000 | 1.000 | 1.000 | 1.000 | 1.000 | 1.000 | 1.000 | 1.000 | 1.000 | 1.000 | 1.000 | 1.000 | 1.000 | 0.462 |
| **V2** | 0.88 [0.72, 1.00] | 1.04 | | 1.000 | 1.000 | 0.490 | 1.000 | 1.000 | 0.998 | 0.287 | 1.000 | 0.594 | 0.610 | 0.980 | 0.958 | 1.000 | 1.000 | 0.586 | 0.002 |
| **V3** | 0.75 [0.61, 0.92] | -0.50 | -1.76 | | 1.000 | 1.000 | 0.879 | 1.000 | 1.000 | 1.000 | 1.000 | 1.000 | 1.000 | 1.000 | 1.000 | 1.000 | 1.000 | 1.000 | 0.583 |
| **hV4** | 0.94 [0.72, 1.00] | 1.03 | 0.05 | 1.69 | | 0.622 | 1.000 | 1.000 | 1.000 | 0.410 | 1.000 | 0.713 | 0.720 | 0.991 | 0.980 | 1.000 | 1.000 | 0.685 | 0.006 |
| **LO1** | 0.70 [0.50, 0.82] | -1.46 | -2.82 | -1.12 | -2.68 | | 0.072 | 0.994 | 1.000 | 1.000 | 1.000 | 1.000 | 1.000 | 1.000 | 1.000 | 1.000 | 1.000 | 1.000 | 1.000 |
| **LO2** | 0.91 [0.75, 1.00] | 1.55 | 0.53 | 2.39 | 0.44 | 3.48 | | 1.000 | 0.745 | 0.033 | 0.990 | 0.107 | 0.122 | 0.553 | 0.436 | 0.980 | 1.000 | 0.129 | $8\times10^{-5}$ |
| **TO1** | 0.81 [0.67, 0.89] | 0.28 | -0.90 | 0.91 | -0.89 | 2.02 | -1.49 | | 1.000 | 0.941 | 1.000 | 0.998 | 0.998 | 1.000 | 1.000 | 1.000 | 1.000 | 0.995 | 0.044 |
| **TO2** | 0.78 [0.70, 0.83] | -0.62 | -1.91 | -0.14 | -1.83 | 0.99 | -2.55 | -1.06 | | 1.000 | 1.000 | 1.000 | 1.000 | 1.000 | 1.000 | 1.000 | 1.000 | 1.000 | 0.686 |
| **V3AB** | 0.69 [0.49, 0.84] | -1.69 | -3.04 | -1.40 | -2.90 | -0.31 | -3.69 | -2.28 | -1.28 | | 1.000 | 1.000 | 1.000 | 1.000 | 1.000 | 1.000 | 0.999 | 1.000 | 1.000 |
| **IPS0** | 0.78 [0.71, 0.89] | -0.22 | -1.45 | 0.32 | -1.41 | 1.43 | -2.07 | -0.59 | 0.46 | 1.70 | | 1.000 | 1.000 | 1.000 | 1.000 | 1.000 | 1.000 | 1.000 | 0.295 |
| **IPS1** | 0.71 [0.50, 0.86] | -1.37 | -2.71 | -1.03 | -2.59 | 0.08 | -3.36 | -1.92 | -0.90 | 0.38 | -1.34 | | 1.000 | 1.000 | 1.000 | 1.000 | 1.000 | 1.000 | 1.000 |
| **IPS2** | 0.79 [0.00, 0.84] | -1.39 | -2.69 | -1.06 | -2.58 | 0.02 | -3.32 | -1.92 | -0.93 | 0.32 | -1.36 | -0.06 | | 1.000 | 1.000 | 1.000 | 1.000 | 1.000 | 1.000 |
| **IPS3** | 0.79 [0.60, 0.84] | -0.89 | -2.14 | -0.47 | -2.06 | 0.61 | -2.76 | -1.34 | -0.34 | 0.89 | -0.77 | 0.52 | 0.57 | | 1.000 | 1.000 | 1.000 | 1.000 | 0.976 |
| **IPS4** | 0.76 [0.66, 0.85] | -0.94 | -2.23 | -0.52 | -2.14 | 0.59 | -2.87 | -1.42 | -0.39 | 0.88 | -0.83 | 0.50 | 0.55 | -0.03 | | 1.000 | 1.000 | 1.000 | 0.971 |
| **IPS5** | 0.79 [0.71, 0.85] | -0.39 | -1.57 | 0.09 | -1.52 | 1.14 | -2.14 | -0.76 | 0.22 | 1.40 | -0.21 | 1.05 | 1.08 | 0.53 | 0.57 | | 1.000 | 1.000 | 0.657 |
| **sPCS1** | 0.84 [0.80, 0.87] | 0.34 | -0.54 | 0.81 | -0.56 | 1.64 | -0.97 | 0.13 | 0.92 | 1.86 | 0.57 | 1.57 | 1.59 | 1.15 | 1.19 | 0.71 | | 1.000 | 0.442 |
| **sPCS2** | 0.76 [0.31, 0.84] | -1.50 | -2.72 | -1.18 | -2.62 | -0.17 | -3.30 | -1.99 | -1.07 | 0.11 | -1.46 | -0.24 | -0.18 | -0.72 | -0.71 | -1.20 | -1.68 | | 1.000 |
| **iPCS** | 0.54 [0.00, 0.76] | -2.84 | -4.33 | -2.73 | -4.10 | -1.61 | -5.03 | -3.62 | -2.61 | -1.27 | -3.03 | -1.68 | -1.57 | -2.16 | -2.19 | -2.65 | -2.87 | -1.30 | |
| **Monotonic: exponent on period** | | | | | | | | | | | | | | | | | | | |
| **V1** | 1.00 [0.96, 1.00] | | 1.000 | 1.000 | 1.000 | 1.000 | 1.000 | 1.000 | 0.829 | 0.363 | 0.901 | 0.608 | 0.008 | 0.008 | 0.041 | 0.001 | 0.008 | 0.002 | 0.466 |
| **V2** | 1.00 [0.47, 1.00] | -0.74 | | 1.000 | 1.000 | 1.000 | 1.000 | 1.000 | 0.995 | 0.800 | 0.998 | 0.947 | 0.033 | 0.031 | 0.160 | 0.002 | 0.032 | 0.007 | 0.892 |
| **V3** | 1.00 [0.51, 1.00] | -0.70 | 0.07 | | 1.000 | 1.000 | 1.000 | 1.000 | 0.985 | 0.689 | 0.995 | 0.901 | 0.017 | 0.016 | 0.090 | 0.001 | 0.020 | 0.003 | 0.795 |
| **hV4** | 1.00 [0.66, 1.00] | 0.09 | 0.88 | 0.84 | | 1.000 | 1.000 | 1.000 | 0.656 | 0.204 | 0.764 | 0.394 | 0.003 | 0.003 | 0.015 | $1\times10^{-4}$ | 0.004 | $5\times10^{-4}$ | 0.282 |
| **LO1** | 1.00 [0.63, 1.00] | -0.07 | 0.77 | 0.72 | -0.18 | | 1.000 | 1.000 | 0.617 | 0.165 | 0.742 | 0.350 | 0.001 | 0.001 | 0.008 | $4\times10^{-5}$ | 0.003 | $2\times10^{-4}$ | 0.239 |
| **LO2** | 1.00 [0.87, 1.00] | 0.43 | 1.35 | 1.33 | 0.35 | 0.58 | | 0.995 | 0.127 | 0.019 | 0.197 | 0.051 | $5\times10^{-5}$ | $4\times10^{-5}$ | $4\times10^{-4}$ | $1\times10^{-6}$ | $3\times10^{-4}$ | $8\times10^{-6}$ | 0.031 |
| **TO1** | 0.70 [0.51, 1.00] | -1.03 | -0.30 | -0.38 | -1.19 | -1.10 | -1.73 | | 1.000 | 0.916 | 1.000 | 0.988 | 0.055 | 0.052 | 0.254 | 0.004 | 0.052 | 0.011 | 0.959 |
| **TO2** | 0.49 [0.43, 0.75] | -2.31 | -1.74 | -1.87 | -2.53 | -2.57 | -3.23 | -1.50 | | 1.000 | 1.000 | 1.000 | 0.943 | 0.943 | 0.999 | 0.393 | 0.773 | 0.587 | 1.000 |
| **V3AB** | 0.44 [0.37, 1.00] | -2.84 | -2.35 | -2.49 | -3.07 | -3.14 | -3.78 | -2.14 | -0.72 | | 1.000 | 1.000 | 1.000 | 1.000 | 1.000 | 0.960 | 0.993 | 0.987 | 1.000 |
| **IPS0** | 0.51 [0.41, 1.00] | -2.19 | -1.61 | -1.74 | -2.40 | -2.43 | -3.08 | -1.37 | 0.12 | 0.83 | | 1.000 | 0.905 | 0.905 | 0.998 | 0.319 | 0.708 | 0.495 | 1.000 |
| **IPS1** | 0.46 [0.34, 1.00] | -2.58 | -2.05 | -2.19 | -2.80 | -2.86 | -3.50 | -1.83 | -0.38 | 0.34 | -0.50 | | 0.998 | 0.998 | 1.000 | 0.790 | 0.946 | 0.901 | 1.000 |
| **IPS2** | 0.34 [0.31, 0.80] | -3.99 | -3.63 | -3.81 | -4.27 | -4.44 | -5.10 | -3.48 | -2.07 | -1.29 | -2.17 | -1.65 | | 1.000 | 1.000 | 1.000 | 1.000 | 1.000 | 1.000 |
| **IPS3** | 0.35 [0.31, 0.98] | -4.00 | -3.64 | -3.83 | -4.28 | -4.46 | -5.13 | -3.49 | -2.07 | -1.29 | -2.17 | -1.65 | 0.02 | | 1.000 | 1.000 | 1.000 | 1.000 | 1.000 |
| **IPS4** | 0.40 [0.34, 0.49] | -3.57 | -3.16 | -3.34 | -3.84 | -3.99 | -4.67 | -2.99 | -1.52 | -0.74 | -1.63 | -1.10 | 0.59 | 0.58 | | 1.000 | 1.000 | 1.000 | 1.000 |
| **IPS5** | 0.34 [0.32, 0.42] | -4.61 | -4.32 | -4.52 | -4.91 | -5.13 | -5.80 | -4.19 | -2.80 | -2.00 | -2.90 | -2.37 | -0.71 | -0.74 | -1.33 | | 1.000 | 1.000 | 0.922 |
| **sPCS1** | 0.31 [0.29, 0.50] | -3.99 | -3.64 | -3.76 | -4.20 | -4.27 | -4.77 | -3.50 | -2.39 | -1.77 | -2.47 | -2.06 | -0.73 | -0.75 | -1.22 | -0.15 | | 1.000 | 0.985 |
| **sPCS2** | 0.36 [0.29, 0.40] | -4.38 | -4.05 | -4.23 | -4.64 | -4.82 | -5.44 | -3.92 | -2.60 | -1.85 | -2.69 | -2.19 | -0.63 | -0.65 | -1.21 | 0.05 | 0.18 | | 0.966 |
| **iPCS** | 0.48 [0.34, 1.00] | -2.72 | -2.22 | -2.36 | -2.95 | -3.01 | -3.65 | -2.01 | -0.59 | 0.13 | -0.70 | -0.21 | 1.42 | 1.41 | 0.87 | 2.13 | 1.87 | 1.97 | |
| **Monotonic: ratio between duration and period components** | | | | | | | | | | | | | | | | | | | |
| **V1** | Inf [Inf, Inf] | | 1.000 | 0.996 | 1.000 | 1.000 | 1.000 | 0.996 | 0.059 | 0.054 | 0.292 | 0.561 | 0.009 | 0.021 | 0.025 | 0.015 | 0.046 | 0.001 | 0.001 |
| **V2** | Inf [2.64, Inf] | -0.91 | | 1.000 | 1.000 | 1.000 | 1.000 | 1.000 | 0.347 | 0.318 | 0.873 | 0.984 | 0.065 | 0.153 | 0.170 | 0.107 | 0.237 | 0.010 | 0.009 |
| **V3** | Inf [1.02, Inf] | -1.81 | -0.99 | | 1.000 | 1.000 | 0.995 | 1.000 | 0.983 | 0.969 | 1.000 | 1.000 | 0.598 | 0.845 | 0.880 | 0.736 | 0.845 | 0.162 | 0.174 |
| **hV4** | Inf [4.18, Inf] | -0.69 | 0.19 | 1.11 | | 1.000 | 1.000 | 1.000 | 0.350 | 0.318 | 0.853 | 0.976 | 0.072 | 0.160 | 0.179 | 0.114 | 0.222 | 0.012 | 0.012 |
| **LO1** | Inf [1.51, Inf] | -1.55 | -0.70 | 0.28 | -0.85 | | 1.000 | 1.000 | 0.911 | 0.873 | 1.000 | 1.000 | 0.385 | 0.654 | 0.713 | 0.529 | 0.693 | 0.088 | 0.092 |
| **LO2** | Inf [6.89, Inf] | -0.25 | 0.77 | 1.83 | 0.53 | 1.52 | | 0.995 | 0.018 | 0.019 | 0.160 | 0.404 | 0.002 | 0.006 | 0.006 | 0.004 | 0.027 | $2\times10^{-4}$ | $1\times10^{-4}$ |
| **TO1** | 2.65 [1.26, Inf] | -1.81 | -0.98 | 0.01 | -1.11 | -0.26 | -1.83 | | 0.979 | 0.962 | 1.000 | 1.000 | 0.567 | 0.828 | 0.867 | 0.713 | 0.831 | 0.151 | 0.160 |
| **TO2** | 1.00 [0.80, 9.32] | -3.50 | -2.89 | -1.97 | -2.89 | -2.21 | -3.83 | -2.00 | | 1.000 | 1.000 | 1.000 | 1.000 | 1.000 | 1.000 | 1.000 | 1.000 | 1.000 | 1.000 |
| **V3AB** | 1.12 [0.39, Inf] | -3.53 | -2.93 | -2.05 | -2.93 | -2.29 | -3.82 | -2.08 | -0.19 | | 1.000 | 1.000 | 1.000 | 1.000 | 1.000 | 1.000 | 1.000 | 1.000 | 1.000 |
| **IPS0** | 1.54 [0.74, Inf] | -2.96 | -2.29 | -1.35 | -2.32 | -1.60 | -3.19 | -1.37 | 0.61 | 0.76 | | 1.000 | 1.000 | 1.000 | 1.000 | 1.000 | 1.000 | 0.981 | 0.994 |
| **IPS1** | 1.25 [0.72, Inf] | -2.66 | -1.96 | -1.03 | -2.02 | -1.28 | -2.82 | -1.05 | 0.88 | 1.02 | 0.28 | | 1.000 | 1.000 | 1.000 | 1.000 | 1.000 | 0.917 | 0.960 |
| **IPS2** | 0.89 [0.36, 2.97] | -4.01 | -3.47 | -2.62 | -3.44 | -2.84 | -4.37 | -2.65 | -0.78 | -0.57 | -1.35 | -1.59 | | 1.000 | 1.000 | 1.000 | 1.000 | 1.000 | 1.000 |
| **IPS3** | 0.92 [0.53, 9.18] | -3.78 | -3.21 | -2.34 | -3.19 | -2.57 | -4.11 | -2.37 | -0.48 | -0.27 | -1.05 | -1.30 | 0.30 | | 1.000 | 1.000 | 1.000 | 1.000 | 1.000 |
| **IPS4** | 1.25 [0.55, 2.69] | -3.74 | -3.16 | -2.27 | -3.14 | -2.50 | -4.09 | -2.30 | -0.35 | -0.15 | -0.94 | -1.20 | 0.44 | 0.13 | | 1.000 | 1.000 | 1.000 | 1.000 |
| **IPS5** | 1.03 [0.62, 1.62] | -3.88 | -3.32 | -2.47 | -3.30 | -2.70 | -4.20 | -2.50 | -0.65 | -0.45 | -1.21 | -1.45 | 0.11 | -0.18 | -0.31 | | 1.000 | 1.000 | 1.000 |
| **sPCS1** | 0.68 [0.42, 5.36] | -3.57 | -3.04 | -2.34 | -3.07 | -2.53 | -3.72 | -2.36 | -0.89 | -0.72 | -1.33 | -1.53 | -0.26 | -0.50 | -0.61 | -0.35 | | 1.000 | 1.000 |
| **sPCS2** | 0.63 [0.32, 1.37] | -4.46 | -3.97 | -3.18 | -3.93 | -3.38 | -4.82 | -3.21 | -1.46 | -1.23 | -1.99 | -2.20 | -0.69 | -0.98 | -1.12 | -0.79 | -0.32 | | 1.000 |
| **iPCS** | 0.36 [0.10, Inf] | -4.48 | -4.00 | -3.15 | -3.93 | -3.37 | -4.92 | -3.19 | -1.30 | -1.05 | -1.86 | -2.09 | -0.48 | -0.78 | -0.93 | -0.59 | -0.12 | 0.25 | |

**Supplementary Table 8E: Monotonic response function parameters change over visual field.** Like Supplementary Table 8A, but for the areas in the human brain. Therefore here, medians and IQRs are computed over included hemispheres. Data adapted from Hendrikx et al. (2022)

| | Median [IQR] | TLO | TTOP | TTOA | TPO | TLS | TPCI | TPCM | TPCS | TFI | TFS |
|---|---|---|---|---|---|---|---|---|---|---|---|
| **Tuned: preferred duration** | | | | | | | | | | | |
| **TLO** | 0.48 [0.40, 0.52] | | 1.000 | 0.979 | 1.000 | 0.443 | 1.000 | 1.000 | 0.944 | 0.459 | 1.000 |
| **TTOP** | 0.47 [0.41, 0.50] | -0.03 | | 0.980 | 1.000 | 0.459 | 1.000 | 1.000 | 0.940 | 0.471 | 1.000 |
| **TTOA** | 0.41 [0.37, 0.49] | -1.43 | -1.40 | | 0.851 | 0.999 | 0.998 | 0.954 | 0.094 | 0.999 | 0.997 |
| **TPO** | 0.45 [0.43, 0.54] | 0.43 | 0.46 | 1.85 | | 0.169 | 1.000 | 1.000 | 0.996 | 0.186 | 1.000 |
| **TLS** | 0.38 [0.33, 0.43] | -2.43 | -2.40 | -1.02 | -2.85 | | 0.717 | 0.336 | 0.003 | 1.000 | 0.667 |
| **TPCI** | 0.44 [0.38, 0.52] | -0.28 | -0.26 | 1.09 | -0.70 | 2.07 | | 1.000 | 0.851 | 0.728 | 1.000 |
| **TPCM** | 0.48 [0.40, 0.52] | 0.15 | 0.17 | 1.57 | -0.28 | 2.57 | 0.42 | | 0.973 | 0.359 | 1.000 |
| **TPCS** | 0.50 [0.47, 0.54] | 1.62 | 1.65 | 3.05 | 1.19 | 4.02 | 1.85 | 1.48 | | 0.003 | 0.826 |
| **TFI** | 0.39 [0.36, 0.44] | -2.39 | -2.36 | -1.01 | -2.81 | -0.01 | -2.04 | -2.53 | -3.96 | | 0.683 |
| **TFS** | 0.45 [0.39, 0.50] | -0.28 | -0.26 | 1.14 | -0.71 | 2.15 | 0.01 | -0.43 | -1.91 | 2.12 | |
| **Tuned: preferred period** | | | | | | | | | | | |
| **TLO** | 0.66 [0.56, 0.75] | | 0.331 | 0.007 | 0.990 | 0.001 | 0.869 | 0.623 | 1.000 | 0.987 | 0.834 |
| **TTOP** | 0.54 [0.50, 0.57] | -2.57 | | 0.995 | 0.995 | 0.878 | 1.000 | 1.000 | 0.862 | 0.997 | 1.000 |
| **TTOA** | 0.50 [0.42, 0.54] | -3.77 | -1.21 | | 0.432 | 1.000 | 0.869 | 0.957 | 0.086 | 0.568 | 0.863 |
| **TPO** | 0.56 [0.52, 0.66] | -1.34 | 1.23 | 2.44 | | 0.107 | 1.000 | 1.000 | 1.000 | 1.000 | 1.000 |
| **TLS** | 0.47 [0.42, 0.50] | -4.31 | -1.78 | -0.59 | -2.99 | | 0.472 | 0.654 | 0.013 | 0.184 | 0.443 |
| **TPCI** | 0.55 [0.46, 0.68] | -1.82 | 0.66 | 1.83 | -0.53 | 2.37 | | 1.000 | 0.996 | 1.000 | 1.000 |
| **TPCM** | 0.54 [0.48, 0.60] | -2.19 | 0.37 | 1.58 | -0.86 | 2.15 | -0.30 | | 0.974 | 1.000 | 1.000 |
| **TPCS** | 0.57 [0.55, 0.61] | -0.70 | 1.87 | 3.07 | 0.63 | 3.62 | 1.14 | 1.49 | | 1.000 | 0.995 |
| **TFI** | 0.58 [0.48, 0.65] | -1.38 | 1.10 | 2.26 | -0.09 | 2.80 | 0.42 | 0.74 | -0.71 | | 1.000 |
| **TFS** | 0.55 [0.50, 0.60] | -1.92 | 0.65 | 1.85 | -0.59 | 2.42 | -0.04 | 0.27 | -1.22 | -0.47 | |
| **Tuned: major extent** | | | | | | | | | | | |
| **TLO** | 0.96 [0.81, 1.09] | | 0.986 | 0.998 | 0.969 | 0.984 | 0.016 | 0.641 | 0.006 | 0.419 | 0.031 |
| **TTOP** | 1.04 [0.89, 1.10] | 1.02 | | 0.984 | 0.395 | 0.485 | $3 \times 10^{-4}$ | 0.067 | $6 \times 10^{-5}$ | 0.029 | 0.001 |
| **TTOA** | 0.90 [0.84, 1.03] | -0.22 | -1.25 | | 0.984 | 0.986 | 0.033 | 0.789 | 0.014 | 0.568 | 0.062 |
| **TPO** | 0.88 [0.79, 0.90] | -1.41 | -2.43 | -1.18 | | 0.998 | 0.567 | 0.993 | 0.416 | 0.986 | 0.724 |
| **TLS** | 0.87 [0.78, 0.94] | -1.29 | -2.29 | -1.06 | 0.10 | | 0.522 | 0.992 | 0.385 | 0.986 | 0.681 |
| **TPCI** | 0.70 [0.62, 0.82] | -3.54 | -4.53 | -3.32 | -2.18 | -2.24 | | 0.938 | 0.998 | 0.986 | 0.998 |
| **TPCM** | 0.83 [0.75, 0.90] | -2.07 | -3.09 | -1.84 | -0.66 | -0.75 | 1.54 | | 0.852 | 0.998 | 0.984 |
| **TPCS** | 0.69 [0.65, 0.79] | -3.80 | -4.82 | -3.58 | -2.39 | -2.45 | -0.14 | -1.73 | | 0.984 | 0.998 |
| **TFI** | 0.78 [0.72, 0.87] | -2.38 | -3.37 | -2.16 | -1.02 | -1.10 | 1.12 | -0.38 | 1.29 | | 0.989 |
| **TFS** | 0.72 [0.60, 0.85] | -3.35 | -4.37 | -3.12 | -1.94 | -2.01 | 0.30 | -1.28 | 0.45 | -0.85 | |
| **Tuned: minor extent** | | | | | | | | | | | |
| **TLO** | 0.25 [0.22, 0.28] | | 1.000 | 0.996 | 1.000 | 1.000 | 0.383 | 1.000 | 0.999 | 1.000 | 0.999 |
| **TTOP** | 0.25 [0.22, 0.27] | -0.04 | | 0.997 | 1.000 | 1.000 | 0.409 | 1.000 | 0.999 | 1.000 | 0.999 |
| **TTOA** | 0.22 [0.21, 0.25] | -1.38 | -1.34 | | 1.000 | 1.000 | 0.999 | 0.898 | 0.448 | 0.999 | 0.413 |
| **TPO** | 0.23 [0.21, 0.27] | -0.42 | -0.38 | 0.96 | | 1.000 | 0.717 | 1.000 | 0.991 | 1.000 | 0.988 |
| **TLS** | 0.24 [0.21, 0.26] | -0.65 | -0.62 | 0.70 | -0.24 | | 0.898 | 0.999 | 0.958 | 1.000 | 0.947 |
| **TPCI** | 0.20 [0.16, 0.23] | -2.53 | -2.49 | -1.20 | -2.13 | -1.86 | | 0.108 | 0.017 | 0.567 | 0.015 |
| **TPCM** | 0.26 [0.23, 0.28] | 0.49 | 0.53 | 1.87 | 0.91 | 1.14 | 3.00 | | 1.000 | 1.000 | 1.000 |
| **TPCS** | 0.26 [0.23, 0.29] | 1.05 | 1.09 | 2.43 | 1.47 | 1.69 | 3.54 | 0.56 | | 0.999 | 1.000 |
| **TFI** | 0.25 [0.20, 0.28] | -0.16 | -0.12 | 1.17 | 0.24 | 0.48 | 2.29 | -0.63 | -1.18 | | 0.999 |
| **TFS** | 0.27 [0.25, 0.29] | 1.10 | 1.14 | 2.48 | 1.52 | 1.74 | 3.59 | 0.61 | 0.05 | 1.22 | |
| **Tuned: ratio between extents** | | | | | | | | | | | |
| **TLO** | 0.32 [0.26, 0.35] | | 0.984 | 0.984 | 0.948 | 0.984 | 0.984 | 0.260 | 0.001 | 0.943 | 0.002 |
| **TTOP** | 0.28 [0.26, 0.30] | -1.10 | | 1.000 | 0.328 | 0.532 | 0.587 | 0.009 | $4 \times 10^{-6}$ | 0.295 | $1 \times 10^{-5}$ |
| **TTOA** | 0.28 [0.27, 0.31] | -0.93 | 0.17 | | 0.442 | 0.651 | 0.699 | 0.017 | $9 \times 10^{-6}$ | 0.389 | $3 \times 10^{-5}$ |
| **TPO** | 0.36 [0.30, 0.39] | 1.34 | 2.44 | 2.27 | | 1.000 | 1.000 | 0.968 | 0.115 | 1.000 | 0.195 |
| **TLS** | 0.33 [0.30, 0.36] | 1.07 | 2.16 | 1.99 | -0.25 | | 1.000 | 0.943 | 0.064 | 1.000 | 0.115 |
| **TPCI** | 0.33 [0.28, 0.40] | 1.02 | 2.08 | 1.91 | -0.28 | -0.04 | | 0.943 | 0.067 | 1.000 | 0.117 |
| **TPCM** | 0.35 [0.33, 0.42] | 2.58 | 3.67 | 3.50 | 1.23 | 1.46 | 1.47 | | 0.852 | 0.984 | 0.943 |
| **TPCS** | 0.44 [0.37, 0.47] | 4.26 | 5.36 | 5.19 | 2.92 | 3.12 | 3.10 | 1.69 | | 0.211 | 1.000 |
| **TFI** | 0.35 [0.29, 0.38] | 1.45 | 2.51 | 2.35 | 0.15 | 0.39 | 0.42 | -1.04 | -2.67 | | 0.322 |
| **TFS** | 0.42 [0.37, 0.47] | 4.05 | 5.15 | 4.98 | 2.71 | 2.91 | 2.90 | 1.48 | -0.21 | 2.46 | |
| **Tuned: angulation towards x-axis** | | | | | | | | | | | |
| **TLO** | 47.27 [44.43, 51.94] | | 0.610 | 0.001 | 0.610 | 0.002 | 0.610 | 0.017 | 0.001 | 0.610 | 0.001 |
| **TTOP** | 39.26 [37.82, 40.89] | 4.50 | | 1.000 | 1.000 | 0.025 | 1.000 | 1.000 | 0.017 | 0.610 | 0.163 |
| **TTOA** | 36.27 [35.67, 39.75] | 18.00 | 0.50 | | 1.000 | 0.666 | 1.000 | 1.000 | 0.610 | 0.610 | 0.163 |
| **TPO** | 38.72 [35.36, 42.57] | 4.50 | 0.00 | 0.50 | | 0.216 | 1.000 | 1.000 | 0.163 | 0.610 | 0.163 |
| **TLS** | 32.98 [26.38, 34.97] | 17.05 | 11.63 | 3.89 | 7.24 | | 1.000 | 0.666 | 1.000 | 1.000 | 1.000 |

| | | | | | | | | | | | |
|---|---|---|---|---|---|---|---|---|---|---|---|
| **TPCI** | 33.95 [30.83, 38.43] | 4.83 | 0.54 | 0.54 | 0.54 | 0.32 | | 1.000 | 1.000 | 1.000 | 0.955 |
| **TPCM** | 38.77 [31.88, 41.16] | 12.50 | 0.00 | 0.50 | 0.00 | 3.89 | 0.54 | | 0.610 | 0.610 | 0.610 |
| **TPCS** | 31.57 [29.64, 34.78] | 18.00 | 12.50 | 4.50 | 8.00 | 0.03 | 0.54 | 4.50 | | 1.000 | 1.000 |
| **TFI** | 32.15 [26.58, 46.85] | 4.83 | 4.82 | 4.82 | 4.82 | 0.03 | 0.57 | 4.82 | 0.00 | | 1.000 |
| **TFS** | 31.41 [30.12, 34.36] | 18.00 | 8.00 | 8.00 | 8.00 | 0.28 | 2.14 | 4.50 | 0.00 | 0.12 | |
| **Tuned: exponent on frequency** | | | | | | | | | | | |
| **TLO** | 0.73 [0.69, 0.81] | | 0.996 | 0.945 | 0.132 | 0.105 | 0.005 | $2 \times 10^{-4}$ | $7 \times 10^{-9}$ | 0.002 | $2 \times 10^{-9}$ |
| **TTOP** | 0.68 [0.65, 0.72] | -0.53 | | 0.996 | 0.392 | 0.341 | 0.027 | 0.002 | $2 \times 10^{-7}$ | 0.014 | $5 \times 10^{-8}$ |
| **TTOA** | 0.61 [0.57, 0.70] | -1.15 | -0.61 | | 0.779 | 0.719 | 0.152 | 0.020 | $6 \times 10^{-6}$ | 0.092 | $2 \times 10^{-6}$ |
| **TPO** | 0.52 [0.46, 0.59] | -2.78 | -2.25 | -1.63 | | 0.996 | 0.945 | 0.719 | 0.009 | 0.905 | 0.004 |
| **TLS** | 0.50 [0.40, 0.61] | -2.87 | -2.35 | -1.74 | -0.14 | | 0.956 | 0.779 | 0.018 | 0.945 | 0.009 |
| **TPCI** | 0.43 [0.35, 0.52] | -3.82 | -3.31 | -2.72 | -1.14 | -0.99 | | 0.996 | 0.341 | 0.996 | 0.215 |
| **TPCM** | 0.39 [0.31, 0.47] | -4.55 | -4.02 | -3.41 | -1.77 | -1.61 | -0.57 | | 0.673 | 0.996 | 0.505 |
| **TPCS** | 0.26 [0.20, 0.33] | -6.41 | -5.88 | -5.26 | -3.63 | -3.43 | -2.37 | -1.85 | | 0.450 | 0.996 |
| **TFI** | 0.41 [0.32, 0.48] | -4.03 | -3.52 | -2.93 | -1.35 | -1.19 | -0.20 | 0.37 | 2.16 | | 0.341 |
| **TFS** | 0.25 [0.19, 0.29] | -6.62 | -6.09 | -5.48 | -3.84 | -3.65 | -2.58 | -2.07 | -0.22 | -2.37 | |

**Supplementary Table 8F: Tuned response function parameters change over timing maps.** Like Supplementary Table 8A, but for the areas in the human brain. Therefore here, medians and IQRs are computed over included hemispheres. Data adapted from Harvey et al. (2020)

| **Network** | **Exponent on duration (monotonic)** | | **Exponent on period (monotonic)** | | **Exponent on function (tuned)** | |
|---|---|---|---|---|---|---|
| | **HDS** | ***p*-value** | **HDS** | ***p*-value** | **HDS** | ***p*-value** |
| **Layer-size-matched 1-layer** | | | | | | |
| Layer 1 | 0.08 | $<10^{-7}$ | 0.14 | $<10^{-7}$ | 0.19 | $<10^{-7}$ |
| **Layer-size-matched 5-layer** | | | | | | |
| Layer 1 | 0.12 | $<10^{-7}$ | 0.19 | $<10^{-7}$ | 0.09 | 0.21 |
| Layer 2 | 0.07 | $8 \times 10^{-6}$ | 0.09 | $<10^{-7}$ | 0.13 | $<10^{-7}$ |
| **Layer-size-matched 3-layer** | | | | | | |
| Layer 1 | 0.13 | $<10^{-7}$ | 0.14 | $<10^{-7}$ | 0.18 | $<10^{-7}$ |
| Layer 2 | 0.07 | $6 \times 10^{-6}$ | 0.09 | $<10^{-7}$ | 0.22 | $<10^{-7}$ |
| Layer 3 | 0.10 | $<10^{-7}$ | 0.09 | $6 \times 10^{-7}$ | 0.19 | $<10^{-7}$ |
| **Layer-size-matched 4-layer** | | | | | | |
| Layer 1 | 0.15 | $<10^{-7}$ | 0.16 | $<10^{-7}$ | 0.17 | 0.046 |
| Layer 2 | 0.07 | $9 \times 10^{-6}$ | 0.14 | $<10^{-7}$ | 0.14 | $<10^{-7}$ |
| Layer 3 | 0.10 | $<10^{-7}$ | 0.14 | $<10^{-7}$ | 0.12 | $<10^{-7}$ |
| Layer 4 | 0.12 | $<10^{-7}$ | 0.11 | $2 \times 10^{-5}$ | 0.20 | $<10^{-7}$ |
| **Layer-size-matched 5-layer** | | | | | | |
| Layer 1 | 0.14 | $<10^{-7}$ | 0.11 | $<10^{-7}$ | 0.08 | 0.900 |
| Layer 2 | 0.08 | $6 \times 10^{-6}$ | 0.12 | $<10^{-7}$ | 0.11 | 0.024 |
| Layer 3 | 0.07 | 0.0008 | 0.13 | $<10^{-7}$ | 0.11 | $2 \times 10^{-5}$ |
| Layer 4 | 0.11 | $<10^{-7}$ | 0.16 | $<10^{-7}$ | 0.16 | $<10^{-7}$ |
| Layer 5 | 0.11 | $<10^{-7}$ | 0.10 | $8\times10^{-6}$ | 0.20 | $<10^{-7}$ |
| **Layer-size-matched 5-layer no recurrency** | | | | | | |
| Layer 1 | 0.19 | $<10^{-7}$ | 0.18 | $<10^{-7}$ | No tuned response | No tuned response |
| Layer 2 | 0.22 | $<10^{-7}$ | 0.15 | $<10^{-7}$ | No tuned response | No tuned response |
| Layer 3 | 0.24 | $<10^{-7}$ | 0.17 | $<10^{-7}$ | No tuned response | No tuned response |
| Layer 4 | 0.19 | $<10^{-7}$ | 0.19 | $<10^{-7}$ | No tuned response | No tuned response |
| Layer 5 | 0.15 | $<10^{-7}$ | 0.22 | $<10^{-7}$ | No tuned response | No tuned response |
| **Layer-size-matched 5-layer before training** | | | | | | |
| Layer 1 | 0.14 | $<10^{-7}$ | 0.10 | $<10^{-7}$ | 0.13 | 0.010 |
| Layer 2 | 0.05 | $<10^{-7}$ | 0.11 | $<10^{-7}$ | 0.21 | $<10^{-7}$ |
| Layer 3 | 0.06 | $<10^{-7}$ | 0.10 | $<10^{-7}$ | 0.07 | 0.908 |
| Layer 4 | 0.05 | $<10^{-7}$ | 0.08 | $<10^{-7}$ | 0.07 | 0.908 |
| Layer 5 | 0.04 | $<10^{-7}$ | 0.09 | $<10^{-7}$ | 0.10 | 0.060 |
| **Layer-size-matched 5-layer trained on shuffled data** | | | | | | |
| Layer 1 | 0.17 | $<10^{-7}$ | 0.20 | $<10^{-7}$ | 0.17 | 0.0004 |
| Layer 2 | 0.06 | $4 \times 10^{-5}$ | 0.14 | $<10^{-7}$ | 0.13 | 0.004 |
| Layer 3 | 0.09 | $<10^{-7}$ | 0.12 | $<10^{-7}$ | 0.18 | $<10^{-7}$ |
| Layer 4 | 0.09 | $6 \times 10^{-5}$ | 0.16 | $<10^{-7}$ | 0.17 | $<10^{-7}$ |
| Layer 5 | 0.08 | 0.001 | 0.13 | $<10^{-7}$ | 0.22 | $<10^{-7}$ |

**Supplementary Table 9: Monotonic (duration and period) and some tuned exponents are bimodally distributed.** Given are the test statistics of the Hartigan & Hartigan's dip test for all network layers. These p-values are FDR-corrected. Significant deviations from unimodality underlined

| Response function | Parameter | Initial lower bound | Initial upper bound | Final lower bound | Final upper bound |
|---|---|---|---|---|---|
| Monotonic | $\beta_{duration}$ | 0 | 10 | $1 \times 10^{-8}$ | Inf |
| | expDuration | 0 | 1 | 0 | 1 |
| | $\beta_{period}$ | 0 | 10 | $1 \times 10^{-8}$ | Inf |
| | expPeriod | 0 | 1 | 0 | 1 |
| | Slope | 0.1 | 10 | $1 \times 10^{-8}$ | Inf |
| | Intercept | -2 | 2 | -Inf | Inf |
| Tuned | $\mathrm{State}_{\mathrm{pref}}$ | -0.05 | 1.1 | -0.05 | 1.1 |
| | $\theta$ | 0 | 180 | 0 | 180 |
| | $\mathrm{Period}_{\mathrm{pref}}$ | -0.05 | 1.1 | -0.05 | 1.1 |
| | $\sigma_{\mathrm{min}}$ | 0.001 | 1.5 | 0.001 | 10 |
| | $\sigma_{\mathrm{maj}}$ | 0.001 | 3 | 0.001 | 10 |
| | expFreq | 0 | 1 | 0 | 1 |
| | Slope | 0.1 | 10 | $1 \times 10^{-8}$ | Inf |
| | Intercept | -2 | 2 | -Inf | Inf |

**Supplementary Table 10. Bounds used for scipy's curve_fit function to fit the monotonic and tuned response functions.** Note that the betas for the monotonic components could still become 0 if the corresponding exponents were 0 or 1 for duration or period, respectively.

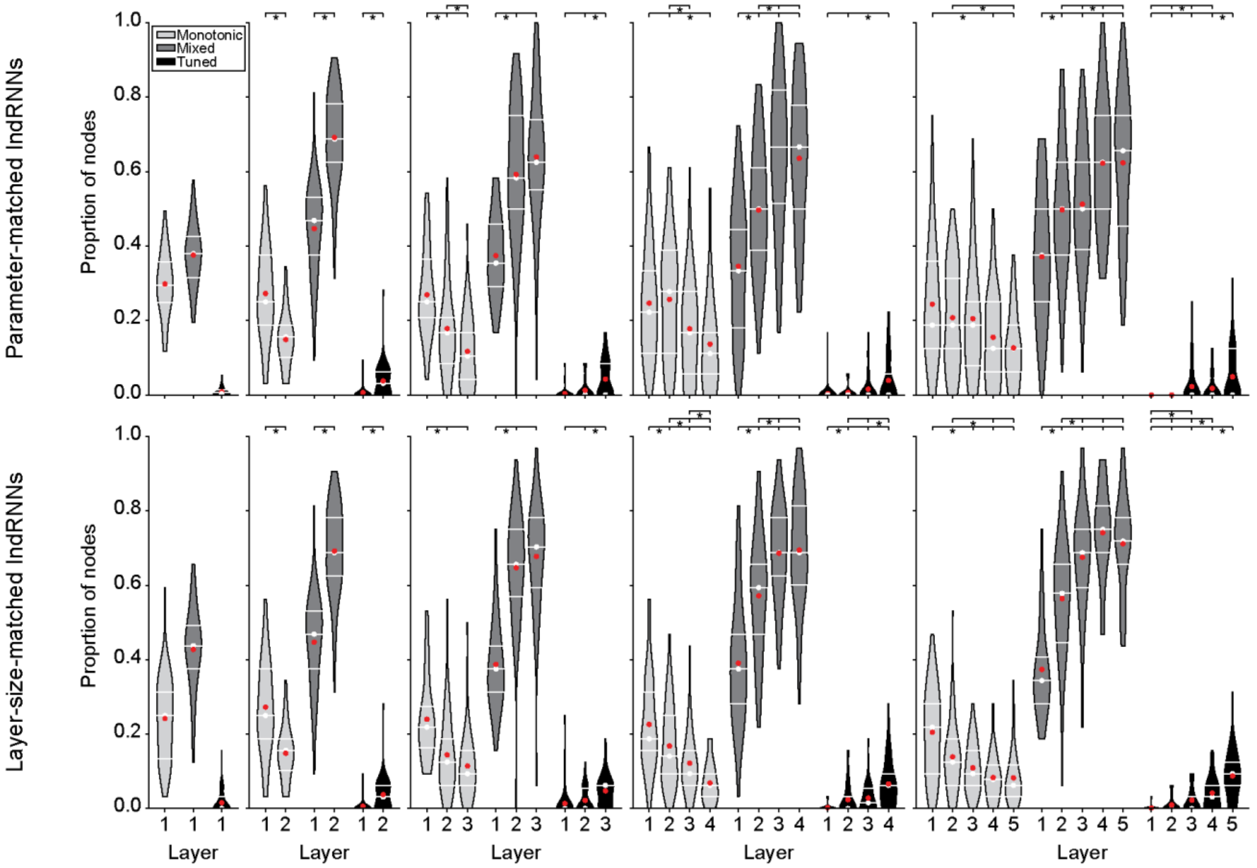


**Supplementary Fig. 6: Progressive transitions from monotonic to mixed and tuned nodes when response functions are allowed to be negative.** Proportions of nodes classified as having monotonic, mixed, or tuned responses in parameter-matched (top) or layer-size-matched (bottom) IndRNNs of different depths. Format follows Fig. 2A.

| | Median [IQR] | Test statistic | | | | |
|---|---|---|---|---|---|---|
| | | **Layer 1** | **Layer 2** | **Layer 3** | **Layer 4** | **Layer 5** |
| | | | **Monotonic** | | | |
| **1-layer network** | | | | | | |
| **Layer 1** | 0.30 [0.25, 0.36] | | | | | |
| **2-layer network** | | | | | | |
| **Layer 1** | 0.25 [0.19, 0.38] | | | | | |
| **Layer 2** | 0.16 [0.10, 0.19] | | | | | |
| **3-layer network** | | | | | | |
| **Layer 1** | 0.25 [0.21, 0.36] | | $3 \times 10^{-4}$ | $4 \times 10^{-9}$ | | |
| **Layer 2** | 0.17 [0.08, 0.21] | -3.83 | | 0.026 | | |
| **Layer 3** | 0.10 [0.04, 0.17] | -6.05 | -2.22 | | | |
| **4-layer network** | | | | | | |
| **Layer 1** | 0.22 [0.11, 0.33] | | 0.757 | 0.078 | 0.001 | |
| **Layer 2** | 0.28 [0.11, 0.39] | 0.31 | | 0.045 | $3 \times 10^{-4}$ | |
| **Layer 3** | 0.17 [0.06, 0.28] | -2.22 | -2.53 | | 0.239 | |
| **Layer 4** | 0.11 [0.06, 0.17] | -3.74 | -4.05 | -1.52 | | |
| **5-layer network** | | | | | | |
| **Layer 1** | 0.19 [0.12, 0.36] | | 0.634 | 0.629 | 0.047 | 0.001 |
| **Layer 2** | 0.19 [0.12, 0.31] | -0.90 | | 0.744 | 0.333 | 0.032 |
| **Layer 3** | 0.19 [0.08, 0.25] | -1.23 | -0.33 | | 0.499 | 0.066 |
| **Layer 4** | 0.12 [0.06, 0.25] | -2.75 | -1.84 | -1.52 | | 0.634 |
| **Layer 5** | 0.12 [0.06, 0.19] | -3.82 | -2.91 | -2.59 | -1.07 | |
| | | | **Mixed** | | | |
| **1-layer network** | | | | | | |
| **Layer 1** | 0.38 [0.31, 0.43] | | | | | |
| **2-layer network** | | | | | | |
| **Layer 1** | 0.47 [0.38, 0.53] | | | | | |
| **Layer 2** | 0.69 [0.62, 0.78] | | | | | |
| **3-layer network** | | | | | | |
| **Layer 1** | 0.35 [0.29, 0.46] | | $6 \times 10^{-9}$ | $4 \times 10^{-12}$ | | |
| **Layer 2** | 0.58 [0.50, 0.75] | 5.93 | | 0.239 | | |
| **Layer 3** | 0.62 [0.55, 0.74] | 7.11 | 1.18 | | | |
| **4-layer network** | | | | | | |
| **Layer 1** | 0.33 [0.18, 0.44] | | 0.004 | $5 \times 10^{-10}$ | $1 \times 10^{-9}$ | |
| **Layer 2** | 0.50 [0.39, 0.61] | 3.16 | | 0.003 | 0.004 | |
| **Layer 3** | 0.67 [0.51, 0.82] | 6.50 | 3.34 | | 0.895 | |
| **Layer 4** | 0.67 [0.50, 0.78] | 6.37 | 3.21 | -0.13 | | |
| **5-layer network** | | | | | | |
| **Layer 1** | 0.38 [0.25, 0.50] | | 0.019 | 0.006 | $2 \times 10^{-8}$ | $1 \times 10^{-8}$ |
| **Layer 2** | 0.50 [0.38, 0.62] | 2.90 | | 0.871 | 0.011 | 0.010 |
| **Layer 3** | 0.50 [0.39, 0.62] | 3.36 | 0.47 | | 0.027 | 0.027 |
| **Layer 4** | 0.62 [0.50, 0.75] | 6.02 | 3.12 | 2.66 | | 0.960 |
| **Layer 5** | 0.66 [0.45, 0.75] | 6.07 | 3.17 | 2.71 | 0.05 | |
| | | | **Tuned** | | | |
| **1-layer network** | | | | | | |
| **Layer 1** | 0.01 [0.00, 0.01] | | | | | |
| **2-layer network** | | | | | | |
| **Layer 1** | 0.00 [0.00, 0.00] | | | | | |
| **Layer 2** | 0.03 [0.00, 0.06] | | | | | |
| **3-layer network** | | | | | | |
| **Layer 1** | 0.00 [0.00, 0.00] | | 0.169 | $1 \times 10^{-6}$ | | |
| **Layer 2** | 0.00 [0.00, 0.00] | 1.38 | | $4 \times 10^{-4}$ | | |
| **Layer 3** | 0.00 [0.00, 0.08] | 5.08 | 3.71 | | | |
| **4-layer network** | | | | | | |
| **Layer 1** | 0.00 [0.00, 0.00] | | 0.416 | 0.133 | $2 \times 10^{-4}$ | |
| **Layer 2** | 0.00 [0.00, 0.00] | 1.19 | | 0.416 | 0.013 | |
| **Layer 3** | 0.00 [0.00, 0.00] | 2.08 | 0.90 | | 0.133 | |
| **Layer 4** | 0.00 [0.00, 0.06] | 4.19 | 3.01 | 2.11 | | |
| **5-layer network** | | | | | | |
| **Layer 1** | 0.00 [0.00, 0.00] | | 1.000 | 0.029 | 0.046 | $4 \times 10^{-7}$ |
| **Layer 2** | 0.00 [0.00, 0.00] | -0.00 | | 0.029 | 0.046 | $4 \times 10^{-7}$ |
| **Layer 3** | 0.00 [0.00, 0.00] | 2.90 | 2.90 | | 0.941 | 0.046 |
| **Layer 4** | 0.00 [0.00, 0.00] | 2.60 | 2.60 | -0.31 | | 0.029 |
| **Layer 5** | 0.00 [0.00, 0.12] | 5.50 | 5.50 | 2.60 | 2.91 | |

**Supplementary Table 11A: Gradual transition from monotonic to mixed and tuned responses over parameter-matched IndRNN layers when response functions are allowed to be negative.** Like Table S6, but with response functions allowing negative slopes.

| | Median [IQR] | Test statistic | | | | |
|---|---|---|---|---|---|---|
| | | Layer 1 | Layer 2 | Layer 3 | Layer 4 | Layer 5 |
| **Monotonic** | | | | | | |
| **1-layer network** | | | | | | |
| **Layer 1** | 0.25 [0.13, 0.31] | | | | | |
| **2-layer network** | | | | | | |
| **Layer 1** | 0.25 [0.19, 0.38] | | | | | |
| **Layer 2** | 0.16 [0.10, 0.19] | | | | | |
| **3-layer network** | | | | | | |
| **Layer 1** | 0.22 [0.16, 0.27] | | $3 \times 10^{-5}$ | $1 \times 10^{-8}$ | | |
| **Layer 2** | 0.12 [0.06, 0.19] | -4.35 | | 0.126 | | |
| **Layer 3** | 0.09 [0.06, 0.16] | -5.88 | -1.53 | | | |
| **4-layer network** | | | | | | |
| **Layer 1** | 0.19 [0.16, 0.31] | | 0.039 | $6 \times 10^{-5}$ | $2 \times 10^{-12}$ | |
| **Layer 2** | 0.14 [0.09, 0.25] | -2.34 | | 0.045 | $4 \times 10^{-6}$ | |
| **Layer 3** | 0.09 [0.06, 0.16] | -4.34 | -2.01 | | 0.009 | |
| **Layer 4** | 0.06 [0.03, 0.09] | -7.30 | -4.96 | -2.95 | | |
| **5-layer network** | | | | | | |
| **Layer 1** | 0.22 [0.09, 0.28] | | 0.065 | 0.001 | $5 \times 10^{-7}$ | $9 \times 10^{-8}$ |
| **Layer 2** | 0.12 [0.06, 0.19] | -2.47 | | 0.370 | 0.018 | 0.007 |
| **Layer 3** | 0.09 [0.06, 0.16] | -3.74 | -1.26 | | 0.244 | 0.163 |
| **Layer 4** | 0.08 [0.03, 0.12] | -5.44 | -2.96 | -1.70 | | 0.750 |
| **Layer 5** | 0.06 [0.03, 0.12] | -5.75 | -3.28 | -2.02 | -0.32 | |
| **Mixed** | | | | | | |
| **1-layer network** | | | | | | |
| **Layer 1** | 0.44 [0.38, 0.49] | | | | | |
| **2-layer network** | | | | | | |
| **Layer 1** | 0.47 [0.38, 0.53] | | | | | |
| **Layer 2** | 0.69 [0.62, 0.78] | | | | | |
| **3-layer network** | | | | | | |
| **Layer 1** | 0.38 [0.31, 0.44] | | $2 \times 10^{-11}$ | $5 \times 10^{-14}$ | | |
| **Layer 2** | 0.66 [0.57, 0.75] | 6.81 | | 0.383 | | |
| **Layer 3** | 0.7 [0.59, 0.78] | 7.68 | 0.87 | | | |
| **4-layer network** | | | | | | |
| **Layer 1** | 0.38 [0.28, 0.47] | | $2 \times 10^{-4}$ | $1 \times 10^{-12}$ | $7 \times 10^{-13}$ | |
| **Layer 2** | 0.59 [0.47, 0.66] | 4.10 | | 0.003 | 0.003 | |
| **Layer 3** | 0.69 [0.62, 0.78] | 7.30 | 3.20 | | 0.904 | |
| **Layer 4** | 0.69 [0.60, 0.81] | 7.42 | 3.32 | 0.12 | | |
| **5-layer network** | | | | | | |
| **Layer 1** | 0.34 [0.28, 0.41] | | $2 \times 10^{-4}$ | $6 \times 10^{-12}$ | $3 \times 10^{-19}$ | $1 \times 10^{-15}$ |
| **Layer 2** | 0.58 [0.45, 0.66] | 4.10 | | 0.008 | $2 \times 10^{-6}$ | $2 \times 10^{-4}$ |
| **Layer 3** | 0.69 [0.59, 0.75] | 7.18 | 3.08 | | 0.118 | 0.460 |
| **Layer 4** | 0.75 [0.69, 0.81] | 9.22 | 5.12 | 2.04 | | 0.460 |
| **Layer 5** | 0.72 [0.66, 0.78] | 8.29 | 4.19 | 1.11 | -0.93 | |
| **Tuned** | | | | | | |
| **1-layer network** | | | | | | |
| **Layer 1** | 0.00 [0.00, 0.03] | | | | | |
| **2-layer network** | | | | | | |
| **Layer 1** | 0.00 [0.00, 0.00] | | | | | |
| **Layer 2** | 0.03 [0.00, 0.06] | | | | | |
| **3-layer network** | | | | | | |
| **Layer 1** | 0.00 [0.00, 0.00] | | 0.051 | $2 \times 10^{-7}$ | | |
| **Layer 2** | 0.00 [0.00, 0.05] | 1.95 | | 0.001 | | |
| **Layer 3** | 0.06 [0.00, 0.06] | 5.41 | 3.46 | | | |
| **4-layer network** | | | | | | |
| **Layer 1** | 0.00 [0.00, 0.00] | | 0.013 | $4 \times 10^{-4}$ | $9 \times 10^{-12}$ | |
| **Layer 2** | 0.00 [0.00, 0.03] | 2.72 | | 0.246 | $7 \times 10^{-5}$ | |
| **Layer 3** | 0.02 [0.00, 0.05] | 3.89 | 1.16 | | 0.004 | |
| **Layer 4** | 0.06 [0.00, 0.09] | 7.08 | 4.35 | 3.19 | | |
| **5-layer network** | | | | | | |
| **Layer 1** | 0.00 [0.00, 0.00] | | 0.172 | 0.009 | $2 \times 10^{-7}$ | $3 \times 10^{-17}$ |
| **Layer 2** | 0.00 [0.00, 0.00] | 1.37 | | 0.169 | $2 \times 10^{-4}$ | $2 \times 10^{-12}$ |
| **Layer 3** | 0.00 [0.00, 0.03] | 3.07 | 1.71 | | 0.043 | $1 \times 10^{-7}$ |
| **Layer 4** | 0.03 [0.00, 0.06] | 5.51 | 4.15 | 2.44 | | 0.007 |
| **Layer 5** | 0.09 [0.06, 0.12] | 8.70 | 7.33 | 5.63 | 3.19 | |

**Supplementary Table 11B: Gradual transition from monotonic to mixed and tuned responses over layer-size-matched IndRNN layers when response functions are allowed to be negative.** Like Table S6, but with response functions allowing negative slopes in layer-size-matched IndRNNs.

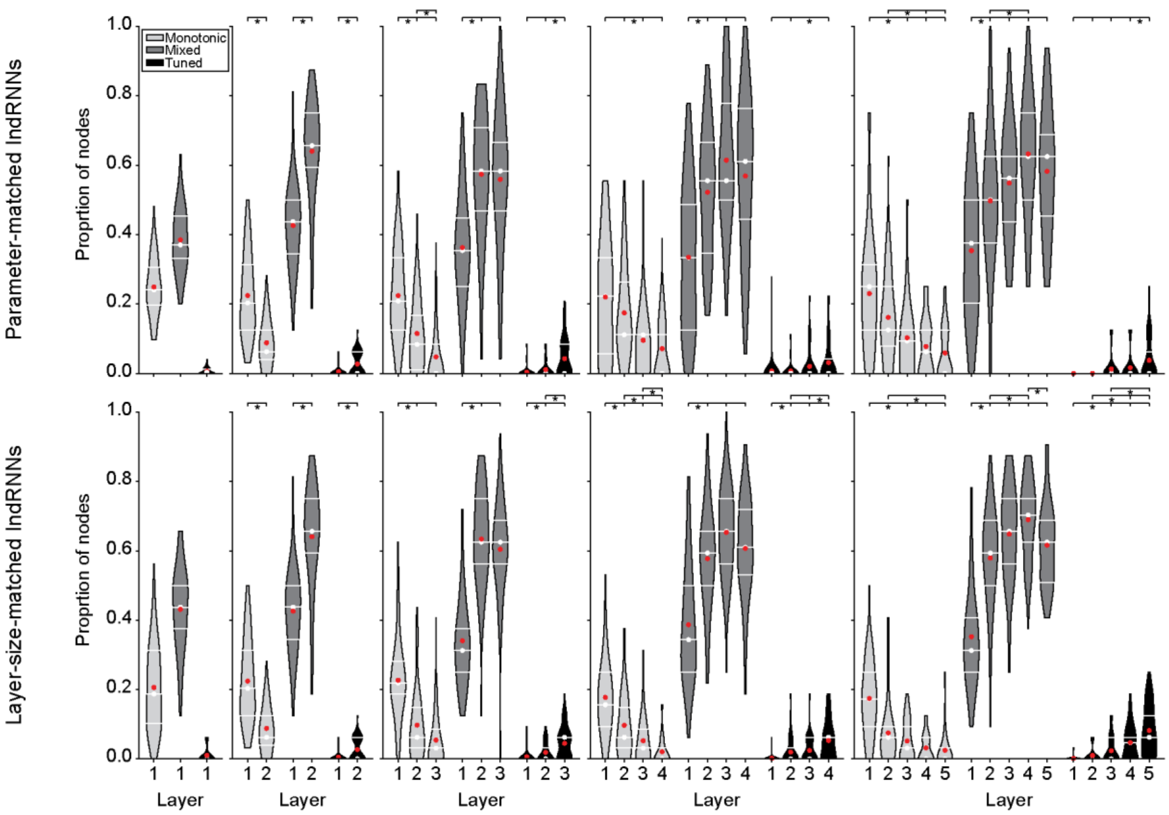


**Supplementary Fig. 6: Progressive transitions from monotonic to mixed and tuned nodes when response function fits are thresholded at 0.8 (instead of 0.2).** Proportions of nodes classified as having monotonic, mixed, or tuned responses in parameter-matched (top) or layer-size-matched (bottom) IndRNNs of different depths. Format follows Fig. 2A.

| | Median [IQR] | Test statistic | | | | |
|---|---|---|---|---|---|---|
| | | Layer 1 | Layer 2 | Layer 3 | Layer 4 | Layer 5 |
| **Monotonic** | | | | | | |
| **1-layer network** | | | | | | |
| **Layer 1** | 0.24 [0.20, 0.31] | | | | | |
| **2-layer network** | | | | | | |
| **Layer 1** | 0.20 [0.12, 0.31] | | | | | |
| **Layer 2** | 0.06 [0.04, 0.12] | | | | | |
| **3-layer network** | | | | | | |
| **Layer 1** | 0.21 [0.12, 0.33] | | $2 \times 10^{-4}$ | $3 \times 10^{-12}$ | | |
| **Layer 2** | 0.08 [0.01, 0.17] | -3.84 | | 0.001 | | |
| **Layer 3** | 0.00 [0.00, 0.08] | -7.14 | -3.31 | | | |
| **4-layer network** | | | | | | |
| **Layer 1** | 0.22 [0.06, 0.33] | . | 0.470 | 0.001 | $7 \times 10^{-6}$ | |
| **Layer 2** | 0.11 [0.11, 0.26] | -0.86 | . | 0.011 | $3 \times 10^{-4}$ | |
| **Layer 3** | 0.11 [0.00, 0.11] | -3.76 | -2.91 | . | 0.470 | |
| **Layer 4** | 0.00 [0.00, 0.11] | -4.86 | -4.01 | -1.10 | | |
| **5-layer network** | | | | | | |
| **Layer 1** | 0.25 [0.12, 0.31] | . | 0.248 | $2 \times 10^{-4}$ | $3 \times 10^{-6}$ | $1 \times 10^{-8}$ |
| **Layer 2** | 0.12 [0.08, 0.25] | -1.82 | . | 0.073 | 0.006 | $2 \times 10^{-4}$ |
| **Layer 3** | 0.09 [0.00, 0.12] | -4.25 | -2.43 | . | 0.581 | 0.248 |
| **Layer 4** | 0.06 [0.00, 0.12] | -5.12 | -3.30 | -0.86 | . | 0.581 |
| **Layer 5** | 0.00 [0.00, 0.12] | -6.05 | -4.23 | -1.79 | -0.93 | |
| **Mixed** | | | | | | |
| **1-layer network** | | | | | | |
| **Layer 1** | 0.37 [0.33, 0.45] | | | | | |
| **2-layer network** | | | | | | |
| **Layer 1** | 0.44 [0.34, 0.50] | | | | | |
| **Layer 2** | 0.66 [0.59, 0.75] | | | | | |
| **3-layer network** | | | | | | |
| **Layer 1** | 0.35 [0.25, 0.45] | | $3 \times 10^{-7}$ | $2 \times 10^{-6}$ | | |
| **Layer 2** | 0.58 [0.47, 0.71] | 5.34 | | 0.683 | | |
| **Layer 3** | 0.58 [0.47, 0.67] | 4.93 | -0.41 | | | |
| **4-layer network** | | | | | | |
| **Layer 1** | 0.33 [0.12, 0.49] | | $4 \times 10^{-4}$ | $6 \times 10^{-8}$ | $3 \times 10^{-6}$ | |
| **Layer 2** | 0.56 [0.35, 0.67] | 3.86 | | 0.171 | 0.467 | |
| **Layer 3** | 0.56 [0.50, 0.78] | 5.74 | 1.88 | | 0.467 | |
| **Layer 4** | 0.61 [0.44, 0.76] | 4.97 | 1.10 | -0.77 | | |
| **5-layer network** | | | | | | |
| **Layer 1** | 0.38 [0.20, 0.50] | | 0.013 | $1 \times 10^{-4}$ | $8 \times 10^{-10}$ | $2 \times 10^{-6}$ |
| **Layer 2** | 0.50 [0.38, 0.62] | 3.07 | | 0.478 | 0.004 | 0.154 |
| **Layer 3** | 0.56 [0.44, 0.62] | 4.36 | 1.29 | | 0.154 | 0.478 |
| **Layer 4** | 0.62 [0.50, 0.75] | 6.50 | 3.42 | 2.13 | | 0.478 |
| **Layer 5** | 0.62 [0.45, 0.69] | 5.20 | 2.13 | 0.84 | -1.30 | |
| **Tuned** | | | | | | |
| **1-layer network** | | | | | | |
| **Layer 1** | 0.00 [0.00, 0.01] | | | | | |
| **2-layer network** | | | | | | |
| **Layer 1** | 0.00 [0.00, 0.00] | | | | | |
| **Layer 2** | 0.00 [0.00, 0.06] | | | | | |
| **3-layer network** | | | | | | |
| **Layer 1** | 0.00 [0.00, 0.00] | | 0.185 | $5 \times 10^{-6}$ | | |
| **Layer 2** | 0.00 [0.00, 0.00] | 1.33 | | 0.001 | | |
| **Layer 3** | 0.00 [0.00, 0.08] | 4.78 | 3.45 | | | |
| **4-layer network** | | | | | | |
| **Layer 1** | 0.00 [0.00, 0.00] | | 0.604 | 0.168 | 0.009 | |
| **Layer 2** | 0.00 [0.00, 0.00] | 0.52 | | 0.357 | 0.041 | |
| **Layer 3** | 0.00 [0.00, 0.00] | 2.01 | 1.49 | | 0.435 | |
| **Layer 4** | 0.00 [0.00, 0.04] | 3.16 | 2.64 | 1.15 | | |
| **5-layer network** | | | | | | |
| **Layer 1** | 0.00 [0.00, 0.00] | . | 1.000 | 0.376 | 0.157 | $2 \times 10^{-6}$ |
| **Layer 2** | 0.00 [0.00, 0.00] | -0.00 | . | 0.376 | 0.157 | $2 \times 10^{-6}$ |
| **Layer 3** | 0.00 [0.00, 0.00] | 1.59 | 1.59 | . | 0.795 | 0.002 |
| **Layer 4** | 0.00 [0.00, 0.00] | 2.20 | 2.20 | 0.60 | . | 0.016 |
| **Layer 5** | 0.00 [0.00, 0.06] | 5.24 | 5.24 | 3.64 | 3.04 | |

**Supplementary Table 12A: Gradual transition from monotonic to mixed and tuned responses over parameter-matched IndRNN layers when response function fits are thresholded at 0.8.** Like Table S6, but with a higher threshold.

| | Median [IQR] | Test statistic | | | | |
|---|---|---|---|---|---|---|
| | | Layer 1 | Layer 2 | Layer 3 | Layer 4 | Layer 5 |
| | | **Monotonic** | | | | |
| **1-layer network** | | | | | | |
| **Layer 1** | 0.19 [0.10, 0.31] | | | | | |
| **2-layer network** | | | | | | |
| **Layer 1** | 0.20 [0.12, 0.31] | | | | | |
| **Layer 2** | 0.06 [0.04, 0.12] | | | | | |
| **3-layer network** | | | | | | |
| **Layer 1** | 0.22 [0.19, 0.28] | | $3 \times 10^{-7}$ | $3 \times 10^{-13}$ | | |
| **Layer 2** | 0.06 [0.03, 0.15] | -5.23 | | 0.027 | | |
| **Layer 3** | 0.03 [0.00, 0.09] | -7.44 | -2.21 | | | |
| **4-layer network** | | | | | | |
| **Layer 1** | 0.16 [0.09, 0.25] | | 0.002 | $1 \times 10^{-8}$ | $1 \times 10^{-15}$ | |
| **Layer 2** | 0.06 [0.03, 0.15] | -3.42 | | 0.021 | $6 \times 10^{-6}$ | |
| **Layer 3** | 0.03 [0.00, 0.09] | -5.99 | -2.56 | | 0.025 | |
| **Layer 4** | 0.00 [0.00, 0.03] | -8.23 | -4.81 | -2.25 | | |
| **5-layer network** | | | | | | |
| **Layer 1** | 0.17 [0.09, 0.25] | | $4 \times 10^{-5}$ | $1 \times 10^{-7}$ | $2 \times 10^{-11}$ | $5 \times 10^{-15}$ |
| **Layer 2** | 0.06 [0.00, 0.09] | -4.55 | | 0.449 | 0.064 | 0.002 |
| **Layer 3** | 0.03 [0.00, 0.09] | -5.68 | -1.13 | | 0.444 | 0.064 |
| **Layer 4** | 0.00 [0.00, 0.06] | -7.03 | -2.48 | -1.35 | | 0.449 |
| **Layer 5** | 0.00 [0.00, 0.03] | -8.11 | -3.56 | -2.43 | -1.08 | |
| | | **Mixed** | | | | |
| **1-layer network** | | | | | | |
| **Layer 1** | 0.44 [0.38, 0.50] | | | | | |
| **2-layer network** | | | | | | |
| **Layer 1** | 0.44 [0.34, 0.50] | | | | | |
| **Layer 2** | 0.66 [0.59, 0.75] | | | | | |
| **3-layer network** | | | | | | |
| **Layer 1** | 0.31 [0.25, 0.38] | | $9 \times 10^{-14}$ | $1 \times 10^{-11}$ | | |
| **Layer 2** | 0.62 [0.56, 0.75] | 7.60 | | 0.480 | | |
| **Layer 3** | 0.62 [0.56, 0.69] | 6.89 | -0.71 | | | |
| **4-layer network** | | | | | | |
| **Layer 1** | 0.34 [0.25, 0.50] | | $5 \times 10^{-6}$ | $7 \times 10^{-12}$ | $4 \times 10^{-8}$ | |
| **Layer 2** | 0.59 [0.50, 0.66] | 4.84 | | 0.068 | 0.358 | |
| **Layer 3** | 0.66 [0.56, 0.75] | 7.11 | 2.27 | | 0.323 | |
| **Layer 4** | 0.61 [0.53, 0.72] | 5.76 | 0.92 | -1.35 | | |
| **5-layer network** | | | | | | |
| **Layer 1** | 0.31 [0.25, 0.41] | | $3 \times 10^{-7}$ | $5 \times 10^{-13}$ | $9 \times 10^{-18}$ | $3 \times 10^{-9}$ |
| **Layer 2** | 0.59 [0.50, 0.69] | 5.47 | | 0.152 | 0.004 | 0.459 |
| **Layer 3** | 0.66 [0.56, 0.75] | 7.53 | 2.05 | | 0.459 | 0.459 |
| **Layer 4** | 0.70 [0.62, 0.75] | 8.85 | 3.38 | 1.33 | | 0.045 |
| **Layer 5** | 0.62 [0.51, 0.69] | 6.24 | 0.77 | -1.28 | -2.61 | |
| | | **Tuned** | | | | |
| **1-layer network** | | | | | | |
| **Layer 1** | 0.00 [0.00, 0.00] | | | | | |
| **2-layer network** | | | | | | |
| **Layer 1** | 0.00 [0.00, 0.00] | | | | | |
| **Layer 2** | 0.00 [0.00, 0.06] | | | | | |
| **3-layer network** | | | | | | |
| **Layer 1** | 0.00 [0.00, 0.00] | | 0.020 | $8 \times 10^{-7}$ | | |
| **Layer 2** | 0.00 [0.00, 0.03] | 2.33 | | 0.010 | | |
| **Layer 3** | 0.06 [0.00, 0.06] | 5.14 | 2.80 | | | |
| **4-layer network** | | | | | | |
| **Layer 1** | 0.00 [0.00, 0.00] | | 0.036 | 0.002 | $1 \times 10^{-8}$ | |
| **Layer 2** | 0.00 [0.00, 0.03] | 2.50 | | 0.306 | 0.002 | |
| **Layer 3** | 0.00 [0.00, 0.06] | 3.53 | 1.02 | | 0.036 | |
| **Layer 4** | 0.06 [0.00, 0.06] | 5.96 | 3.46 | 2.44 | | |
| **5-layer network** | | | | | | |
| **Layer 1** | 0.00 [0.00, 0.00] | . | 0.240 | 0.019 | $8 \times 10^{-7}$ | $4 \times 10^{-13}$ |
| **Layer 2** | 0.00 [0.00, 0.00] | 1.18 | . | 0.164 | $2 \times 10^{-4}$ | $2 \times 10^{-9}$ |
| **Layer 3** | 0.00 [0.00, 0.06] | 2.89 | 1.72 | . | 0.057 | $2 \times 10^{-5}$ |
| **Layer 4** | 0.05 [0.00, 0.06] | 5.34 | 4.16 | 2.44 | . | 0.079 |
| **Layer 5** | 0.06 [0.00, 0.12] | 7.55 | 6.37 | 4.65 | 2.21 | |

**Supplementary Table 12B: Gradual transition from monotonic to mixed and tuned responses over layer-size-matched IndRNN layers when response function fits are thresholded at 0.8.** Like Table S6, but with a higher threshold.in layer-size-matched IndRNNs.
